# “The self” as shifting coalitions of brain systems in a continuous configuration space

**DOI:** 10.64898/2026.11.12.766263

**Authors:** Jeremy I Skipper, George Blackburne, Greg Cooper, Magdalena Jaglinska, Krisztina Jedlovszky, Joanna Kuc, Rosalind McAlpine, Christopher Timmermann

## Abstract

Philosophers and psychologists distinguish many forms of self, yet neuroimaging results often implicate a shared core in the cortical midline or insula. We propose self-related processing arises from shifting coalitions of distributed brain systems, without a core. In a meta-analysis of 912 neuroimaging publications, titles and abstracts clustered into ten forms, including autobiographical memory, body ownership and interoception. No single location was significant in all ten associated brain maps, although each occupied different parts and amounts of midline and insular territory. Their differences were distributed across the brain. Regional patterns varied with study context, corresponded selectively with aspects of consciousness and resembled maps of general functions. Brain patterns combined at continuously varying strengths predicted regions reported in held-out publications better than single patterns or on/off combinations. Self-related processing therefore has no core and arises from shifting coalitions of distributed brain systems whose graded contributions define a continuous configuration space.

## 1 Introduction

Recognising our body as our own, acting on our intentions and remembering our past all seem to belong to the same “I”, which philosophers and psychologists call “the self”. This unity may come from one enduring subject of experience [3], or “self” may be a name for many bodily and mental processes linked by causal relations such as memory and intention [4, 5]. Parfit called the two kinds of views ego theories and bundle theories [5]. The bundle view appears most plainly in the Buddhist chariot dialogue, in which the monk Nāgasena asks whether a chariot is its wheels, axle or frame. No part alone is the chariot, and no further chariot exists alongside the parts, so “chariot” names their arrangement, just as “person” names an arrangement of bodily and mental parts [6]. Because an arrangement can change, selves could share a name without sharing any one part. Psychological accounts sit closer to the bundle view, dividing the self into only a few kinds. For example, James [7] distinguished selves concerned with bodies and possessions, social relationships and mental life, and Gallagher [8] separated a minimal self, as a subject of immediate sensory experience, from a narrative self extended through time. Self-related processing, everything the brain does with information about oneself, takes many more forms than these few kinds. We introduce the term self-form for each family of related activities.

One possible neurobiological counterpart of an enduring subject is a core, a set of locations engaged in every self-form. One candidate territory for a core is the cortical midline, particularly medial prefrontal and posterior cingulate cortex, belonging to the putative default-mode network and repeatedly identified by contrasting self-evaluation and personal memory with control conditions [10–12]. A second candidate territory is the insula, implicated at all three levels of a meta-analytic model by Qin et al. that divides the self into interoceptive (internal bodily states), extero-ceptive (face and body recognition, ownership and agency) and mental (self-related names, memories and attributes) processing [13].

In earlier meta-analyses, however, the midline candidate did not appear in every self-form. Seghezzi et al. [14] compared body ownership, the feeling that a body belongs to oneself, with agency, the feeling of controlling one’s actions. The two shared activation in the left middle insula but otherwise engaged distinct parietal, visual and action-related regions, with no convergence in medial prefrontal or posterior cingulate cortex. Similarly, Qin et al. [13] found that the exteroceptive and mental levels shared anteromedial prefrontal activity, but this overlap did not extend to the interoceptive level. Whether any location is common to every self-form remains open, and a region’s repeated presence across a variety of self-forms still requires an explanation.

Where the midline or insula does recur across self-forms, general functions can explain its presence. In meta-analyses, judgements about one-self and about other people engage overlapping medial prefrontal regions [15, 16]. More broadly, medial prefrontal and posterior midline regions are reported across autobiographical memory, imagining the future, understanding other minds, emotion and models of the current situation [17–20]. Insular regions similarly contribute to bodily and emotional signalling, attention and cognitive control [21, 22]. Tasks involving memory, evaluation or bodily signals can therefore engage the same regions, so finding a region across self-forms need not imply a self-specific function [23, 24].

Regional responses also change with context within a single self-form. Chiao et al. [25], for example, primed bicultural participants with individualistic or collectivistic values. Individualistic priming produced greater medial prefrontal and posterior cingulate responses to general than situational self-judgements, and collectivistic priming reversed that ordering. Recalling a personal memory from an observer’s rather than one’s own visual perspective likewise changes brain responses [26]. Responses can also vary in degree. In the rubber-hand illusion, premotor activity increased with felt ownership of an artificial hand, consistent with the sense of body ownership varying by degree [27].

This variation extends to consciousness, meaning having any experience at all, which has often been treated as inseparable from the self. Descartes took every conscious thought to come with an “I” [3], yet the two may come apart in both directions. Information about oneself can be processed without awareness, as when a participant’s own name, masked so that it could not be consciously identified, still accelerated responses to the names that followed [28]. Conversely, experienced meditators describe continuing awareness with reduced self-identification [29], as can some participants given high doses of psychedelic drugs such as 5-MeO-DMT [30, 31]. Furthermore, it has been proposed that during specific experiences, self-forms are not just reduced but completely absent [32]. Self and consciousness therefore need not have one uniform relationship.

Taken together, no location in the midline, insula or elsewhere has been shown to engage in every self-form, and both candidate territories support general functions beyond self-related processing. Regional patterns also vary with context and by degree, as does their relation to consciousness. We propose that self-related processing arises from shifting coalitions of brain systems, which offers a parsimonious explanation of these observations. A system is a set of brain regions that work together to support a general function, such as retrieving memories, sensing the body or evaluating, and a coalition is the group of systems contributing on a particular occasion. Systems need not be fixed modules, as what a system contributes can depend on its partners [33]. We use the word coalition because its members also work elsewhere and their alliances change. For example, recalling a conversation with embarrassment can strongly involve bodily feeling and evaluation, whereas recalling it to report what was said can place greater weight on memory. We call the pattern of relative contributions a configuration. The word “self” then names systems working together in a configuration, as “chariot” names wheels, axle and frame arranged together.

We formalise configurations as positions in a continuous multidimensional neurobiological configuration space. Each system supplies one axis of the space, and a configuration’s position on that axis is the strength of the system’s contribution. Context would change the strengths, and a familiar self-form would be a set of strengths that recurs often. The proposal thus draws on 1) pattern accounts, which, like bundle theories, describe the self as a set of interacting contributions [34, 35], 2) predictive accounts, in which the sense of one’s own body is inferred from sensory evidence [36, 37], and 3) neural reuse, the principle that the same regions are redeployed across many functions [33].

No single experiment covers all self-forms, so we tested the account in a meta-analysis of 912 neuroimaging publications on self-related processing from NeuroStore, a database of publications and the brain coordinates they report [38]. We estimated the self-forms by grouping publications according to the wording of their titles and abstracts, without their reported coordinates, and mapped the recurring brain locations of each. We ask three questions. The first is whether different self-forms share a core. Our coalition account for the self predicts instead many self-forms with distinguishable regional patterns, which a division into only two or three kinds of self would describe less well. These patterns should also be decentred, with candidate territories engaged to different degrees and no location common to all, and distributed, with differences that persist beyond the regions that best distinguish them.

The second question is where variation in these patterns comes from, and we examine three sources. Contributions within a self-form should change with context, which would not happen if each self-form always engaged one fixed set of regions. Correspondence with consciousness should likewise depend on the particular self-form and aspect of consciousness being compared. The contributing systems should be reused in general activities unrelated to oneself. If instead some systems were dedicated to the self, they would perform functions unique to it.

The third question is how systems combine. We expect graded composition, in which the strength of each system varies continuously, so that configurations can lie anywhere in the configuration space, and we compare it with two more restrictive accounts. In the inventory account, the brain keeps a few complete regional patterns, and each occasion of self-related processing reuses one of them whole, so configurations can sit only at those few points. In the on/off account, systems combine freely, but each contributes at a constant strength or not at all, so configurations can sit only at the corners of the space.

## 2 Results

### 2.1 The corpus and its ten self-forms

We searched neuroimaging titles and abstracts with several sets of self-related terms and found 2,068 candidate records. Three language-model families then judged whether each record investigated self-related processing or used such terms only incidentally, and 1,023 were retained unanimously. We then excluded 108 records with ineligible designs, such as studies of brain structure alone, reviews and case reports. The remaining 915 records, which contained 2,075 analyses and 31,991 reported locations, represented 912 publications (Fig. 1d).

**Fig. 1.**
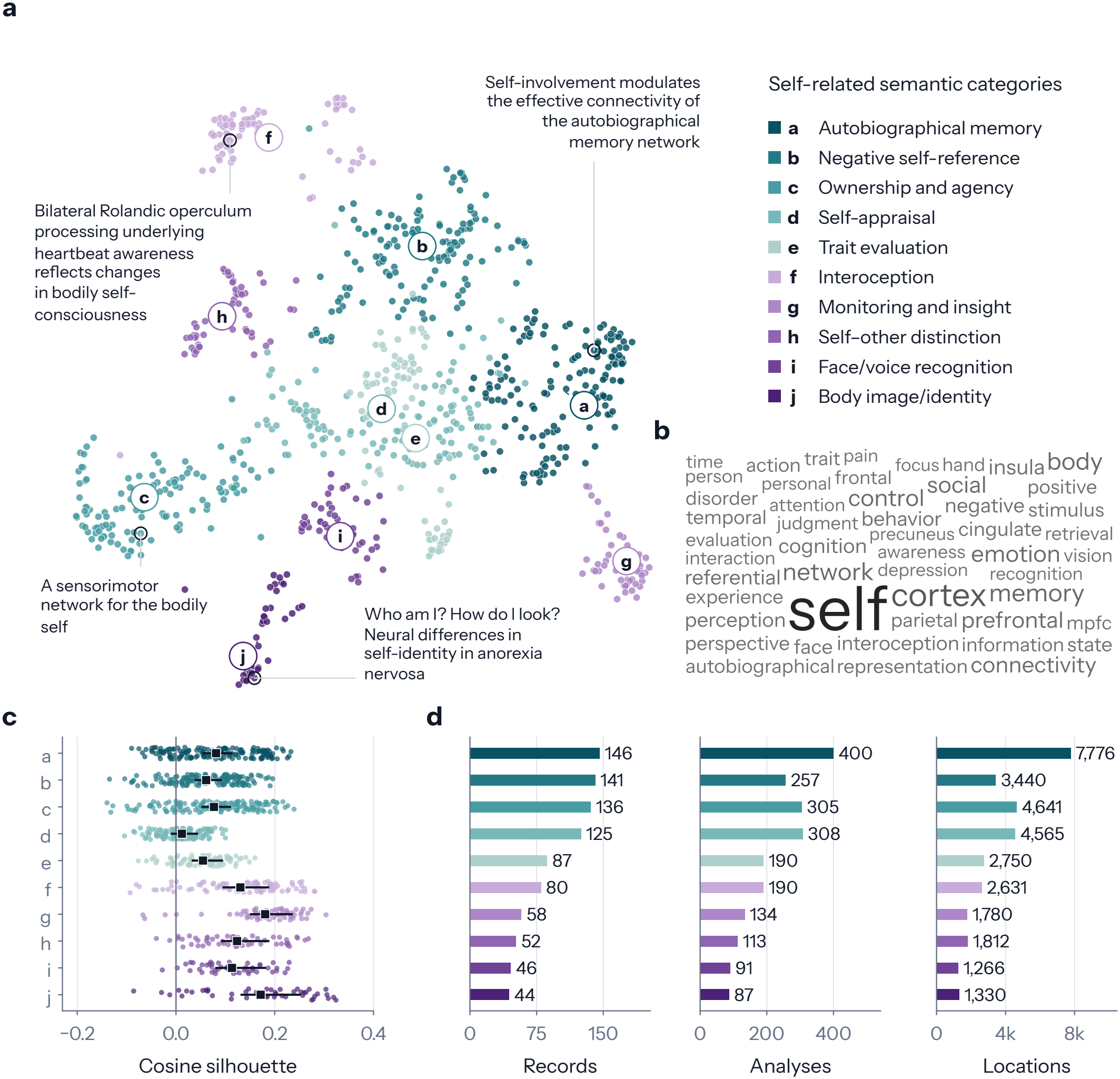
Ten categories of titles and abstracts estimate the self-forms. **a**, Dots show the 915 records, each a title and abstract, arranged by similarity of wording and coloured by category, with four example titles. Lettered circles mark category centres. **b**, The 50 most frequent terms. **c**, Dots show how well each record fits its category. Positive silhouette values mean greater average similarity to a record’s own category, and negative values to another. Squares mark category means and lines their central 95% bootstrap intervals. **d**, Records, analyses and reported locations in each category, before the analyses of a publication were combined.

We grouped the retained titles and abstracts by similarity of wording, without reported coordinates or a preset number of groups. Ten categories occurred most frequently across the settings tried, and language models proposed their names only after membership was fixed (Fig. 1a,b; record counts in Fig. 1d; Supplementary Table 1; Supplementary Method 3). We treat each category as an estimate of a self-form, although wording also reflects methods and populations (Discussion). From here on we use the self-form’s name for its publications and its map. Four concerned remembering or evaluating oneself (Autobiographical memory, Negative self-reference, Self-appraisal and Trait evaluation), and we call these the reflective self-forms. The other six were Ownership and agency, Interoception, Monitoring and insight, Self–other distinction, Face/voice recognition and Body image/identity. In wording, the self-forms were distinct on average but overlapped at their edges. About one fifth of records (18.4%) were closer in wording, on average, to the records of another category than to those of their own (Fig. 1c).

We mapped each self-form with activation likelihood estimation (ALE), testing where its reported locations recurred more often than chance while allowing for uncertainty in coordinates. We also recorded whether each of 910 publications reported a location within or near each of 1,041 atlas regions, which we call its regional profile. In predictive analyses, we controlled for reporting characteristics such as publication year and the numbers of reported locations and analyses (Methods 4.2.4). We repeated most analyses using only coordinates whose standard brain space was known, and the main conclusions held (Supplementary Note 12).

### 2.2 Is there a core?

#### 2.2.1 Self-forms have different maps

We first asked whether the self-forms share a core, beginning with the simplest possibility, that one regional pattern describes them all, whereas the coalition account predicts many distinguishable patterns. The ten maps differed in their regional emphases (Fig. 2a–j). Autobiographical memory, for example, involved superior frontal gyrus, hippocampus and precuneus, whereas Ownership and agency emphasised supramarginal, intraparietal and inferior precentral regions, and Interoception emphasised insula. The midline and insular candidates were each prominent in different maps. Regional profiles differed between self-forms more than when the labels were shuffled (p = 0.0005). The proportion of publications reporting a region differed among self-forms in 82 of the 1,041 regions (7.9%) after correction, and the whole profiles of 25 of the 45 pairs of self-forms differed after Holm correction (Supplementary Fig. 1; Supplementary Table 4). The differences remained in 720 publications explicitly mentioning self (p = 0.00010).

**Fig. 2.**
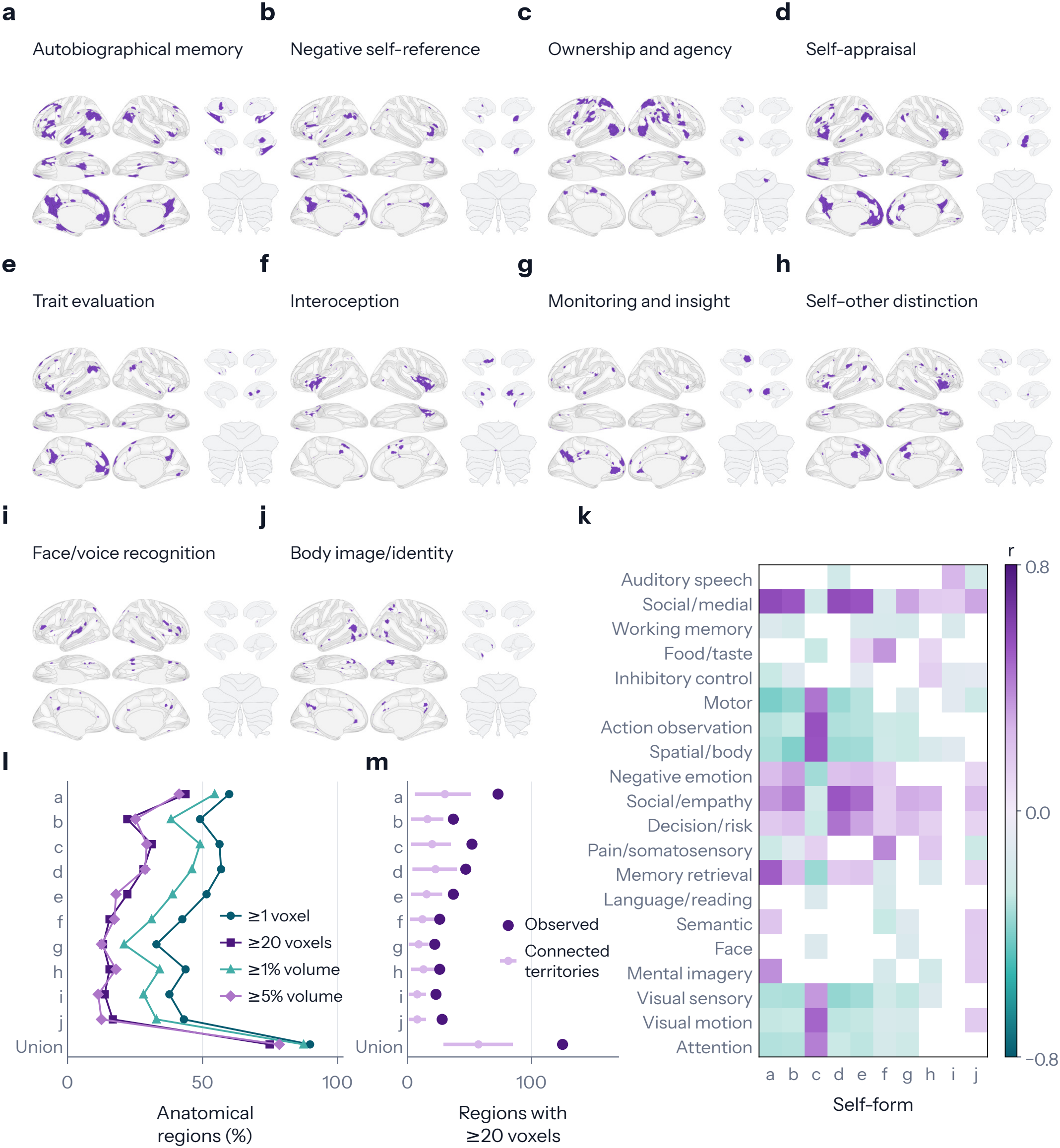
The self-forms have different, widespread maps that resemble maps of general functions. **a–j**, Purple marks where the reported locations of each self-form recur more often than expected by chance (FDR-corrected at *q <* 0.05 across all ten maps). **k**, Each cell gives the correlation of a self-form’s map (columns, lettered as the panels) with a Neurosynth map of a research topic (rows), adjusted for reporting frequency in the Neurosynth source publications. Twenty of 50 topics are shown, chosen by name before the correlations were inspected. Purple and teal mark positive and negative correlations that survived FDR correction against spatially randomised maps, and white cells did not. **l**, Lines give the percentage of 167 anatomical regions that each map covers when a region must hold one significant voxel, 20 voxels, or 1% or 5% of its volume. Union is the ten maps together. **m**, Observed counts of anatomical regions holding at least 20 significant voxels in each map, against the mean and central 95% of counts for 10,000 randomly grown, connected territories of matching atlas-assigned volume.

Earlier syntheses pooled studies across self-forms [9–11], and when self-forms with different maps are pooled, regions reported by the most numerous self-forms can dominate and be mistaken for a common core. To test whether pooling produces such an apparent core, we combined all publications and recomputed ALE. The pooled map occupied 22.1% of the brain mask (Fig. 3a), about seven times the average map of a self-form (3.2%; Supplementary Table 1). Each self-form covered 13.0% of the pooled map on average, while 12.7% of its own territory lay outside it (Fig. 3b). Before thresholding, resemblance to the pooled map was highest for the reflective self-forms (r = 0.74–0.84) and lowest for Ownership and agency, Face/voice recognition and Interoception (r = 0.38–0.46). The pooled map was therefore dominated by the reflective self-forms, which also contributed more than half of the publications.

**Fig. 3.**
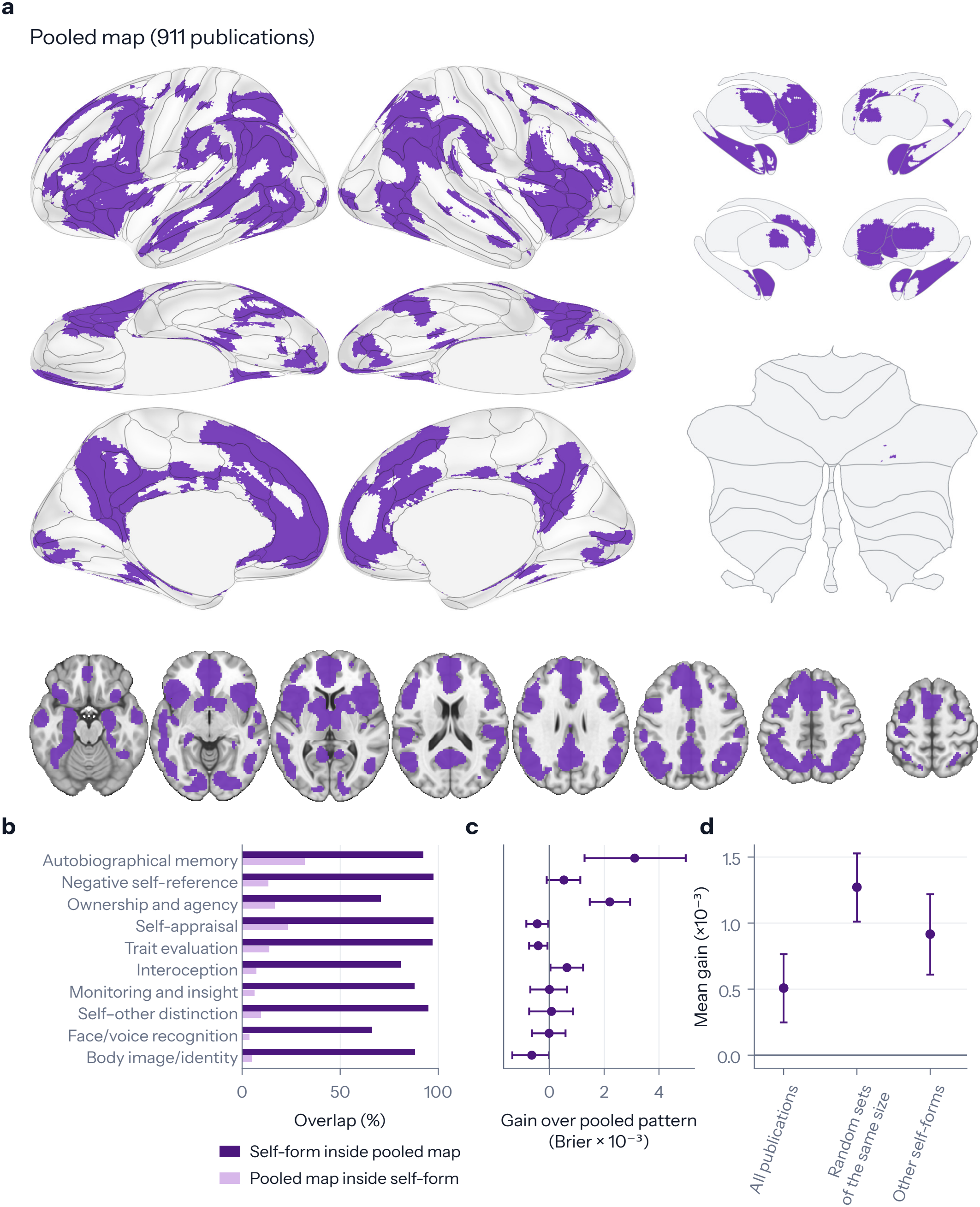
A pooled map covers much of every self-form but loses what distinguishes them. **a**, Purple marks where reported locations recur more often than expected by chance when all 911 publications are pooled. **b**, Overlap of each self-form’s map with the pooled map, in both directions. **c**, Points show the gain in predicting which regions a held-back publication reported when the pattern of its own self-form replaces the pooled pattern. Gain is the reduction in Brier loss, the squared error of predicted probabilities. **d**, Mean gain against each of three pooled patterns, made from all publications, from random sets as large as the self-form, and from the other nine self-forms. Whiskers in c and d are 95% bootstrap intervals over publications. Supplementary Fig. 2a compares every pair of self-forms.

We next asked how much is lost when the regions reported in an unseen publication are predicted from coarser descriptions. Brier loss is the mean squared error between predicted probabilities and observed reporting, and lower values mean better prediction [39]. We express each gain as a percentage of the loss of the model it improves on, and these percentages are small even for reliable gains (Methods 4.2.4). Knowing a publication’s self-form improved prediction over a single pattern pooled across all training publications by 0.55% (reduction 0.000508, 95% interval 0.000248–0.000764, p = 0.0005; Supplementary Note 1). Gains were clearest for Autobiographical memory and for Ownership and agency, and were uncertain for the other self-forms (Fig. 3c,d; Supplementary Table 5). Because existing divisions have two or three kinds of self, we also compared ten groups with three, each learned anew from the text of the training publications rather than taken from a published taxonomy. Ten groups out-performed three by a small but reliable 0.14% after accounting for reporting characteristics and task and clinical indicators (reduction 0.000121, 95% interval 0.000065–0.000177, q = 0.0002). The ten original categories, formed once from the text of the whole corpus, also improved prediction when added to those characteristics and indicators (Supplementary Fig. 2b; Supplementary Tables 6 and 7). Because abstracts can name the regions a study reports, we also regrouped them after filtering anatomical and result statements. The pooling advantage remained, but the ten-versus-three gain was smaller and uncertain with original category weighting (Supplementary Note 1).

#### 2.2.2 No location is common to every self-form

Maps that differ could still share a core. We therefore overlaid the ten thresholded maps and counted how many included each voxel. No voxel was significant in nine or ten maps. Family-wise error correction left no voxel significant in all ten original or filtered maps (Supplementary Table 27). Of 43,414 significant voxels, 27,891 (64%) belonged to one self-form, 4,342 (10%) to at least four, 705 (1.6%) to at least six, and 19 (0.04%) to eight (Fig. 4a–c).

**Fig. 4.**
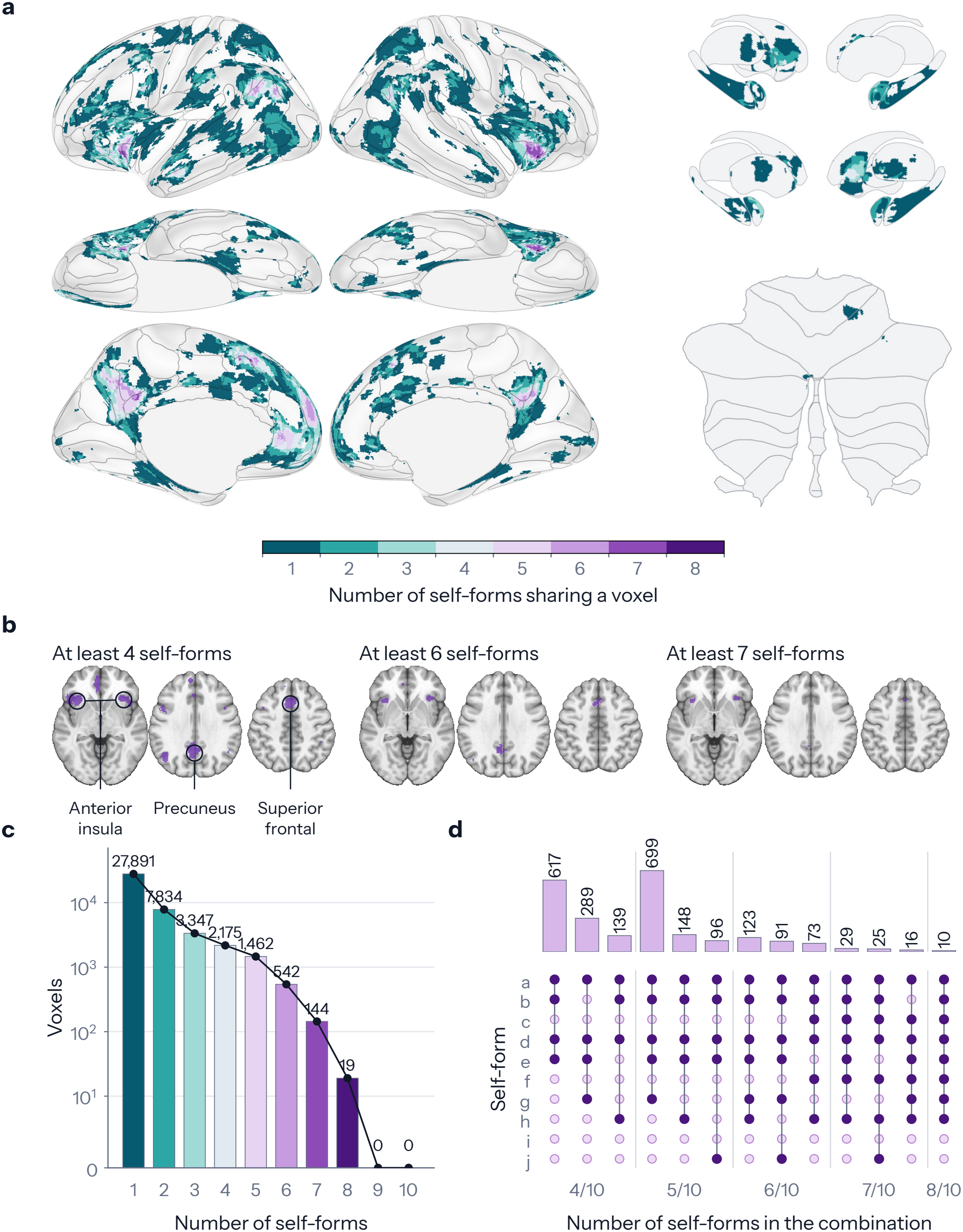
No voxel is shared by all ten self-forms. **a**, Number of self-forms whose corrected maps contain each voxel. **b**, Voxels shared by at least four, six or seven self-forms, at *z* = −4, 32 and 46 mm. **c**, Bars count the voxels shared by exactly each number of self-forms, on a scale that is logarithmic above ten voxels. **d**, Voxels held by particular combinations of exactly four to eight self-forms. For each number, up to three combinations with the most voxels are shown, each holding at least ten. Filled dots identify the self-forms by their letters in Fig. 2.

Sharing was concentrated among the reflective self-forms. Each of the four included 69–97% of the 4,342 voxels shared by at least four self-forms, compared with 8–41% for each of the other six. Even the most widely shared locations did not involve one fixed group of self-forms. The 19 voxels shared by eight maps formed five combinations (Fig. 4d; Supplementary Note 2). Each combination lacked a different pair of self-forms, with Face/voice recognition absent most often.

Territory shared by at least four of the ten self-forms included medial frontal, posterior cingulate/precuneus, anterior insular and lateral temporal regions (Fig. 4b), which overlap the usual midline and insular candidates for a core. We therefore asked whether every map reached the candidate territories, taking medial prefrontal cortex and posterior cingulate/precuneus for the midline candidate and anterior insula for the insular candidate. Every map contained significant voxels in all three territories, but to very different extents, from 10 voxels (Body image/identity) to 994 (Autobiographical memory) in medial prefrontal cortex and from 33 (Body image/identity) to 841 (Interoception) in anterior insula. Eight of 167 anatomical regions and two of the 1,041 finer regions contained voxels from all ten maps, although at different locations, and requiring at least 20 significant voxels from each map reduced these counts to one and none (Supplementary Note 2).

#### 2.2.3 Differences between self-forms are distributed across the brain

One region or division could still carry what distinguishes the self-forms. If so, it should predict a publication’s self-form as well as the whole brain does. We looked first at how widely the maps extended. Because larger maps reach more regions, we compared each with randomly grown territories of equal volume, and every map reached more of 167 anatomical regions, counting those with at least 20 significant voxels, than these references (Holm-adjusted p = 0.0022–0.0264; Fig. 2l,m). Across 19 atlas divisions, each map placed more than 1% of its significant atlas-assigned voxels in 42–68% of divisions (Fig. 5b).

**Fig. 5.**
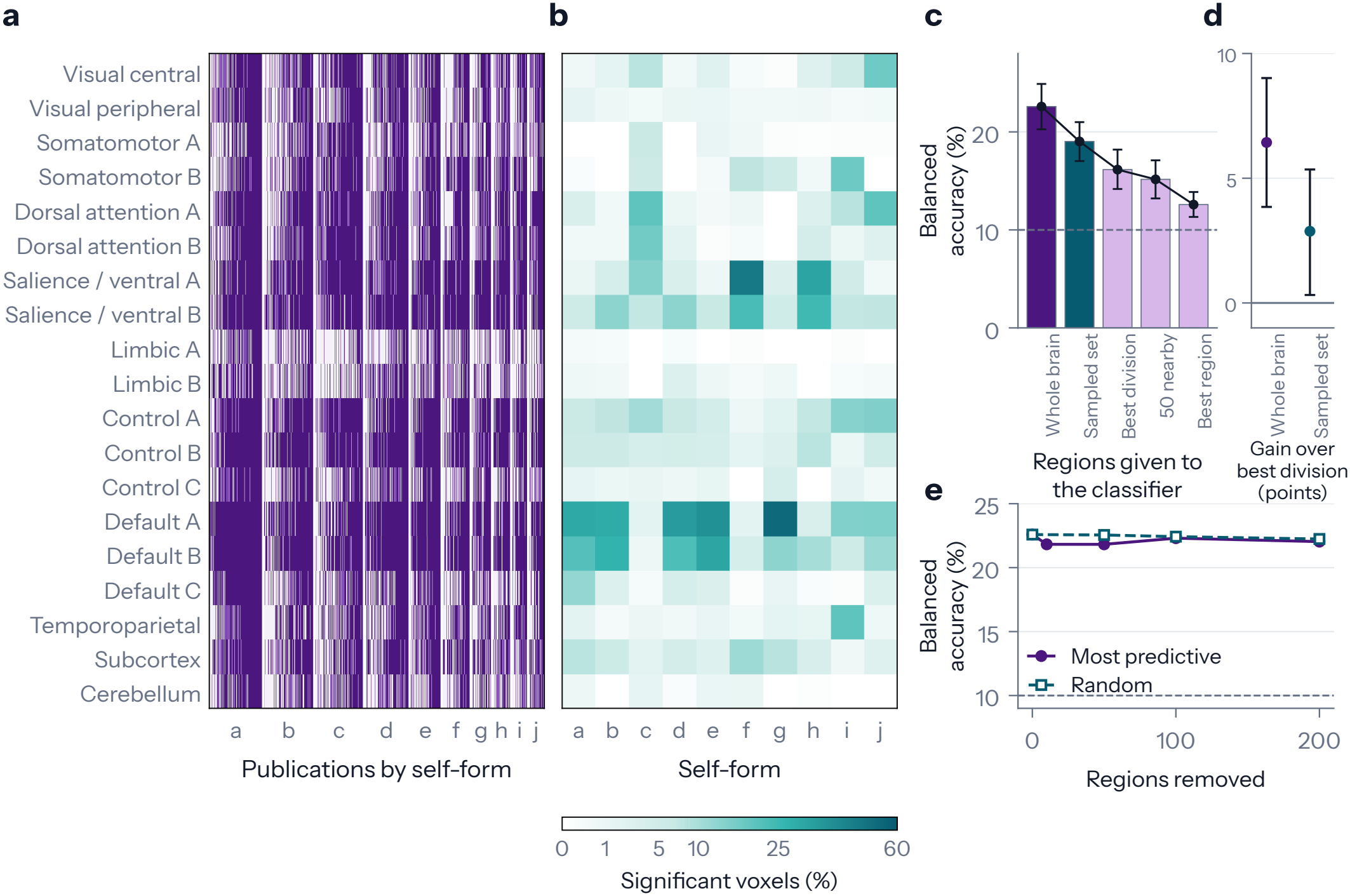
Differences between self-forms are distributed across the brain. **a**, Divisions reported by each of 910 publications, grouped by self-form (letters as in Fig. 2). Divisions are 17 putative cortical resting-state networks, the subcortex and the cerebellum, and colour marks a division in which a publication reported any region. **b**, Percentage of each map’s significant atlas-assigned voxels in each division. **c**, Balanced accuracy of a classifier that predicts each publication’s self-form from its reported regions after training on other publications. Balanced accuracy averages the percentage classified correctly within each self-form. The sampled set draws as many regions as the best division from across the brain, and every choice of regions used the training publications only. **d**, Gain of the whole brain and of the sampled set over the best division, in percentage points. **e**, Balanced accuracy after up to 200 of the most predictive regions, or as many random regions, were removed and the classifier retrained. Dashed lines mark chance (10%), and whiskers are 95% bootstrap intervals over publications.

To test whether this spread helps distinguish self-forms, we trained a logistic-regression classifier to predict an unseen publication’s self-form from its reported regions. With the whole brain, balanced accuracy, which gives self-forms equal weight, reached 22.6%, more than twice the 10% chance level (permutation p = 0.001). Restricting the classifier to the most predictive region, the most predictive atlas division or 50 nearby regions gave 12.6%, 16.2% and 15.2%, respectively. The best region and the best division always fell within the putative default-mode network, which contains the midline core candidate, yet the whole brain outperformed the best division by 6.4 percentage points (95% interval 3.8–9.0, q = 0.0002). To distinguish the value of spatial spread from having more predictors, we used a set of the same size sampled uniformly from all 1,041 regions. It reached 19.0%, a gain of 2.9 points (95% interval 0.3–5.4; Fig. 5c,d), so scattered regions predicted better than the same number concentrated in the best division.

Information that distinguishes the self-forms also remained after the strongest predictors were removed. Removing the 200 most predictive regions and retraining left accuracy at 22.0%, close to the 22.2% that followed removal of 200 random regions (difference -0.2 percentage points, 95% interval -1.7 to 1.3, q = 0.865; Fig. 5e; Supplementary Tables 9 and 10).

### 2.3 Where does the variation come from?

#### 2.3.1 Context within a self-form

We next asked whether context explains part of the variation within a self-form. Publications within a self-form reported different regions (Fig. 5a; Supplementary Note 4). If each self-form had one fixed set of regions, the circumstances of a study, which titles and abstracts convey through tasks, stimuli and participants, should not predict this variation.

Differences between descriptions within a self-form, obtained by subtracting each category’s average numerical summary of wording from those of its publications (Methods 4.4.1), added small but reliable predictive gains of 0.13% beyond category and reporting characteristics (reduction 0.000114, 95% interval 0.000049–0.000181, q = 0.0008) and 0.06% beyond task and clinical indicators (0.000056, q = 0.0002; Supplementary Fig. 2c; Supplementary Table 11). Because abstracts can state the brain results being predicted, we omitted titles and filtered anatomical and result statements, and a gain remained in 867 publications (5.73 × 10^−5^, 95% interval 3.67 × 10^−5^–7.80 × 10^−5^, Holm-adjusted p = 0.0003; Supplementary Table 12).

#### 2.3.2 The aspect of consciousness

We next asked whether correspondence with consciousness depends on which self-form is paired with which aspect of consciousness. The aspects are seven groupings of consciousness research from our companion meta-analysis [40], and each aspect has a map of the proportion of publications reporting each region, as does each self-form. After removing 113 publications that appeared in both literatures, leaving 799 self-only and 465 consciousness-only publications, we correlated the maps across regions. One uniform relationship would give every self-form a similar profile of correlations across the aspects. Instead, correspondence was strong but selective. Autobiographical memory matched Awareness in memory and learning more closely than Conscious perception and attention (r = 0.636 versus 0.363), whereas Ownership and agency showed the reverse (0.398 versus 0.666; Fig. 6a,b). The strongest of the 70 pairings linked Interoception to Interoceptive awareness and pain experience (r = 0.691).

**Fig. 6.**
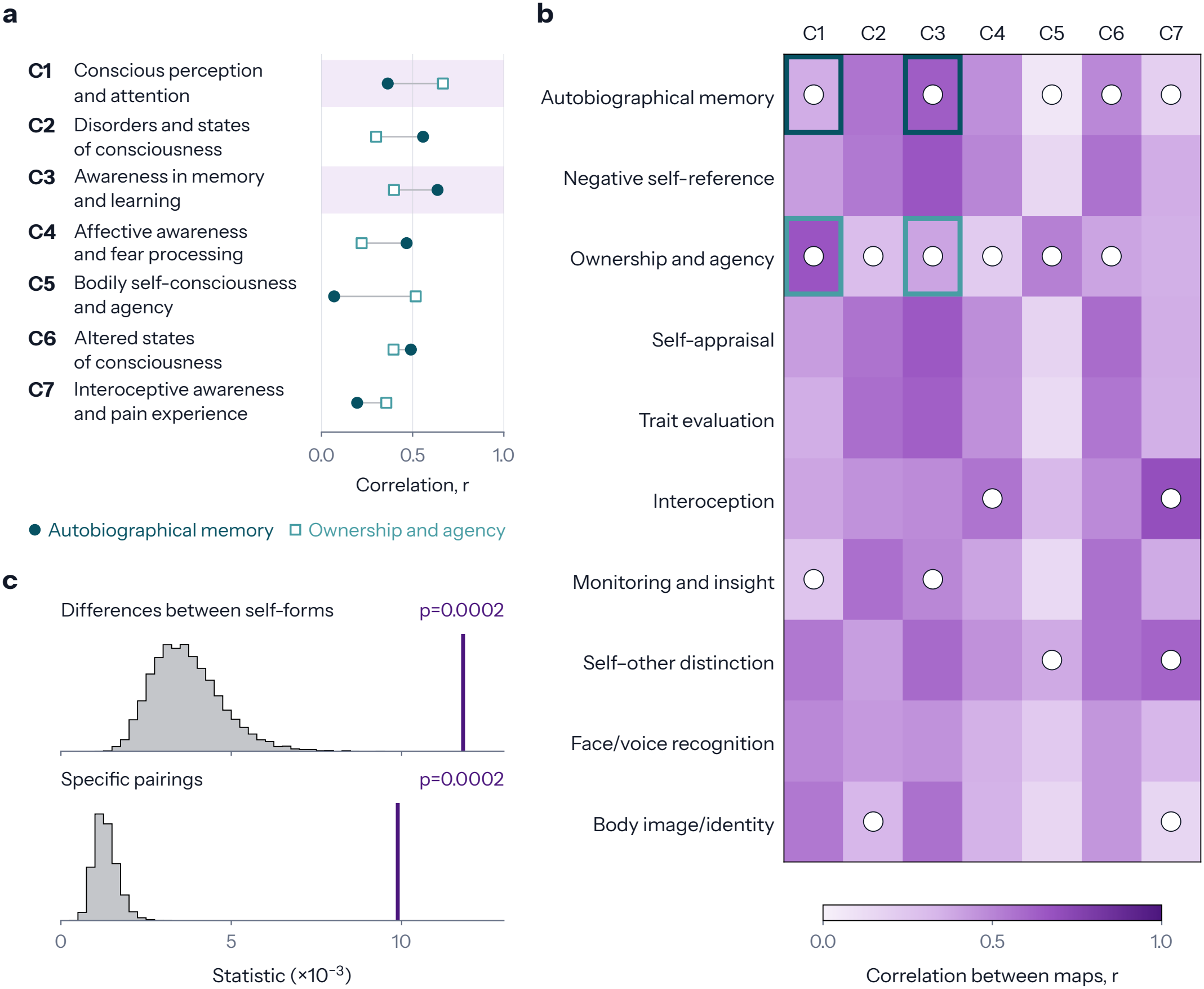
Correspondence with consciousness depends on the self-form and the aspect of consciousness. Maps give the proportion of 799 self-only and 465 consciousness-only publications that reported each region. **a**, Correlations of two illustrative self-forms with seven groupings of consciousness research, C1–C7 [40]. Shading marks C1 and C3, between which the ordering of the two self-forms reverses. **b**, Cells give the correlation between the map of each self-form and each grouping. Circles mark the 19 correlations that departed from their values when the self-form labels were shuffled (FDR-corrected across 70 cells), and outlines mark the four shaded in a. **c**, Purple lines show two observed statistics against their distributions under 9,999 shuffles of the labels (grey). The upper measures differences between the self-forms’ profiles of correlation across the groupings. The lower measures differences specific to pairings, after allowing for the average correlation of each self-form and each grouping.

Such selectivity is what the coalition account expects, so we assessed the specificity of this ten-by-seven pattern of pairings with two tests. The self-forms’ profiles of seven correlations differed more than expected after label shuffling (Holm-adjusted p = 0.0002), and the pattern remained after subtracting each row’s and column’s average, so it did not arise from some maps resembling all others more strongly (Holm-adjusted p = 0.0002; Fig. 6b,c; Supplementary Fig. 3). Correspondence was therefore strong for some pairings and weak for others (r = 0.07–0.69), so no single uniform relationship was found.

#### 2.3.3 Systems shared with other functions

Variation should also follow the general systems on which each self-form draws. We compared the ten maps with 50 Neurosynth topic maps, each summarising the imaging literature on one topic, after excluding identifiable publications from the self-corpus (Fig. 2k). Every self-form matched general functions, and 310 of 500 correlations adjusted for common reporting density survived correction against spatially randomised maps.

Each self-form’s strongest match was a topic of related content. The strongest match for Negative self-reference, Self-appraisal, Trait evaluation and Body image/identity was the medial prefrontal social topic. Autobiographical memory (r = 0.606) and Monitoring and insight matched the resting-state topic most strongly, and Ownership and agency matched action observation most strongly (0.541). For Face/voice recognition, Interoception and Self–other distinction, the strongest matches were auditory speech, the anterior cingulate topic and the social/empathy/moral topic, respectively. These matches survived correction, as did those of Autobiographical memory with memory retrieval/encoding (0.508) and of Interoception with pain/somatosensation (0.392; all q = 0.00029).

Neural reuse also predicts that regions reported together in self-related studies follow the organisation seen across imaging studies in general. To compare that organisation across literatures, we took the correlations of reporting between the 1,041 regions and derived gradients, continuous axes on which regions with similar links to the rest of the network lie close together (Methods 4.4.3). Each of the first ten was matched with its strongest counterpart among gradients from general-imaging publications matched on year and coordinate count. Absolute correlations ranged from 0.342 to 0.818, and the first seven passed both corrected tests against spatially randomised gradients, as reuse predicts, although later gradients were less reproducible across split samples (Fig. 7a,c; Supplementary Fig. 4). With each publication placed at the average position of its reported regions, self-forms differed on seven of the ten gradients (Fig. 7b). Membership of a self-form explained 1.4–21.3% of positional variance, leaving most variation within self-forms along continuous axes (Supplementary Fig. 5a–j; Supplementary Table 13).

**Fig. 7.**
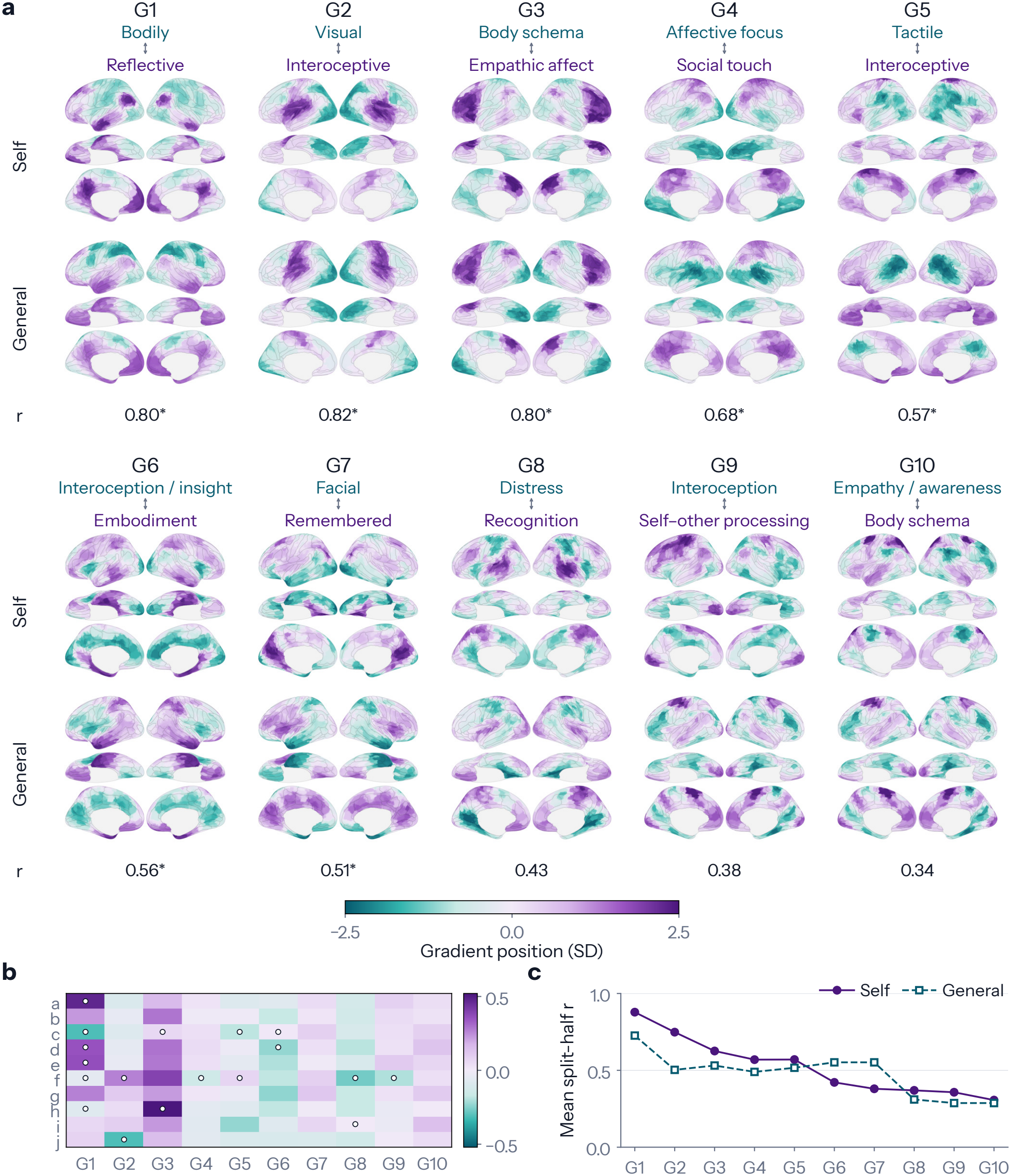
Gradients of the self literature match those of imaging research in general. **a**, The first ten gradients from self publications (Self) appear above their strongest matches among 25 gradients from imaging publications that our search did not retrieve (General), with each pair’s correlation, *r*. A gradient orders regions by how similarly they are reported across publications, and labels summarise the research at its two ends. Asterisks mark matches that passed both corrected tests against spatially randomised gradients. **b**, Mean position of each self-form’s publications on each gradient (letters as in Fig. 2). Circles mark means that departed from their values when the self-form labels were shuffled (FDR-corrected across 100 cells). **c**, Reliability of each gradient and of its match, the mean correlation between matched gradients from the two halves of the publications across 25 random splits.

### 2.4 How do systems combine?

The third question was whether the contributions of systems vary continuously in strength, as we propose, or in one of two discrete ways. We fitted a model for each account (Fig. 8a). The inventory model learned a fixed set of complete regional patterns and assigned each publication one of them. The other two models built each publication’s pattern from components. A component is a fixed spatial pattern, learned from the publications, that raises the probability of reporting some regions and lowers it for others, and it stands in for a system. A component’s strength is how much it contributes to a publication’s pattern. In the on/off model each component was present at a constant strength or absent, as a fixed module would be, and in the graded model its strength could take any value, as in a coalition. A publication’s strengths defined its configuration, and each component’s strength supplied an axis of the fitted configuration space.

**Fig. 8.**
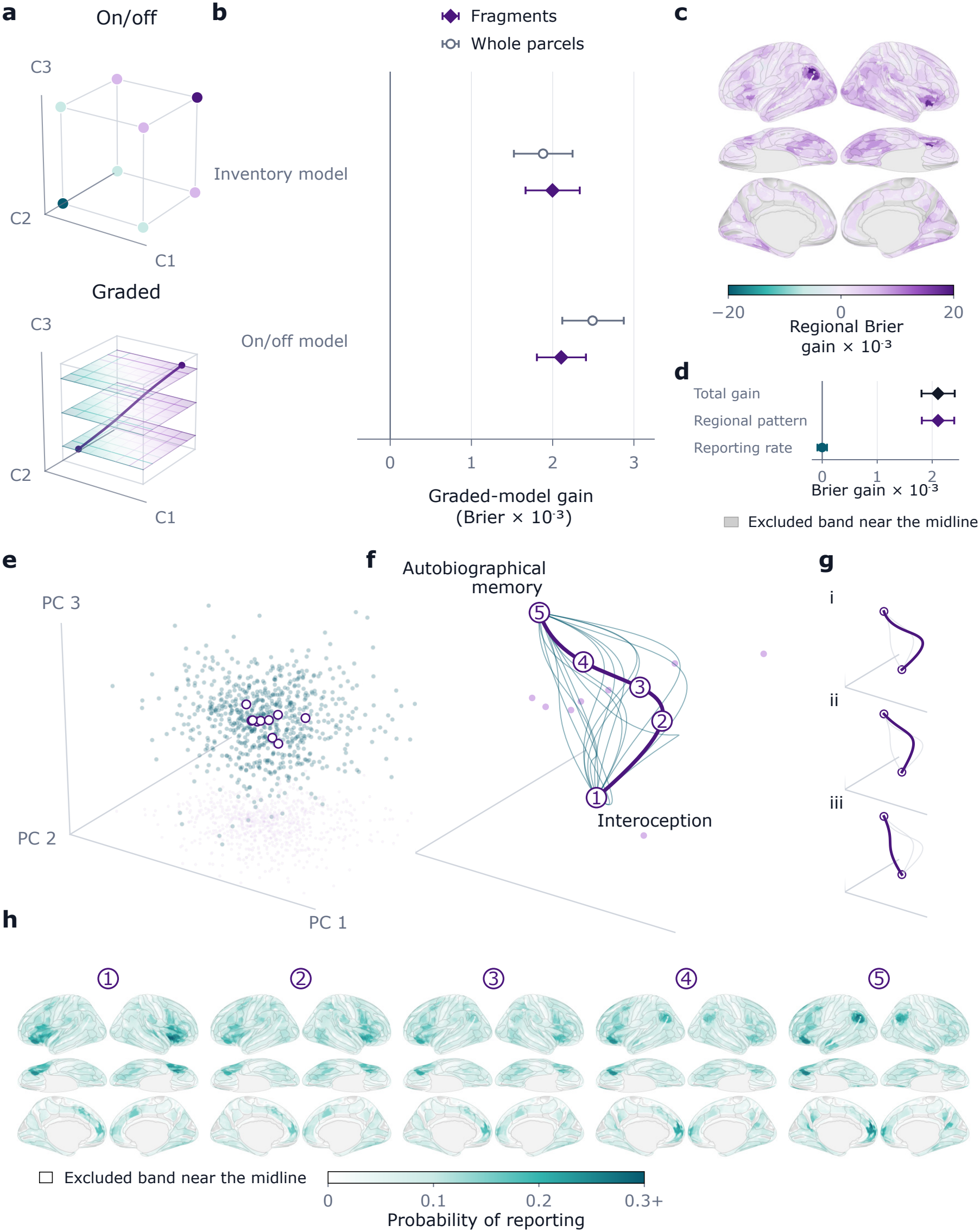
Graded combinations predict unseen regional patterns better than fixed patterns or on/off combinations. **a**, Three components, each a fixed spatial pattern, give eight combinations when each is present or absent (top) and a continuous space when their strengths vary (bottom). The line illustrates a path through that space. **b**, Points show how much the graded model reduces Brier loss relative to the inventory model of fixed patterns and to the on/off model, in 910 held-back publications. One hemisphere predicted the other, using the parts of cortical parcels away from the midline (fragments) or only parcels wholly away from it (whole parcels). **c**, The same gain over the on/off model for each fragment. Purple favours the graded model and teal the on/off model, and grey marks the excluded midline cortex and background. **d**, Gain over the on/off model, separated into regional pattern (which regions were reported) and reporting rate (how much was reported). **e**, Dots are the training publications of one graded model on three axes (PC 1–PC 3) that summarise their ten component strengths. Circles mark the means of the self-forms. **f**, Paths from the mean of Interoception to that of Autobiographical memory. The thick path carries the five numbered points mapped in h, and thin lines are other such paths. Pale dots mark self-form means. **g**, Three such paths, each bending towards a different pair of other self-forms (Supplementary Note 11). Path i is the thick path in f. **h**, Predicted probability of reporting each fragment at the five numbered points. The midline band is excluded. The paths are constructed illustrations. Whiskers in b and d are 95% bootstrap intervals over publications.

We fitted all three models to the cortical regions reported by the same 910 publications, with their reporting characteristics, and none was given the categories or gradients (Methods 4.5.2). We then gave each model the regions that a heldout publication reported in one hemisphere and scored how well it predicted those reported in the other (Fig. 8b).

Graded combinations outperformed every tested version of the inventory model, including a training-selected choice of one to 20 patterns. Brier loss fell by 2.5% (reduction 0.001998, 95% interval 0.001666–0.002334, q = 0.0001; Supplementary Tables 14 and 15). Graded combinations also outperformed the on/off model, by 2.6% (0.002105, 95% interval 0.001806–0.002410, Holm-adjusted p = 0.0004). The advantage held with more on/off components, and both comparisons held for log loss (Supplementary Table 16). Almost all the gain over the on/off model concerned which regions were reported, with little contribution from total reporting rate (Fig. 8c,d), and both advantages held across folds, directions, restricted subsets and stricter whole-parcel exclusion (Supplementary Note 10).

The best graded model had the maximum of ten components, so each configuration was a set of ten continuously varying strengths. The centres of the self-forms occupied different positions in this space (Fig. 8e). To illustrate it, we constructed several paths between two centres, which passed through blended regional patterns that varied with the route (Fig. 8f–h).

## 3 Discussion

Here, we systematically examined the brain organisation of ten forms of self-related processing, investigating their shared and distinct regional patterns, their relationship to general functions and consciousness, their dependence on context, and whether they are better explained by a common core or changing configurations of distributed brain systems.

We grouped titles and abstracts into ten self-forms whose maps emphasised different regions (Figs. 1 and 2), and knowing a publication’s self-form improved prediction of the regions it reported, with ten groups outperforming three (Fig. 3c,d). Pooling all publications instead produced a large map dominated by the reflective self-forms (Fig. 3a,b). No location was common to every self-form, although every map reached midline and insular territory at different locations and to very different degrees (Fig. 4). The differences were distributed across the whole brain, which predicted a publication’s self-form better than the best region or division did, and prediction survived removal of the 200 most predictive regions (Fig. 5).

Within a self-form, study descriptions predicted part of the variation in reported regions (Supplementary Fig. 2). Correspondence with consciousness was strong for some pairings of self-form and aspect of consciousness and weak for others (Fig. 6). Every self-form matched maps of general functions studied without reference to the self, and regions reported together followed gradients shared with imaging research in general (Figs. 2k, 7 and 7).

Graded combinations of components predicted the regions reported by held-out publications better than either the inventory model or the on/off model, and the fitted strengths defined a continuous configuration space in which the self-forms occupied different positions (Fig. 8). On the coalition account, what distinguishes self-related processing is the configuration through which shared systems relate information to one’s body, actions or history. Overall, these findings support a coalition account of the self, in which self-related processing emerges from dynamic, graded configurations of shared brain systems rather than a fixed, self-specific core.

The case against a core rests on these results together rather than on any one of them. A core would have to 1) appear in every self-form, 2) do work specific to the self and 3) carry what distinguishes the self-forms. No location met the first requirement in even nine of the ten maps, and the most widely shared locations involved changing combinations of self-forms. Recurring midline and insular territory failed the second, as neural reuse predicts and earlier critics argued [23, 24, 33], because every self-form’s map resembled maps of topics studied without reference to the self. The third failed because prediction survived removal of the most informative regions, so no small set of locations carried the distinctions. Even a few voxels significant in all ten maps would therefore not restore a core.

Existing divisions separate minimal from narrative selves [8] or interoceptive, exteroceptive and mental levels [13]. We grouped the literature by its wording instead of testing these divisions, and some self-forms resemble them, as Interoception resembles the interoceptive level. Ten is where the clustering settled, not a count of independent selves, because the self-forms overlapped at their edges (Fig. 1c). There are therefore many distinguishable configurations, more than two or three kinds of self would describe, and because configurations vary continuously, there may be no fixed number of selves to count.

The midline core of earlier syntheses mostly describes the reflective self-forms. Northoff et al. reported midline recurrence across self-related tasks, and separating the self-forms explains why. Our pooled map resembled the reflective self-forms most and Ownership and agency least, and the locations shared across maps belonged mostly to the same four reflective self-forms (Figs. 3, 4). Read as a shared core, a pooled map would therefore present regions for remembering and evaluating oneself as regions common to the self, which is how averaging across self-forms can mislead. Earlier meta-analyses of narrower comparisons fit this reading, with partial overlap alongside distinct regions [13–16, 41, 42].

Reported activity cannot establish which regions are necessary, because that requires testing whether an ability survives damage. A patient with extensive bilateral damage involving insula, anterior cingulate and medial prefrontal cortex retained several abilities involving self-awareness [43], yet in a patient group focal medial prefrontal damage abolished the self-reference memory advantage [44]. Which regions are necessary may therefore depend on the ability tested.

Context changes which systems contribute within a self-form, and how strongly. Within one study, for example, cultural priming reversed the ordering of midline responses [25]. Across the literature, study descriptions predicted part of the variation within a self-form, a small effect consistent with such dependence (Supplementary Fig. 2). When two studies of one self-form disagree, their tasks should therefore be compared before their maps are treated as conflicting accounts of the same ability.

How self-related processing relates to consciousness likewise depended on which self-form and which aspect of consciousness was being compared (Fig. 6). We predict that the sense of being a self fades when contributions that place experience in a personal context weaken while systems supporting perception remain active, so that seeing and hearing continue without seeming to belong to “me”. Regional patterns in such states should then resemble the maps of reflective self-forms less while still resembling maps of conscious perception and attention. Reports from experienced meditators and from participants given 5-MeO-DMT motivate this prediction [29, 30].

Pattern, predictive and reuse accounts concern which systems contribute to the self [33, 35, 36]. On how those contributions combine, graded composition predicted best in our comparison, and it resembles compositional codes, in which contributions learned in familiar circumstances are recombined into new behaviour and knowledge [45, 46]. A continuous space does not require gradual change over time, because configurations could still switch abruptly [47].

The coalition account must explain why a person feels like one continuing self. We propose that continuity comes from recurring members and from the shared name “self”. Every self-form reached midline and insular territory, though at different locations within it, so some of the same systems may take part in many configurations without any one location taking part in all. That these systems also serve general functions does not matter, because continuity requires recurrence, not self-specificity. Self-reference also binds perception and memory [48], and we suggest that applying “I” to bodily feelings, memories and actions marks them as belonging to the same person. This identification does not presuppose an owner, because applying “I” is itself one of the recurring activities. Continuity would then rest on identification, through which changing activities come to feel like “me”, and not on a part that persists [1, 2].

The account also bears on mental health. The self-forms closest to clinical themes (Negative self-reference, Monitoring and insight, and Body image/identity) showed different regional patterns. Clinical disturbances of the self may therefore involve particular configurations rather than the self as a whole. We predict that wellbeing tracks the ability to enter, sustain and leave configurations as circumstances require. Rumination would then be a failure to leave a self-critical configuration, which recurs even when attention is needed elsewhere. Work relating depressive rumination to the putative default-mode network, and wellbeing to separable self- and visual-attention-related networks during naturalistic viewing, motivates this prediction [49, 50]. Contemplative training could act in two ways [51]. Practices that train noticing thoughts as thoughts should make configurations easier to leave, whereas practices that question identification should loosen identification with any one configuration.

Several limitations qualify these conclusions. Each observation summarises a publication. Without within-person or temporal observations, graded patterns could reflect mixtures of discrete states in different people or at different moments, which an experiment comparing the three models on withheld single trials within individuals would test. Publications report selected coordinate peaks and maps are thresholded, so a weak contribution shared by all self-forms could escape detection. We estimated self-forms from researchers’ descriptions, no contrast was verified by a human as isolating self-related processing, and two categories concern clinical populations, so populations, methods and accompanying tasks may explain some differences attributed to self-forms. Abstracts can name reported regions, but categories regrouped from filtered abstracts retained the pooling advantage (Supplementary Note 1). The evidence for context was small and was not corroborated by log loss. Components are statistical stand-ins from one model family fitted only to cortex, and their correspondence to biological systems remains to be established.

In conclusion, in the chariot dialogue as in bundle theories, a name such as “chariot” or “self” refers to parts arranged to work together, with no further owner behind them [6]. Our account gives that view a neurobiological form. Brain systems that also do other work come together in changing coalitions to recognise a body, guide an action or remember a life, and “self” names what these systems do together as their contributions change.

## 4 Methods

### 4.1 Design

The analyses follow the three questions of the Results. The first is whether the self-forms share a core, the second is where the variation among self-related patterns comes from, and the third is how systems combine. All three use one corpus of publications, grouped by the wording of their titles and abstracts into ten categories, each an estimate of a self-form. Each section below gives the purpose and the main steps of an analysis, and the Supplementary Information gives its full specification.

### 4.2 The corpus and its ten self-forms

#### 4.2.1 Search and screening

Publications were selected in three steps, a search, a screen for relevance to the self and a screen for eligible methods. We searched NeuroStore, a database of neuroimaging publications and the coordinate tables extracted from them [38]. A fixed export held 32,411 records with reported coordinates. The query combined terms for self-related topics, such as self-reference, own face, body ownership, autobiographical memory and interoception, with the names of experimental paradigms and 75 theoretical labels, and it returned 2,068 candidate records after duplicates were removed.

Three language-model families (gpt-5.5, claude-opus-5 and gemini-3.1-pro-preview) then read each title and abstract. In the first screen, the models judged whether a publication investigated self-related processing or used self-related words only incidentally, as in self-paced presentation. A record passed only if all three models judged that it investigated self-related processing, and 1,023 did. The second screen removed designs that could not supply the evidence we needed, which were studies of nonhuman animals, of brain structure alone, of lesions, and of single patients or small case series, and secondary research such as reviews. A record was removed only when at least two models found explicit evidence of such a design, and records with unclear methods were kept. This screen removed 108 records and left 915 (Supplementary Method 1 gives the query, the criteria and the prompts).

Selection was made at the level of the publication. All the coordinate tables of an eligible publication were used, whatever its tasks, comparisons or control conditions, so differences between publications in baselines and statistical analyses can contribute to differences between self-forms.

#### 4.2.2 Publications, maps and regional profiles

Records were matched to publications by DOI, PubMed identifier and title with year, so that no publication counted twice, and the 915 records resolved to 912 publications. Each record kept its own text for the grouping described below, but every analysis of reported coordinates kept the records of a publication together.

Coordinates reported in Talairach space were transformed to Montreal Neurological Institute (MNI) space. In the primary analysis, which we call the all-coordinate analysis, we treated coordinates in an unknown or other space as MNI, which keeps the largest sample. To test that assumption, we excluded those coordinates in a known-coordinate analysis. Both were run throughout unless stated otherwise (Supplementary Note 12). Activation likelihood estimation (ALE) starts from a modelled-activation map for each analysis, which spreads each reported location over neighbouring voxels to allow for uncertainty about where it lies. Sample sizes for individual analyses could not be verified, so we used the spread that corresponds to a sample of 20 for every map and tested broader and narrower spreads with assumed samples of 10, 40 and 80. The analysis maps of a publication were combined by taking the largest value at each voxel, so that its overlapping contrasts did not count as independent evidence [52]. This gave one map for each of 911 publications. We then used ALE to test whether the publications of each self-form reported nearby locations more often than expected by chance, with NiMARE 0.21.0 [53, 54] on a 2 mm brain mask of 228,483 voxels, which includes white matter and ventricles. The maps of the ten self-forms are examined together, so they were corrected together. Benjamini–Hochberg correction held the false discovery rate (FDR), the expected share of false positives among the voxels declared significant, at q < 0.05 across all voxels of all ten maps (Supplementary Method 2 and Supplementary Tables 2 and 3 give sensitivity analyses).

A regional profile records, for every region of an atlas, whether the publication reported a location in or near it. The atlas has 1,041 regions, made up of 1,000 Schaefer cortical parcels [55], 15 Harvard–Oxford subcortical regions and 26 cerebellar regions from the AAL atlas. A region counted as reported when any of its voxels lay within 6 mm of a reported location, a margin that allows for uncertainty in localisation. Profiles were available for 910 publications in the all-coordinate analysis and 784 in the known-coordinate analysis (Supplementary Method 2). For summaries by division, the cortical parcels were also assigned to the 17 putative resting-state networks of Yeo et al. [56], which with subcortex and cerebellum make 19 divisions.

#### 4.2.3 Grouping titles and abstracts into categories

We grouped publications by what their titles and abstracts say, so that reported regions could be compared between groups that had been formed without reported coordinates. A language-model embedding turns a text into a list of numbers such that texts with similar wording receive similar lists, and we used the 1,024-number embeddings of BAAI/bge-large-en-v1.5. UMAP reduced each embedding to ten dimensions [57]. In this reduced form, each record was linked to the records most similar to it, which makes a graph, and Leiden clustering divided the graph into groups of closely linked records [58]. The number of groups was not set in advance and can depend on the settings of UMAP and Leiden clustering. We therefore ran 27 combinations of settings and kept the number of groups that occurred most often, which was ten. Among the solutions with that number, we selected the one that agreed best on average with all the others. We call the ten groups categories. We treat each category as an estimate of a self-form, and below we write category where the grouping of text is at issue and self-form elsewhere. Language models proposed a name for each category after membership was fixed, and one of them chose among the proposals without knowing which model had made each.

Supplementary Method 3 reports how distinct and how stable the categories are, including the proportion of records that lie closer to another category than to their own (Fig. 1).

#### 4.2.4 Prediction, scoring and uncertainty

Several analyses predict something about publications that a model has not seen, and they share the following conventions. Publications were divided into five groups, or folds, with similar proportions of each self-form. A model was fitted to four folds and tested on the fifth, in rotation, and the procedure was run twice with different assignments of publications to folds. We prepared the data and chose all settings using the training publications only, although the ten categories themselves were fixed from the text of the whole corpus. The configuration models followed the scheme described in their own section.

Predictions of which regions a publication reported were scored by Brier loss, the mean squared difference between the predicted probability that a region was reported and the outcome, which is zero or one [39]. Lower values mean better prediction. A gain is the reduction in loss relative to a comparison model, and where the Results give a gain as a percentage it is divided by the loss of that model. Because whether a region was reported is a yes-or-no outcome, part of the loss remains even when a predicted probability is correct, so a reliable gain can be a small percentage of the loss. Log loss was a secondary score that penalises confident errors more strongly. Classification was scored by balanced accuracy, the proportion classified correctly within each self-form averaged over self-forms.

Predictive models included reporting characteristics, to account for differences in how publications report. These were publication year, log(1 + reported locations), log(1 + analyses) and the fraction of coordinates in an unknown space. A second set of predictors added 14 task and clinical indicators, extracted from the text by fixed rules (Supplementary Method 4).

Uncertainty was estimated by resampling publications, usually 9,999 times, with each publication’s regions and repeated predictions kept together. Intervals came from these bootstrap samples, and p values from randomly reversing the sign of each publication’s difference in loss between two models. Both hold the fitted models fixed, so they describe uncertainty in these predictions alone (Supplementary Method 4).

### 4.3 Is there a core?

#### 4.3.1 Self-forms have different maps

The first step towards asking whether the self-forms share a core was to test whether their regional profiles differ at all. Self-forms could differ in how much their publications report, or in how many concern clinical groups, as well as in the regions themselves. We therefore compared the observed differences with those obtained after shuffling the self-form labels among publications that were alike in both respects. Publications were divided into four groups by the amount they reported and, within each, by whether the title contained a clinical term such as patient, disorder or depression. Labels were shuffled only within these groups, and each publication kept its whole regional profile.

For the test across all ten self-forms, we summed, over the 1,041 regions, a statistic for the association between self-form and whether a region was reported and compared the sum with those from 1,999 shuffles [59]. To test individual regions, we set a threshold from the distribution of the largest regional statistic across shuffles, which controls errors across regions. The 45 pairs of self-forms were then compared with 9,999 shuffles and Holm correction. The test across all ten was repeated in the 720 publications whose title or abstract contains the word self (Supplementary Method 4; Supplementary Note 1).

Earlier syntheses pooled studies across self-forms, so we examined what a pooled analysis shows and what it loses. The pooled map was an ALE map of all publications together, tested as one literature and corrected separately. We calculated the share of each self-form’s significant voxels that lie inside the pooled map and the share of the pooled map that each self-form covers, and we correlated each self-form’s map with the pooled map before thresholding.

In the predictive comparison, we asked whether knowing the self-form of a publication improves prediction of the regions it reported, compared with a single pooled pattern. In each training set, the probability that a region is reported was estimated for each self-form as the proportion of its publications that reported it. Small self-forms give unstable proportions, so each estimate was drawn towards the proportion among all training publications, as if 20 publications at that pooled proportion had been added. The pooled comparison model used the proportion among all training publications. Overall gains weighted self-forms equally, and comparisons for individual self-forms were FDR-corrected (Fig. 3c,d; Supplementary Table 5; Supplementary Note 1).

Because existing divisions have two or three kinds of self, we also asked whether a finer description of what a publication is about predicts its regions better than a coarser one. Within each training set, the embeddings were reduced to their 30 principal axes, the directions along which they vary most, and grouped anew by k-means clustering into three groups and into ten. Each unseen publication was assigned to a group from its text alone before its regions were predicted. The two groupings were compared in models that also contained the reporting characteristics and the task and clinical indicators. This comparison belongs to a family of seven, FDR-corrected together, which also included tests of the original ten categories and a continuous description of the text (Supplementary Note 1; Supplementary Fig. 2b; Supplementary Tables 6 and 7).

#### 4.3.2 No location is common to every self-form

A core requires a location that is present in the maps of all ten self-forms. For every voxel we counted how many of the ten corrected maps contained it and recorded which self-forms those were (Fig. 4; Supplementary Note 2). To ask whether every map nevertheless reached the candidate territories, we counted its significant voxels within regions of the Harvard–Oxford atlas that stand for medial prefrontal cortex, posterior cingulate/precuneus and anterior insula. In the atlas used for regional profiles and in the anatomical atlas introduced below, we also counted the regions in which every map had at least one, or at least 20, significant voxels, which need not be the same voxels (Supplementary Note 2).

A contribution shared by every self-form might be too weak to reach significance in all ten, which would leave no common voxel even though the contribution exists. To gauge how easily that happens, we simulated a single feature whose probability of being reported was raised in every self-form by construction and counted how often tests detected the increase in all ten [60] (Supplementary Note 2; Supplementary Table 8).

#### 4.3.3 Differences between self-forms are distributed across the brain

If one region or division carried what distinguishes the self-forms, their differences would be concentrated there. We tested this with three analyses, first describing how widely each map extends. For each of the 19 divisions, we calculated the percentage of a map’s significant voxels that fall within it and counted the divisions holding more than 1%, a descriptive threshold that sets aside divisions holding only a small fraction. A second atlas of 167 anatomical regions, the FreeSurfer Destrieux and aseg parcellation [61], gave a count that follows anatomical boundaries, in which a region counted when it held at least 20 significant voxels. Larger maps reach more regions through size alone, so each map was compared with 10,000 randomly grown, connected territories containing as many voxels as the map has within the atlas, with Holm correction (Supplementary Note 3).

Second, we tested whether this spread carries information that distinguishes the self-forms. A classifier predicted the self-form of each publication from its regional profile, by logistic regression with weights that balance the self-forms. We compared the classifier given all 1,041 regions with versions restricted to one region, to one division or to 50 neighbouring regions, each chosen within the training set. The best division contains fewer regions than the whole brain, so we also fitted a version with the same number of regions sampled uniformly from the whole atlas, to test whether the number of predictors explains the difference. Whether the whole-brain classifier exceeded chance was tested with 999 shuffles of the labels (Supplementary Note 3, which also describes the simpler model used for a single region; Supplementary Tables 9 and 10).

Third, we asked whether distinguishing information survives removal of the regions that carry most of it. Regions were ranked by the size of their largest coefficient across self-forms in the classifier fitted to the training set. The highest-ranked 10, 50, 100 or 200 were removed and the classifier was refitted to the remainder, and 50, 100 or 200 randomly chosen regions were removed for comparison (Fig. 5e).

### 4.4 Where does the variation come from?

#### 4.4.1 Context within a self-form

If each self-form had one fixed set of regions, the context of a study should not predict which regions it reports. Titles and abstracts describe the tasks, stimuli and participants of a study, and we used them as a description of its context. To isolate differences in context within a self-form, we subtracted from the embedding of each record the mean embedding of its category, calculated in the training set. What remains describes how a study differs from others on the same self-form. We summarised these remainders by their principal axes and added the values of each record on these axes to a comparison model that already contained the category and the reporting characteristics, with or without the task and clinical indicators. The number of axes and the strength of a ridge penalty, which discourages large regression coefficients, were chosen by validation within the training set, and the comparison model alone could also be selected. The four Brier comparisons, from two comparison models in two coordinate analyses, were FDR-corrected together (Supplementary Fig. 2c; Supplementary Table 11).

An abstract can state the findings that are being predicted, so in a further analysis we reduced that information, omitting titles and removing sentences that state an outcome, name a brain region or give a statistical result, by rules fixed before any coordinates were loaded. The original text and the filtered abstracts were then compared on the same 867 eligible publications, with Holm correction over the three comparisons for each score (Supplementary Note 4; Supplementary Table 12).

#### 4.4.2 The aspect of consciousness

Here we asked whether correspondence between maps of self-related processing and maps of consciousness depends on which self-form is paired with which aspect of consciousness. The self literature was compared with the corpus of a companion meta-analysis of consciousness [40], whose seven groupings of research each concern one aspect of consciousness. The 113 publications found in both were removed so that none contributed to both sides, which left 799 self-only and 465 consciousness-only publications. For each of the ten self-forms and each of the seven groupings, a reporting-frequency map gave the proportion of its publications that reported each of the 1,041 regions.

Correlating the map of every self-form with the map of every grouping gave a table of ten rows and seven columns, which two statistics summarised. The first is a measure of how much the rows differ from one another, and so of whether self-forms have different profiles of correspondence across the groupings. The second is a measure of the pattern that remains after the average of each row and each column is subtracted, and so of whether particular pairings stand out once the tendency of some maps to resemble all others is removed. Both were compared with 9,999 shuffles of the self-form labels within the groups of similar reporting and clinical content, with Holm correction over the two statistics, and the 70 individual pairings were FDR-corrected. The pairings named in the Results were selected to illustrate the table. In 2,000 resamples of publications, we counted how often the two self-forms kept their opposite orderings of correspondence with the two groupings in the same resample (Fig. 6; Supplementary Note 5).

#### 4.4.3 Systems shared with other functions

If self-forms draw on general systems, their maps should resemble maps of functions studied without reference to the self. We compared the unthresholded ALE map of each self-form with maps of 50 topics from Neurosynth version 7 [62], where a topic is a set of words that recur together across a broad imaging literature. So that shared publications could not create resemblance, we rebuilt the topic maps after removing publications identifiable in our corpus and any others with self in the title, which removed 549 of 14,371 publications. Some locations are reported often whatever the topic, and this general pattern of reporting density was removed from both maps before they were correlated. Each correlation was compared with 10,000 phase-randomised versions of the self-form’s map, which keep its spatial smoothness while scrambling where its values fall, and FDR correction covered all 500 comparisons. Fig. 2k shows 20 topics, selected by name before the correlations were inspected (Supplementary Note 6).

Neural reuse also predicts that regions reported together in self-related studies are organised as they are across imaging studies in general. To describe that organisation, every pair of the 1,041 regions was linked by the correlation, across publications, between the two regions’ entries in the regional profiles. Diffusion embedding in BrainSpace [63] then ordered the regions along continuous axes called gradients, on which two regions lie close together when their links to all other regions are similar. We estimated 25 gradients and displayed and tested the first ten. To measure their reliability, the corpus was split into random halves 25 times, the gradients of each half were matched one to one with those of the full solution, and the matched gradients of the two halves were correlated.

To compare the gradients with imaging research in general, a sample of 914 publications that the self query did not retrieve, matched to the corpus on year and number of reported locations, was processed identically. Each self gradient was compared with all 25 general-imaging gradients and its strongest absolute correlation was kept. Selecting the strongest of 25 correlations inflates it, and neighbouring regions have similar values, so chance was estimated from surrogate gradients that preserve spatial structure, with the whole search repeated on every surrogate. Two kinds of surrogate were used, from BrainSMASH [64] and from Moran spectral randomisation [65], each with 999 surrogates and FDR correction over the ten matches (Fig. 7; Supplementary Fig. 4; Supplementary Note 6).

To relate the gradients to the self-forms, each publication was placed on a gradient at the average gradient value of its reported regions. The association between self-form and position was the proportion of the variance in position that membership explains. It was tested with 9,999 shuffles of the labels within the groups of similar reporting and clinical content, with FDR correction over the ten gradients (Supplementary Note 6).

### 4.5 How do systems combine?

#### 4.5.1 The prediction task

The three accounts were compared on how well each predicts a part of a publication’s regional pattern that it has not been shown. A model was given the regions reported in one hemisphere of an unseen publication, the observed hemisphere, and predicted those reported in the other, the concealed hemisphere. Both directions were predicted. A location near the midline could mark regions on both sides, because of the 6 mm margin, and so make part of the prediction automatic. Only cortical voxels at least 8 mm from the midline were therefore eligible (|*x*| ≥ 8 mm), which keeps the two sides at least 16 mm apart. The parts of the atlas parcels that remained, which we call fragments, numbered 489 on the left and 498 on the right. Subcortex and cerebellum took no part in this task. A stricter version used only the 728 parcels that lie wholly outside the excluded band (Supplementary Note 7).

#### 4.5.2 The three models

All three models start from the same description of how much a publication reports (Supplementary Note 8). Let *X*_*ij*_ indicate whether publication *i* reported fragment *j*. For each fragment, a logistic regression on reporting characteristics gave *o*_*ij*_, the log odds that publication *i* reports it before any pattern is added. The characteristics were publication year, log(1 + reported locations), log(1 + analyses), the fraction of coordinates in an unknown space and, from the observed hemisphere alone, log(1 + fragments reported). The regions reported in the concealed hemisphere were used only for scoring.

The inventory model describes each publication by one of *K* complete regional patterns. The variable *c*_*i*_ says which pattern applies to publication *i*,

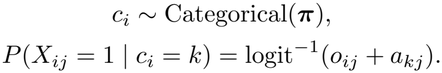

Here, *π*_*k*_ is the prevalence of pattern *k* and *a*_*kj*_ adjusts the log odds of fragment *j* when that pattern applies. The inverse logit turns log odds into the probability that the fragment is reported. For an unseen publication, the model calculated the probability that each complete pattern applied, given the observed hemisphere. It then averaged the predictions of the patterns by those probabilities, so the average reflects uncertainty about which complete pattern applies.

The graded model builds a pattern from *R* components. Each component is a fixed set of weights across fragments, and its contribution to a publication is multiplied by a strength that varies continuously,

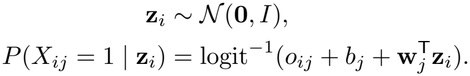

Here, z_*i*_ holds the strengths of publication *i*, one for each component, w_*j*_ holds the weights of the components on fragment *j*, and *b*_*j*_ is an intercept for the fragment. The model multiplies the weight of each component by its strength and adds these contributions for fragment *j*. Raising the strength of a component therefore raises the probability of reporting where its weight is positive and lowers it where its weight is negative. The strengths of an unseen publication were estimated, with their uncertainty, from the observed hemisphere alone, and predictions were averaged over that uncertainty.

With the on/off model, we tested whether graded strength matters, or only the freedom to combine components. It has the graded model’s equation for the probability of reporting, with each component either present or absent,

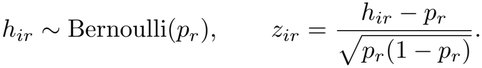

Here, *h*_*ir*_ indicates whether component *r* is present in publication *i* and *p*_*r*_ is the prevalence of that component. This scaling gives each *z*_*ir*_ a mean of zero and a variance of one, as in the graded model, while allowing it only two values.

Predictions summed exactly over all 2^*R*^ combinations of present and absent components, which number 1,024 for ten components.

In the terms of the Results, a component stands in for a system. The strengths of a publication’s components define its configuration, and the strength of each component supplies one axis of the fitted configuration space. No model was given the categories or the gradients.

#### 4.5.3 Fitting and choice of settings

The size and penalty of each model were chosen before its predictions were tested. Five outer folds supplied the unseen publications, and within the training set of each, settings were chosen on two inner folds to give the lowest Brier loss on concealed fragments. *K* was chosen from {1, 2, 3, 5, 10, 20} for the inventory model, *R* from {1, 2, 3, 5, 10} for the graded and on/off models, and each model’s penalty from {1, 10, 100, 1000}. In a further comparison, the on/off model was fixed at ten components. The inventory and graded models were fitted with the Adam optimiser [66, 67]. Supplementary Note 9 gives the fitting procedure and its checks, and reports simulations that tested whether the procedure could favour either a discrete or a graded account.

#### 4.5.4 Scoring

Losses from the two directions were combined in proportion to the number of fragments predicted, which weights the 987 fragments equally, and publications were then weighted equally. Brier loss was the primary score and log loss the secondary one. Comparisons with the inventory model were based on 9,999 paired resamples of publications, with FDR correction over five Brier comparisons. Comparisons with the on/off model were based on 10,000 resamples, with Holm correction over both coordinate analyses and both scores.

To see where the advantage of the graded model lies, the Brier loss of each model was separated into two parts for each publication and each direction of prediction. The first is the squared difference between the mean predicted probability of reporting and the proportion of fragments reported, which concerns how much a publication reports. The second is the mean squared difference between the predicted and the reported patterns after the mean of each is subtracted, which concerns which regions it reports (Fig. 8d; Supplementary Table 17). Supplementary Note 10 describes further scoring of the same models (Supplementary Tables 18, 19, 20 and 21).

#### 4.5.5 Illustrating the fitted configuration space

Fig. 8e–h shows where publications lie in the fitted graded model and what regional patterns its combinations produce. The figure is based on a single model with ten components, from the first outer fold of left-to-right prediction in the primary analysis. For display, principal component analysis of the estimated strengths of its 728 training publications gave three axes that hold 40.8% of the variation in strengths. Paths between self-forms were smooth curves through all ten dimensions, steered by the mean strengths of self-forms, and points along a path were converted into regional patterns at one set of reporting characteristics. The paths are constructed illustrations of patterns that the model permits (Supplementary Note 11).

## 4.6 Use of artificial intelligence

Large language models assisted with screening publications, proposing names for categories and labels for gradients, writing analysis code, generating figures, and drafting and editing the manuscript. The authors made the scientific decisions and take responsibility for the analyses, the interpretations and the final manuscript. No language model was treated as an author.

## 4.7 Data availability

Corpus metadata, screening decisions, statistical maps and source-data tables are indexed in the project repository at https://github.com/jskipper/distributed-self. A source-data index links every reported analysis to its inputs, settings, publications, predictions and the source data of its figure or table. The NeuroStore export was retrieved July 2026, and its SHA-256 checksum is recorded in the repository.

## 4.8 Code availability

Code, model prompts and outputs, the software environment and the workflow are indexed in the same repository.

## Acknowledgements

J.I.S. thanks Ursula Oliveira for her support, patience and enthusiasm for “self” research. We thank Bailey Bova at Anthropic for assistance.

## Author contributions

J.I.S. conceived the study, developed the methodology, curated the data, performed the analyses, prepared the figures and drafted the manuscript. All authors contributed to writing the manuscript.

## Ethics declaration

Not applicable.

