## Supplementary Information for "“The self” as shifting coalitions of brain systems in a continuous configuration space"

**Supplementary Tables** **38**

**Supplementary Figures** **57**

**References** **62**

#### Reading guide

The Supplementary Information follows the main text and has the same headings. Supplementary Methods 1–4 take the four headings under which the Methods describe the corpus and its ten self-forms. Supplementary Notes 1–11 take the headings under which the Results and the Methods address the three questions of the paper. Supplementary Note 12 summarises the known-coordinate analysis across all of them. The Supplementary Tables and Figures are numbered in the same order, apart from Supplementary Tables 22–27, which follow the others. Each Method and Note begins with links to the Results and Methods of the same heading, gives its results next and ends with the specification of its analyses.

We use the terms of the main text. A category is a group of publications with similar titles and abstracts, and we treat each category as an estimate of a self-form. We write category when discussing how the texts were grouped and self-form elsewhere. A publication is said to *report* a location or region when it supplied a coordinate assigned to it. The all-coordinate analysis is the primary analysis, and the known-coordinate analysis repeats it without coordinates in unknown spaces. A division is one of 17 putative cortical resting-state networks, the subcortex or the cerebellum. The inventory, on/off and graded models are the three models of how systems combine. The source-data index in the repository links every figure, table and analysis to its records.

### Supplementary Methods

### *The corpus and its ten self-forms*

#### Supplementary Method 1 | Search and screening

*Related main text. Results 2.1 and Methods 4.2.1.*

The search was meant to be broad and to avoid imposing a small set of theoretical kinds in advance. We searched a fixed NeuroStore export of 32,411 records with reported coordinates, of which 32,357 had abstracts of at least 80 characters. The query combined core self-related vocabulary, further terms for experimental paradigms, the word self standing alone in a title, terms for social cognition and embodiment when a self-related term accompanied them, and 75 theoretical labels. Duplicate records were removed by DOI, then PubMed identifier (PMID), then the database's own identifier, and 2,068 candidates remained. The complete query is the file `config/self_candidate_query.py` in the repository.

Screening separated publications that investigate self-related processing from those that use self-related words only incidentally, as in self-paced presentation. Three language-model families were used, with the archived identifiers `gpt-5.5`, `claude-opus-5` and `gemini-3.1-pro-preview`. In the first screen, each model read the title and abstract for an explicit self-related comparison, state or group difference, or an informative association with a measure of the self. The labels of analyses were withheld, and a record passed only if all three models agreed. The second screen concerned methods. It removed studies of nonhuman animals, of brain structure alone, of lesions, and of single patients or small case series without a group analysis, together with secondary research, when at least two models supported removal. A removal required a supporting quotation, and the analysis labels supplied with a record could also count as evidence. No record was removed on an unclear judgement. Records were processed in batches of ten with explicit identifiers, and the prompts, responses, votes, quotations and judgements of confidence and codability were kept.

At this scale, publications and their extracted coordinates were not checked contrast by contrast. The analyses therefore describe the regions reported across the selected literature, and we accepted some imprecision in individual records in exchange for breadth.

#### Supplementary Method 2 | Publications, maps and regional profiles

*Related main text. Results 2.1 and Methods 4.2.2.*

The analyses summarise the same literature in different ways, and the number of publications differs with the summary. The 915 retained records resolve to 912 publications. One publication has no nonzero value in its modelled-activation map, which leaves 911 maps for activation likelihood estimation (ALE). Two publications have no reported location in any region of the atlas, which leaves 910 regional profiles. Excluding coordinates in unknown spaces leaves 785 maps and 784 profiles. Main Figure 1d and Supplementary Table 1 describe the corpus.

##### Coordinate spaces and the two coordinate analyses

Coordinates explicitly reported in Talairach space were transformed to MNI space with the Lancaster transformation in NiMARE. In the *all-coordinate analysis*, which is the primary analysis, we treated coordinates in unknown or other spaces as MNI. The *known-coordinate analysis* excludes those coordinates and keeps a publication when it still has eligible locations. Because it changes the sample, it is a sensitivity analysis of the same corpus. Results of the two are given below as obtained with all coordinates and with known coordinates, and Supplementary Note 12 summarises how they compare.

Where the database held several extractions of a publication's coordinates, regional profiles were built from the one with the most eligible locations, with ties broken by order in the source, and extractions were never merged. Analyses that predict from regional profiles excluded publications with no reported region. The estimation of gradients and the maps for the comparison with consciousness could keep a publication with an empty regional profile if it still had locations in an eligible space. A publication can therefore contribute to those analyses without receiving a position on a gradient. Publication identifiers and the counts for each analysis are indexed with the source data.

The inventory below gives the principal counts. The columns All and Known refer to the two coordinate analyses of the same corpus.

**Inventory of analyses**

| Analysis | Unit | All | Known | Principal exclusion or restriction |
|---|---|---|---|---|
| Grouping of text | Source record | 915 | n/a | Passed both screens |
| ALE maps | Publication | 911 | 785 | Empty maps excluded, and unknown spaces excluded from Known |
| Regional profiles | Publication | 910 | 784 | At least one reported region |
| Self and consciousness maps | Publication in one corpus only | 799 + 465 | 686 + 401 | 113 publications in both corpora removed first |
| Three models of how systems combine | Publication | 910 | 784 | Same publications as the regional profiles |
| Filtered abstracts | Publication | 867 | n/a | Fixed rule for aims, methods and retained sentences |
| Filtered-abstract ALE | Publication | 867 | 749 | Original and filtered categories, matched cohort |
| Explicit mention of self | Publication | 720 | 614 | The word self alone in title or abstract |
| Methodological uses of self discounted | Publication | 714 | 611 | Fixed rule for compounds such as self-paced |

#### Activation likelihood estimation and its sensitivity analyses

ALE allows for uncertainty around each reported location. So that several analyses from one publication did not count as independent evidence, exact duplicate locations were removed and the analysis maps of a publication were combined by taking the largest value at each voxel. ALE was estimated with NiMARE on a common 2 mm MNI152 mask of 228,483 voxels. The assumed samples of 10, 20, 40 and 80 were applied uniformly, because the sample sizes of individual analyses could not be verified. Voxelwise p values in the primary analysis came from the approximate null distribution in NiMARE, and correction of the false discovery rate (FDR) covered every voxel of every self-form's map as one family. The pooled map was corrected as a separate family.

In one sensitivity analysis, we corrected the family-wise error rate, the probability of any false positive across the tests, with 1,000 datasets of randomly placed coordinates. Its p values are those of NiMARE, $b/B$ bounded to $[1/B, 1 - 1/B]$, where $b$ counts the random statistics at least as extreme as the observed statistic and $B = 1{,}000$. They therefore differ from the rule in Supplementary Method 4. Tests of cluster size and cluster mass formed clusters at a p value of 0.001. The voxelwise test evaluated individual voxels directly. Results are given for $\alpha = 0.05$ for each map and for $\alpha = 0.005$, which adjusts across the ten maps. A second sensitivity analysis restricted the types of analysis. It removed sets of coordinates explicitly labelled as connectivity, electrophysiology, structural or lesion evidence, analyses restricted to a region or mixing whole-brain and restricted search, and sets containing a seed, and it kept sets whose type could not be resolved. This adds to the screening of publications, because an eligible publication can contain analyses of other types. Supplementary Tables 2 and 3 report these analyses, and Supplementary Note 2 describes what they show about shared territory.

#### Regional profiles, atlas and divisions

A regional profile is a record of whether a publication reported each region, whatever the ALE result at that region. The atlas combined 1,000 Schaefer cortical parcels, 15 Harvard–Oxford subcortical regions and 26 AAL cerebellar regions. A region counted as reported when at least one of its voxels lay within 6 mm of a reported location in MNI space. Versions of the cortical atlas that order their parcels differently were aligned voxel by voxel, and the orientation and affine of the atlas were matched to the ALE grid before voxels were counted. The 17 putative cortical resting-state networks, the subcortex and the cerebellum make 19 divisions. The networks provide an external division of the cortex for these comparisons.

##### How brain maps are displayed

Brain displays show the final ALE maps, estimated from one modelled-activation map per publication, and the exact counts of overlap. Maps of single self-forms and the pooled map are purple, and overlap is shown on one scale from one to eight. The views, used as appropriate, are lateral, ventral and medial fsaverage surfaces, subcortical structures, a SUIT flat map of the cerebellum [1] and eight fixed axial sections in MNI space. Projection to the cortical surface samples nine positions between the white and pial surfaces. Maps of discrete values take the most frequent value among the nearest voxels, background included, and continuous maps are sampled linearly. Subcortical and cerebellar views are filled from the values at their own locations. Display adds no thresholding and no smoothing, and the surfaces show results that were computed in the volume (main Figs. 2, 3 and 4).

The map of regional gains gives each surface point the fragment found most often among its nine samples, and then the saved gain of that fragment. Points without a fragment stay empty. The surface shows 956 of the 987 fragments, and the source tables and volumetric maps hold all 987. The colour scale for gains is symmetric and wide enough for every estimate. Maps of decoded probabilities are shown on a fixed scale from 0 to 0.30, with larger values given the top colour and labelled 0.30+. These conventions affect display only. The excluded band of cortex around the midline and the background outside cortex are drawn differently from small gains and low probabilities (main Fig. 8c,h).

#### Supplementary Method 3 | Grouping titles and abstracts into categories

*Related main text. Results 2.1 and Methods 4.2.3.*

This Method describes how the ten categories were chosen from the grid of solutions, and how distinct and how stable they are. The two-dimensional display in main Fig. 1 illustrates the categories. We measured their separation in the original embeddings, numerical representations of each record's title and abstract.

##### Choosing the ten categories

The grid contained 27 combinations of settings for UMAP and Leiden clustering, which produced between seven and 24 groups. Ten was the most frequent number and the only one to occur in six combinations. Among those six solutions, we selected the one with the greatest mean agreement with the other 26, including solutions with other numbers of groups. Agreement was measured by the adjusted Rand index, which scores how far two groupings place the same records together. The selected solution had 30 neighbours, resolution 1.0 and seed 73, and its mean agreement was 0.722. The stability of a record was the proportion of the 27 solutions in which it sat in the cluster that held most members of its selected category, and mean stability was 0.819. We therefore treat the ten categories as a working grouping, chosen from the settings explored.

#### Separation and stability of the categories

Separation was measured in the original embeddings of 1,024 numbers by cosine distance, which compares the directions of two embeddings. The silhouette of a record is positive when the record is closer on average to its own category than to the nearest other category. A second measure compares the distance from a record to the centre of its own category with its distance to the nearest other centre. Across 915 records the mean silhouette was 0.0844, 18.4% of records had negative silhouettes, and 10.9% were no closer to their own centre than to another. The mean distance between pairs of records in the same category was 0.1905, and the mean distance from a record to its own centre was 0.0991. Mean silhouettes for the ten categories, in the order used throughout, were 0.0811, 0.0610, 0.0769, 0.0120, 0.0547, 0.1304, 0.1806, 0.1236, 0.1134 and 0.1715.

Self-appraisal was the least separated category, with 40% negative silhouettes, and its centre lay closest to that of Trait evaluation, at a cosine distance of 0.0167. Interoception and Face/voice recognition lay farthest apart, at 0.0795. To test stability, the corpus was split into complementary halves 100 times, and centres estimated from one half assigned the records of the other. The assignments agreed with the fixed categories for 85.1% of records, or 85.4% when the categories are weighted equally. Related categories therefore overlap considerably, but most records keep their category when the centres are estimated from other records.

With one record for each of the 912 publications, the mean silhouette was 0.0843, negative silhouettes made up 18.5% of records, the mean distance within categories was 0.1905 and the equally weighted agreement between halves was 85.2%. With a further 1,000 bootstrap samples, drawn within categories and keeping duplicated records, we recalculated the centres and the distances with membership fixed. The observed margin between a record's own centre and the nearest other centre lay below the 2.5th percentile of its bootstrap distribution, both with all records and with one record per publication, so we report the bootstrap ranges as descriptive only (main Fig. 1c).

The 168 negative-silhouette records had mean grid membership stability 0.663, compared with 0.854 among other records (Supplementary Table 24 gives stability by category). Clinical keyword indicators occurred in 30.4% and 38.4%, respectively. Held-out anatomical classification is scored by publication, and it was more ambiguous for the publications these records represent. Their mean classifier probabilities were 0.149 for their own category and 0.128 for the category nearest in wording, compared with 0.198 and 0.113 for other publications. These class-balanced classifier probabilities are descriptive and do not establish mixtures. The pooling gain stayed positive when scoring was restricted to non-boundary publications without refitting. The clinical, classifier and pooling values are in the source data.

#### Naming the categories

Three language models proposed names after membership was fixed. One of them, Claude, then chose among the proposals with their sources hidden, its own proposal among them. The proposals and the choice are kept with the source data, and the names had no part in forming the categories (Supplementary Table 1).

#### How the displays of text were made

The two-dimensional UMAP was computed from the original embeddings with cosine distance, 30 neighbours, a minimum distance of 0.1 and seed 42, and saved identifiers link its points to the selected categories. The word cloud shows the 50 most frequent eligible terms that occur in at least five records, after words of the same family were merged by fixed rules and generic terms of reporting were removed. The dictionary, the exclusions, the frequencies and the placements accompany the source data (main Fig. 1a,b).

##### Specification of clustering and naming

The text given to the embedding model placed the title before the abstract, and each embedding was scaled to unit length. The ten-dimensional UMAP was computed with cosine distance and a minimum distance of 0.1. In that space, each record was linked to its nearest neighbours by Euclidean distance, with the same number of neighbours as the UMAP, and a link in either direction joined two records in an unweighted graph. Leiden clustering optimised the RB configuration partition with the stated resolution and seed.

The grid crossed neighbourhood sizes of 15, 30 and 50, resolutions of 0.5, 1.0 and 1.5, and seeds 13, 42 and 73. Neighbourhood size sets how many nearby records are considered, resolution sets how finely the graph is divided, and the seed sets the random choices in the procedure. Clustering was always done in the ten-dimensional form and never in the two-dimensional display. A tie for the most frequent number of groups would have been settled in favour of the lower number. Names were proposed from the terms most distinctive of each category, by term frequency weighted by inverse document frequency (TF–IDF), and from representative titles. No brain coordinates were used in naming.

#### Supplementary Method 4 | Prediction, scoring and uncertainty

*Related main text. Methods 4.2.4.*

The unit of analysis is the publication unless stated otherwise. The grouping of text keeps the source records, because records of the same publication can carry different text. Every analysis of reported coordinates keeps the records of a publication together, so that they neither count as independent observations nor fall on both sides of a division into training and test publications.

##### Validation

The shared scheme consisted of two repeats of a fivefold division of publications, with each fold serving in turn as the test set and the remaining publications as the training set. The proportions of the self-forms were kept similar across folds. Each publication was predicted once in each repeat, and the models in a paired comparison were tested on the same unseen publications. Replacement of missing values, standardisation, projections of text, learned groups and the choice of settings were based on the training publications only. The original ten categories are the one exception, because they were fixed from the text of the whole corpus. The penalty of ridge models of text was chosen from 1, 10, 100, 1,000 or 10,000 by Brier loss in a twofold validation within the training set. Predicted probabilities were bounded to $[10^{-5}, 1 - 10^{-5}]$. The three models of how systems combine followed the scheme described in Supplementary Note 9.

##### Scores and weights

Predicted probabilities of reporting are scored by Brier loss, the mean squared difference between the predicted probability and the outcome, which is zero or one. Lower loss means better prediction. Log loss is a second score that penalises confident errors more strongly, and it is calculated for each region or fragment separately. A positive gain means that the named model has lower loss than its comparison model. Classification of self-forms is scored by balanced accuracy, which gives the self-forms equal weight.

Weights depend on the question. The tables of scores for the reporting and text models average losses over publications, whereas their paired comparisons and the pooling comparisons average gains within each self-form and then across self-forms with equal weight. The analysis of filtered abstracts weights self-forms equally for both scores and gains. The comparisons of the three models first combine the losses of the two directions of prediction in proportion to the number of fragments predicted, and then weight publications equally, so each of the 987 fragments has equal weight whatever its volume. Where an estimate that weights self-forms equally is also given, it is labelled as such. Losses calculated with one weighting cannot be subtracted to reproduce a gain calculated with another.

#### Reporting characteristics and indicators of tasks and clinical content

Publications differ in how many coordinates they report, when they were published and what tasks or populations they describe. Several analyses therefore include reporting characteristics as predictors, or keep broad differences in reporting and clinical content intact when labels are shuffled. The reporting characteristics are publication year, log(1 + reported locations), log(1 + analyses) and the fraction of coordinates in unknown spaces. Fourteen task and clinical indicators summarise clinical content, imaging and stimulus modality and broad task demands. They are read from the text and have not been validated against the experiments themselves.

Each indicator is a match to a regular expression, ignoring case, in the title and abstract joined together. A publication can match several indicators. A missing field would have been treated as empty text, but neither field was missing in either coordinate analysis. The expressions below are the rules as implemented. The eligibility of publications was decided in the screens and never from these indicators.

**Task and clinical indicators**

<table>
<tr><th>Indicator</th><th>Regular expression, ignoring case</th><th>All</th><th>Known</th></tr>
<tr><td>Clinical</td><td>patient|disorder|depress|schizophren|autis|anorexi|clinical</td><td>335</td><td>282</td></tr>
<tr><td>Resting</td><td>resting.state|rest.condition</td><td>94</td><td>73</td></tr>
<tr><td>Connectivity</td><td>connectivity|correlation.analysis|psychophysiological</td><td>222</td><td>180</td></tr>
<tr><td>PET</td><td>\bpet\b|positron.emission</td><td>14</td><td>11</td></tr>
<tr><td>Structural</td><td>voxel.based.morphometry|grey.matter|gray.matter|cortical.thickness</td><td>11</td><td>7</td></tr>
<tr><td>Visual</td><td>visual|picture|image|photograph|video</td><td>289</td><td>251</td></tr>
<tr><td>Auditory</td><td>auditory|voice|sound|listen</td><td>54</td><td>50</td></tr>
<tr><td>Motor</td><td>motor|movement|button|hand|finger</td><td>237</td><td>214</td></tr>
<tr><td>Memory task</td><td>memory|recall|retriev|recollect</td><td>208</td><td>175</td></tr>
<tr><td>Decision task</td><td>decision|judg|choice|rating|evaluat</td><td>398</td><td>335</td></tr>
<tr><td>Emotional task</td><td>emotion|affect|valence|negative|positive</td><td>475</td><td>402</td></tr>
<tr><td>Social task</td><td>social|other|empathy|perspective</td><td>526</td><td>447</td></tr>
<tr><td>Passive</td><td>passive|viewing|listening</td><td>89</td><td>75</td></tr>
<tr><td>Longitudinal</td><td>longitudinal|intervention|treatment|therapy</td><td>94</td><td>75</td></tr>
</table>

The numbers of publications in the ten self-forms were 146, 139, 135, 124, 87, 79, 58, 52, 46 and 44 with all coordinates, and 120, 106, 126, 108, 73, 70, 52, 48, 41 and 40 with known coordinates, in the order used throughout. Counts of each indicator in each self-form, and a hash of the code that extracts them, are indexed with the source data.

#### Groups of similar publications for shuffling labels

Several tests shuffle the labels of self-forms among publications that are alike in how much they report and in clinical content. They followed closely related rules for clinical content. For the tests of regional profiles, across all ten self-forms and between pairs, titles were searched for the seven word stems in the Clinical row and for `psychosis` and `bulimi`. The tests of positions on gradients and of correspondence with consciousness were based on the seven stems. Missing titles were treated as empty text in the pair, gradient and consciousness tests, and no title was missing in the test across all ten. Publications were divided into quarters according to how much they reported. For the test across all ten self-forms, this was the number of reported regions. For the tests of pairs, it was the number of reported locations, with quarters recalculated within each pair. For the gradient and consciousness tests, it was the number of reported locations within the relevant coordinate analysis. Shuffling within these groups holds broad measured differences constant, and other differences of task or population remain.

##### Uncertainty

Summaries of uncertainty differ in meaning. Bootstrap intervals for predictions resample whole publications while holding the fitted models fixed. They describe variation in the saved predictions, and they leave out both the uncertainty of refitting the whole procedure and the dependence between training sets that overlap. Ranges from shuffles describe the values expected under the stated randomisation. The ranges from resampling the categories, and from resampling the illustrative pairings with consciousness, are labelled as descriptive because they are uncalibrated. Intervals for the recovery simulations describe success across simulated datasets.

The default paired procedure consisted of 9,999 bootstrap samples that resampled publications within self-forms and 9,999 random reversals of the sign of each publication's gain. The regions and the repeated predictions of a publication were resampled together. Gains for pooling and for text were averaged within each self-form and then equally over the ten. The analyses of filtered abstracts, of the three models and of the two corpora depart from these defaults as their Notes describe.

##### P values from shuffles and surrogates

For our own tests by shuffling and by spatial surrogates, the p value was $p = (b+1)/(B+1)$, where $b$ is the number of randomised statistics at least as extreme as the observed statistic and $B$ is the number of draws [2]. This prevents a p value of zero when no randomised statistic exceeds the observed one. Two-sided tests of individual cells doubled the smaller tail probability and capped the result at one. Each analysis below states what was shuffled, the number of draws and the family of tests that was corrected together.

##### Software environment

We processed the data and made the figures with a versioned Python environment and the cited software [3–6]. The source-data index lists the environment and the dependencies of each analysis.

#### Supplementary Notes

### *Is there a core?*

#### Supplementary Note 1 | Self-forms have different maps

*Related main text. Results 2.2.1 and Methods 4.3.1.*

Three analyses lie behind the statement that the self-forms have different maps (main Fig. 2a–j). In the first we compare the regional profiles of every pair of self-forms. In the second we ask what is lost when the self-forms are pooled, and in the third whether ten groups of text predict reported regions better than three.

##### Differences between pairs of self-forms

Every pair of self-forms was compared directly on the regional profiles, which are not thresholded. Supplementary Figure 1 shows the size of the differences after adjustment for the groups of similar publications, with stars marking the pairs whose whole profiles differed after correction. Supplementary Table 4 reports both coordinate analyses. The whole profiles of 25 of the 45 pairs differed after correction with all coordinates and 21 with known coordinates, with 21 found in both. The comparisons concern what the publications of each self-form reported, across all their analyses.

The test across all ten self-forms was also repeated in the 720 publications whose title or abstract contains the word self, and the self-forms again differed ($p = 0.00010$, 10,000 shuffles).

##### What pooling across self-forms loses

Pooling can preserve many of the regions reported for a self-form while obscuring how often its publications report each region. We tested this by predicting the regions an unseen publication reported, either from the pattern of its own self-form or from a pooled pattern. The pattern of a self-form is the proportion of its training publications that reported each region, stabilised as the Methods describe, and the pooled pattern is that proportion among all training publications. The comparison concerns predicted probabilities of reporting and is separate from the overlap between thresholded ALE maps in main Fig. 3a,b. Positive differences below favour the pattern of the self-form.

For the overall gain, paired losses were averaged within each self-form and then equally across the ten. Against the pattern of all training publications, the gain in Brier loss was 0.000508 with all coordinates, with a 95% interval of 0.000248–0.000764, and 0.000554 with known coordinates, with an interval of 0.000248–0.000871. The two-sided p values, from reversals of sign, were 0.0005 and 0.0013, respectively (main Fig. 3d).

With two further comparison models, we tested how the contents and the size of the pool affect the gain. With the first, we asked whether unequal sample sizes or the stabilisation of small samples could explain it. For each self-form, 20 random sets of training publications were drawn from the pool, each as large as the self-form. The estimates for the self-form and for each random set were both combined with the pattern of all training publications, giving that shared pattern a weight equal to 20 publications. The losses of the 20 sets on unseen publications were then averaged. Against this comparison model the gains were 0.001272 with all coordinates, with an interval of 0.001011–0.001528, and 0.001406 with known coordinates, with an interval of 0.001094–0.001724, and both p values were 0.0001.

The second left the tested self-form out of the pool, so that the pooled pattern contained none of its training publications. The gains were 0.000916, with an interval of 0.000610–0.001218, and 0.001000, with an interval of 0.000638–0.001377, in the two coordinate analyses, both with $p = 0.0001$. The p values of these three overall comparisons are uncorrected, and they stand apart from the family of ten tests for single self-forms, which was FDR-corrected. Intervals resample publications within self-forms and hold the fitted models fixed. Complete values are indexed with the source data.

Gains were not uniform across self-forms. Autobiographical memory and Ownership and agency kept a corrected advantage in both coordinate analyses, and several other differences were uncertain. Trait evaluation went from an uncertain negative estimate with all coordinates to a corrected disadvantage with known coordinates ($q = 0.002$; main Fig. 3c and Supplementary Table 5). Supplementary Figure 2a extends the comparison to every pairing of a self-form that supplies the pattern with a self-form that is predicted. Rows give the self-form supplying the pattern, and prediction of one self-form from another need not equal prediction in the reverse direction.

#### Ten groups compared with three, and finer descriptions of text

The pooling comparison shows that distinguishing self-forms helps prediction, and it leaves open how fine the distinctions should be. We therefore compared coarser and finer groupings of text. These models predict regions from descriptions of studies. The three models of how systems combine are a separate analysis, in which one hemisphere predicts the other (Supplementary Notes 7–10).

The models with three and with ten groups learned their projection of text and their k-means grouping within each training set, and then assigned each unseen publication to a group from its text. The three groups are those that the procedure finds, and they correspond to no published division of the self. The original ten categories were fixed from the text of the whole corpus, and every model was fitted to the regional profiles of training publications only.

Supplementary Tables 6 and 7 compare models with a single shared pattern, reporting characteristics, reporting characteristics with task and clinical indicators, the original categories, the learned groups, a continuous description of text and reduced-rank descriptions. The continuous description consists of positions on the principal axes of the text embeddings. Reduced-rank descriptions limit how many patterns of regional variation the text can predict. The categories improved prediction over the same reporting characteristics, and ten learned groups kept distinctions that three lost (Supplementary Fig. 2b). More detail did not always help. The indicators themselves did not improve Brier loss, so the model with categories and indicators predicted no better than reporting characteristics alone, and it predicted worse with known coordinates ($q = 0.0072$; Supplementary Tables 6 and 7). The continuous description predicted worse on Brier loss than the reporting characteristics with task and clinical indicators, and the reduced-rank description of rank ten predicted worse than that of rank three. The continuous description had a lower log loss with all coordinates, 0.3139 against 0.3223, and the saved paired predictions give a gain of 0.009933107 over reporting and indicators (95% interval 0.008029716–0.011903081, $q = 0.000175$; Supplementary Table 23). This paired gain weights categories equally, whereas the absolute scores above weight publications equally.

#### Categories and predictions from filtered abstracts

We repeated grouping and prediction with the filtered abstracts of Supplementary Note 4. Titles were omitted and fixed sentence rules reduced statements of findings, anatomy and statistics. The same embedding model and 27-setting UMAP–Leiden grid gave tied modal counts of 8, 9, 10 and 13, each in 3 solutions. Following author feedback, we replaced the inherited smallest-on-ties rule. We kept the two middle tied counts and, of their representative partitions, chose the one with the higher mean adjusted Rand index with the other 26 grid solutions. This selected 10 groups on 867 eligible publications, with adjusted Rand index 0.515 against the original categories. The rule was amended after the initial eight-group result and before fitting the revised maps. No anatomical outcome was used in the revised choice.

These are newly formed groups rather than a selection of original categories (Supplementary Table 25). The original full-text grouping had a unique mode at ten, so its selection was unaffected.

Both training and scoring were restricted to eligible publications while preserving the original outer and inner fold memberships. Fixed original and filtered categories remained annotations formed from the whole eligible corpus. Principal axes, scaling, k-means for learned groups and regression settings were fitted to training publications only. Reporting characteristics and the original task and clinical keyword indicators were retained. Losses were averaged across repeats within publication before inference. We reported gains with the filtered categories weighted equally and with the original categories weighted equally, with FDR correction across seven model comparisons and, separately, three pooling comparisons within each weighting. The filtered categories improved prediction over all three pools and over reporting characteristics with or without indicators. The ten-versus-three gain was smaller and uncertain under original-category weighting ($q = 0.0726$), but positive under filtered-category weighting ($q = 0.0273$; Supplementary Table 22). Filtering can discard legitimate context as well as outcome language and does not certify outcome-free text.

For scale, reporting characteristics reduced Brier loss by 5.889% relative to an intercept-only model on the original all-coordinate publications and folds, and by 5.795% with known coordinates. This uses saved predictions with the original categories weighted equally (Supplementary Table 23). The seven model comparisons were also evaluated for log loss, with FDR correction separately within each coordinate analysis.

#### Specification of the tests of regional profiles

The test across all ten self-forms calculated, for each region, a Pearson statistic for the table of ten self-forms by reported or unreported, and summed it over regions. Its 1,999 shuffles moved self-form labels within quarters of the number of reported regions, crossed with the clinical-title indicator. The largest regional statistic in each shuffle gave the distribution that corrects the tests of individual regions for the family-wise error rate. A publication mentions self when the word stands alone, hyphenated compounds included, after normalisation of characters, in any of the texts that belong to it. Words that merely begin with self, and the word selves, do not count. The repeat of the test in the 720 publications that mention self had 10,000 shuffles within the groups defined in the full sample of 910, and its summed statistic was 16,312.261.

For each of the 45 tests of pairs, publications were divided into quarters according to their number of reported locations, using only publications in that pair. In each group of similar publications and each region, a score compared the number of reports in the first self-form with the number expected from the sizes of the two self-forms and their combined total of reports. Scores and their hypergeometric variances were summed across groups within each region, and the statistic for the whole profile summed, over regions, the squared total score divided by its total variance. A region with zero variance contributed zero, and a group containing only one of the two self-forms contributed no comparison. Labels were shuffled 9,999 times within groups, with Holm correction over the 45 pairs in each coordinate analysis. For description, the signed difference in each region averaged the differences within groups with weights $n_A n_B/(n_A + n_B)$, and the root mean square of these differences summarises the whole profile. Supplementary Method 4 gives the rules for clinical titles.

##### Specification of the models of text

The reporting characteristics were year, $\log(1 + \text{reported locations})$, $\log(1 + \text{analyses})$ and the fraction of coordinates in unknown spaces, and the second set of predictors added the 14 indicators of Supplementary Method 4. Each model predicted whether each region was reported by ridge least-squares regression with an intercept. Missing values of the characteristics were replaced by training medians, the characteristics were standardised within the training set, and fixed or learned groups entered as indicator variables. The models with three and ten groups fitted k-means clustering, with ten starts, to the 30 principal axes of the training embeddings after standardisation, and assigned test records from their text. The continuous model was based on the values on these 30 axes without grouping. The reduced-rank models predicted, from text, the residuals, the differences between observed reports and predictions from reporting characteristics and indicators. They kept three or ten singular directions, the main patterns of variation in those predicted residuals, and added the resulting residual predictions to the comparison model's predictions before probabilities were bounded. Supplementary Method 4 gives the grid of penalties and the bounds. Seven paired Brier comparisons formed one FDR family within each coordinate analysis. Three compared the original categories with models without them, one compared ten learned groups with three, two compared the continuous description with the indicators and with the original categories, and one compared rank ten with rank three.

#### Supplementary Note 2 | No location is common to every self-form

*Related main text. Results 2.2.2 and Methods 4.3.2.*

A core requires a location present in the maps of all ten self-forms. This Note reports how much of the significant ALE territory is shared and by which self-forms, how far each map reaches the candidate territories, and whether a weak contribution common to all ten could have gone undetected.

##### Territory shared by self-forms

In the all-coordinate analysis, 4,342 voxels were shared by at least four self-forms, 2,167 by at least five, 705 by at least six, 163 by at least seven and 19 by eight, and no voxel was significant in nine or ten maps (main Fig. 4a,c). The corresponding counts with known coordinates were 3,322, 1,524, 424, 47 and one, again with none in nine or ten. These counts concern the original ten maps. In the sensitivity analysis later in this Note, which regrouped publications from abstracts after filtering anatomical and result statements, 3 voxels (24 $mm^3$, 0.007% of the significant territory) at the border of right anterior insula and orbitofrontal cortex (MNI $x = 32$–$34$, $y = 24$–$26$, $z = -4$) were significant in all ten maps with all coordinates. With known coordinates none were, although up to nine maps were significant there. We do not treat so small an intersection as a core, because it depends on how publications are grouped. In the original ten maps these voxels were significant in seven or eight maps, and the Face/voice recognition map did not reach significance there ($z = 0.46$–$1.27$). Anterior insula also serves general functions, so sharing there would not by itself mark a region specific to the self. The broader spread that corresponds to an assumed sample of ten produced three voxels shared by nine maps and none shared by ten, and the narrower spread for an assumed sample of 80 also left no voxel shared by ten. Supplementary Tables 2 and 3 report the sensitivity to spread and to correction.

The Dice coefficient of two maps is twice the volume they share divided by the sum of their volumes. As the assumed sample rose through 10, 20, 40 and 80, and the spread narrowed, the mean Dice coefficient over pairs of maps fell from 0.175 to 0.132, and the share of the mask covered by at least one map fell from 22.4% to 16.3%. The largest number of maps sharing a voxel was nine under the broadest spread and eight otherwise, and no voxel was significant in all ten under any assumption (Supplementary Table 2).

The shared territory lay in familiar places. At least four self-forms shared 1,180 voxels in medial superior frontal and anterior cingulate cortex, 957 in posterior cingulate and precuneus, 612 in left anterior insula and inferior frontal cortex, 454 in dorsal medial frontal cortex, 360 in right anterior insula and 332 in left posterior superior temporal and angular cortex. At least six self-forms shared 157 voxels in left anterior insula, 157 in posterior cingulate and precuneus, 111 in right anterior insula, 106 in anterior medial superior frontal cortex and 86 in dorsal medial frontal cortex, and the remaining 88 voxels formed smaller clusters. A cluster here is a set of voxels connected through any of their 26 neighbours, and tissue that the atlas does not label was recorded separately.

Which voxels are connected and which self-forms share them are separate matters. The 19 voxels shared by eight self-forms formed five clusters of eight, five, four, one and one voxels. They also showed five different combinations of eight self-forms, and a combination can be spread over more than one cluster. The largest combination held ten voxels and lacked Face/voice recognition and Body image/identity. Autobiographical memory, Negative self-reference, Self-appraisal, Interoception and Self–other distinction are present in all 19 voxels, Face/voice recognition is present in two, and the five combinations contain 10, 5, 2, 1 and 1 voxels. Across all significant voxels there were 79 distinct combinations of four self-forms, 57 of five, 35 of six, 15 of seven and five of eight.

Main Figure 4b shows the voxels shared by at least four, six and seven self-forms. Panel c includes the result for eight, and panel d shows the three largest combinations at each level that hold at least ten voxels. All of these describe overlap among the corrected maps, and no further test of conjunction was applied. They show that the self-forms sharing a location change from place to place. Complete locations and memberships are indexed with the source data.

#### The candidate territories and the regions that contain every self-form

The counts for the candidate territories in Results 2.2.2 are counts of significant voxels in the maps of the all-coordinate analysis, within three territories defined from the Harvard–Oxford cortical atlas (maximum probability, 25% threshold, 2 mm grid). Medial prefrontal cortex comprises Frontal Medial Cortex and the voxels of Superior Frontal Gyrus with $|x| \leq 20$ mm. Posterior cingulate/precuneus combines the posterior division of Cingulate Gyrus with Precuneous Cortex. Anterior insula comprises the voxels of Insular Cortex with $y > 0$ mm. The atlas was reoriented to the ALE grid without interpolation. A count sums the significant voxels of a map anywhere within a territory, so two maps can both reach a territory at different locations. The counts describe extent, and no further test was applied within the territories.

We also counted the regions that contain significant voxels from every self-form. When one voxel from each sufficed, there were 8 such regions in the anatomical atlas of 167 regions, which Supplementary Note 3 describes, and 2 in the atlas of 1,041 regions used for regional profiles. When 20 voxels from each were required, there were 1 and 0, respectively. For the larger atlas, the assignment of voxels to cortical and subcortical regions was rebuilt from the stored atlases and checked against the saved shares for each division. At least one self-form had no significant voxel in the cerebellum, so no cerebellar region could qualify. A region can hold voxels from every self-form at different locations, so these counts are compatible with there being no voxel common to all ten.

Across the 45 original category pairs, cosine distance between text centres correlated with Dice overlap of significant ALE maps (Spearman $\rho = -0.507$, two-sided permutation $p = 0.0242$). With known coordinates, $\rho = -0.406$ and $p = 0.0764$. Each of 9,999 permutations reassigned labels jointly to rows and columns of one matrix, preserving dependence among pairs.

#### Overlap after regrouping filtered abstracts

We repeated ALE on the 867 publications eligible for the filtered-abstract analysis, comparing the original ten labels with the 10 groups selected from abstracts after filtering anatomical and result statements. The selected grouping resolves tied modal counts by retaining the middle counts and then choosing the partition with greatest mean text agreement across the grid. This tie rule was amended following author feedback, after the initial smallest-on-ties result and before computing maps for the revised selection. No anatomical outcome was used to choose the revised partition. Both partitions keep exactly the same publications and all aliased source-record coordinates, removing duplicate foci as in the primary analysis. Known-coordinate exclusion left 749 publications. Fits retained the original broad coordinate policy, assumed sample size 20, common 2mm mask and approximate ALE null, with Benjamini–Hochberg correction across group-by-voxel tests within each partition and coordinate analysis.

The matched original categories retained 0 voxels significant in all ten maps with all coordinates and 0 with known coordinates. The selected filtered-abstract grouping gave 3 intersecting voxels with all coordinates ($24\text{mm}^3$), and 0 with known coordinates. The initial smallest-on-ties rule selected 8 groups and gave 6 intersecting voxels ($48\text{mm}^3$), and 1 with known coordinates (Supplementary Table 26). The small all-coordinate intersections lay at the right insular/orbitofrontal border in the Harvard–Oxford cortical atlas. Coordinates, atlas labels and corrected values for every intersecting voxel are saved with the source data.

The selected all-coordinate intersection occupied 0.007% of the combined significant territory. The initial intersection occupied 0.013% and lay inside larger FDR clusters of 42–2,318 voxels. These are intersections of thresholded maps, not a cluster-FWE test of a tiny activation cluster. The filtered-abstract sensitivity follows the primary voxelwise FDR criterion. The additional FWE check is reported below. Such small intersections do not by themselves establish an anatomically meaningful or self-specific core, or show that the location contributes to every self-related process. They qualify the literal zero-intersection statement while retaining very limited shared extent across these alternative partitions. Weak shared contributions could remain undetected. The graded, inventory and on/off models did not use category labels, so their predictive comparison is a separate result.

We also applied the original FWE sensitivity to the categories regrouped from filtered abstracts, with 1,000 random-coordinate datasets per category, a cluster-forming threshold of $p < 0.001$ and six face neighbours. We report cluster-size, cluster-mass and voxelwise correction at 0.05 within each map and at 0.005 to adjust across ten maps. This additional check was specified after the three-voxel FDR intersection was known. With cluster-size FWE correction, 0 voxels were shared by all ten maps with all coordinates and 0 with known coordinates. With the additional correction across ten maps, the corresponding counts were 0 and 0. Of the original three FDR-shared voxels, 0 remained in all ten cluster-size FWE-corrected maps at 0.05 and 0 at 0.005. Supplementary Table 27 gives all corrections and the original full-corpus reference. The FDR maps and their raw intersections remain the primary results.

#### Whether a contribution common to all self-forms could be detected

A contribution can be present in every self-form and still fail to reach significance in one or more of them. We asked how often a requirement of significance in all ten would miss such a contribution.

We simulated a single feature whose probability of being reported was raised in every self-form by construction, and Supplementary Table 8 reports how often it was detected.

Small increases were rarely detected in every self-form, although every simulated self-form contained them. A requirement of significance in all ten maps can therefore miss a weak contribution that all share, which is the limitation stated in the Discussion. The simulation concerns one reported feature tested in ten groups.

##### Specification of shared voxels, clusters and what is displayed

The self-forms whose maps contain a voxel were stored as a code of ten bits. Clusters of shared voxels joined voxels through any of their 26 neighbours, whereas counts of clusters within the map of a single self-form joined voxels through shared faces only. At each level from four to eight self-forms, the three largest combinations were displayed when they held at least ten voxels, a rule that governs display alone.

##### Specification of the simulation

The simulated self-forms had the sizes of the real ones, 146, 139, 135, 124, 87, 79, 58, 52, 46 and 44 publications. The simulated comparison sample had 905 publications, matching the 905 nonempty extraction profiles of the general-imaging comparison sample used for the gradients, before records of the same publication were combined. For a background probability $p_0$ and a common increase $\delta$, each draw generated one comparison count $c \sim \text{Binomial}(905, p_0)$ and an independent count for each self-form, $k_d \sim \text{Binomial}(n_d, p_0 + \delta)$. All ten tests in a draw shared the same comparison count. The two-by-two table for self-form $d$ was $[[k_d, n_d - k_d], [c, 905 - c]]$.

Each self-form was tested by an exact one-sided hypergeometric test. The feature counted as detected in all ten when every p value met the threshold of 0.05, which is the rule for a conjunction [7]. We generated 10,000 draws for each combination of the background probabilities 0.05, 0.10 and 0.20 with the increases 0.02, 0.05, 0.10, 0.20 and 0.40.

#### Supplementary Note 3 | Differences between self-forms are distributed across the brain

*Related main text. Results 2.2.3 and Methods 4.3.3.*

If one region or division carried what distinguishes the self-forms, their differences would be concentrated there. This Note reports how widely each map extends, how well the self-form of a publication can be predicted from the whole brain and from parts of it, and what remains after the most predictive regions are removed.

##### How widely each map extends

Counting the divisions that hold more than 1% of a map's significant voxels, single maps reached eight to thirteen of the 19 divisions, with a median of eleven (main Fig. 5b). A division is a putative cortical resting-state network, the subcortex or the cerebellum, and a map can reach several divisions while occupying little of each. We therefore also measured how many regions of an independent anatomical atlas each map covers, and compared that number with what a single connected territory of the same size would cover.

The anatomical atlas is the FreeSurfer Destrieux `aparc.a2009s` parcellation with `aseg`, which has 167 regions of grey matter, 148 cortical, 17 subcortical and two of cerebellar cortex [8]. Its cortical regions are those outlined on the cortical figures, and coverage was measured in the volume. All 95,846 labelled voxels lie inside the ALE mask and make up 41.95% of it. The atlas and the ALE images share orientation, affine and voxel grid.

We used four rules to decide whether a map covers a region. Under the most lenient, any significant voxel, the maps of the all-coordinate analysis covered 55–100 regions each, with a median of 77.5, and together covered 150. These are 32.9%–59.9% of regions for single maps and 89.8% for all maps together. Requiring at least 20 significant voxels reduced coverage to 22–73 regions each, with a median of 32.5, and to 125 together. Requiring significant voxels to fill 1% or 5% of the eligible volume of a region gave 146 and 131 regions for all maps together. Coverage therefore depends on the rule, and under every rule it extends well beyond a few regions (main Fig. 2l).

In the known-coordinate analysis, the lenient rule gave 50–95 regions for single maps and 146 for all together, the 20-voxel rule gave 17–62 and 119, and the 1% and 5% rules gave 138 and 128 for all together. All of these are numbers of atlas regions. Values for each self-form are indexed with the source data.

Larger maps reach more regions through size alone. To test whether size explains the spread, we generated 10,000 connected territories for each self-form and for all maps together, in both coordinate analyses, each containing as many voxels as the map has within the atlas. The statistic was the number of regions holding at least 20 voxels. All 22 maps covered more regions than their territories after Holm correction, with adjusted p values from 0.00220 to 0.02640 (main Fig. 2m). All maps together covered 125 regions with all coordinates, against a mean of 57.21 for the territories and a central 95% range of 29–85, and 119 regions with known coordinates, against a mean of 52.49 and a range of 24–79. Both uncorrected p values were 1/10,001, and both Holm-adjusted p values were 0.00220. Summaries are indexed with the source data.

#### Predicting self-form from the whole brain and from parts of it

A map can extend widely while one location still carries most of what distinguishes the self-forms. We therefore asked how well the self-form of an unseen publication can be identified from its regional profile, using all regions or a restricted set. Main Figure 5c,d and Supplementary Tables 9 and 10 report the comparisons. The classifier given the whole brain exceeded the classifier given the best division, chosen within the training set, by 6.43 percentage points with all coordinates and 9.75 with known coordinates, both with $q = 0.0002$.

The restricted classifiers were chosen within the training set. Twenty compact candidate sets were defined by k-means clustering of the three-dimensional centres of the parcels, with ten starts and seed 20260906, and each candidate held the 50 parcels closest to one cluster centre. One candidate, and separately one division, was chosen by threefold validation inside each training set. The compact sets are defined by proximity, so their parcels are near one another without necessarily being contiguous or of equal volume.

With a further classifier, we tested whether the best division could be beaten without more predictors. In each fold, a set of regions was sampled uniformly without replacement from all 1,041, as many as the chosen division contains, using seed 20260906 plus 100 times the repeat number plus the fold number. The sets held 66–82 regions with all coordinates and 32–82 with known coordinates. They reached balanced accuracies of 0.190 and 0.195, against 0.162 and 0.139 for the chosen divisions. The differences were 0.0287 and 0.0561, with descriptive 95% intervals of 0.0032–0.0535 and 0.0316–0.0809 that hold the fitted models fixed. The sampled sets match the division in number of regions and are otherwise free to fall anywhere.

We tested separately whether the whole-brain classifier exceeds chance. In the first repeat its balanced accuracy was 0.233, against a mean of 0.100 over 999 shuffles of the labels ($p = 0.001$). Comparisons between classifiers are based on the mean of the two repeats, 0.226.

Reporting characteristics and task and clinical indicators predicted the categories strongly, as expected of labels derived from text, and adding regional profiles did not improve that prediction. The results above therefore concern the information that regional profiles carry about self-forms on their own.

#### Removing the most predictive regions

We removed the most predictive regions to test whether distinguishing information remains elsewhere. The importance of a region was the largest absolute coefficient it received across self-forms in the multinomial logistic regression fitted to the training set, and the classifier was refitted after removal. The 10, 50, 100 or 200 most important regions were removed. For comparison, 50, 100 or 200 regions were removed at random, taken as successive stages of one random ordering in each fold. Removing the 200 most important regions lowered accuracy, relative to the intact classifier, by 0.0055 with all coordinates and 0.0057 with known coordinates, with intervals of −0.0092 to 0.0203 and −0.0110 to 0.0220. Distinguishing information is therefore available beyond the strongest predictors (main Fig. 5e). The intervals include zero and are compatible with a small loss in classification accuracy after the classifier is refitted. The comparisons with sampled sets and after removal are descriptive and add no family of tests. Their scores are indexed with the source data.

#### Specification of coverage and random territories

A division counted for a map when it held more than 1% of the map's significant voxels, among the voxels that the atlas assigns to a division. For all maps together, a division counted when it held a significant voxel from at least one self-form.

The anatomical atlas and the ALE mask share a voxel grid and affine, so the 95,846 labelled voxels needed no resampling. Each random territory started from a voxel drawn uniformly from the 95,840 labelled voxels that form a connected body large enough for every requested size. Six isolated voxels could not start a territory. The territory then grew by repeatedly adding a voxel drawn from those adjacent to it through any of 26 neighbours, until it reached the required number of voxels. Within one draw, territories of different sizes were successive stages of the same growth. The statistic was the number of regions holding at least 20 voxels. Ten thousand draws were made for each map and for all maps together, with Holm correction over the 22 comparisons of both coordinate analyses. The comparison is with a single connected territory of the same size.

#### Specification of the classifiers

The classifiers were multinomial logistic regressions with $C = 0.1$ and weights that balance the self-forms. The classifier for a single region was simpler. It was based on the probability of reporting that region in each self-form, estimated with add-one smoothing and balanced across self-forms. Restricted locations were chosen by threefold validation inside each training set, and the compact candidates and sampled sets are defined above.

Regions were ranked by their largest absolute coefficient across self-forms in the training set. The removed regions were deleted from both training and test profiles before refitting. The eight saved paired comparisons were FDR-corrected within each coordinate analysis. The test against chance, with 999 shuffles, rebuilt the folds for each shuffle, refitted the classifier through one fivefold repeat and scored it against the shuffled labels.

### *Where does the variation come from?*

#### Supplementary Note 4 | Context within a self-form

*Related main text. Results 2.3.1 and Methods 4.4.1.*

If each self-form had one fixed set of regions, its publications would report the same regions, and what a study describes would not predict which regions it reports. This Note documents the variation among the publications of a self-form and the two analyses that predict it from text.

##### Publications of the same self-form report different regions

A division counted for a publication when any of its regions was reported, using the 910 or 784 regional profiles of the two coordinate analyses. Main Figure 5a displays the publications grouped by self-form and ordered by their number of divisions and then by identifier.

##### Differences in text within a self-form

A self-form such as Autobiographical memory groups studies that differ in their tasks, stimuli and participants. We tested whether the differences in text among the publications of a self-form improve prediction of their regions, beyond a comparison model that contains the category and the reporting characteristics, with or without the task and clinical indicators. We subtracted each category's mean training embedding from the embedding of each of its records. The remaining values describe differences in text among publications of the same self-form, while category membership accounts for differences between self-forms.

All four Brier comparisons, from the two comparison models in the two coordinate analyses, survived FDR correction (Supplementary Fig. 2c and Supplementary Table 11). The gains were small. Log loss worsened when text was added to the comparison model without the indicators with all coordinates, and the other three intervals for log loss included zero, so the result depends on the score. On Brier loss, what a study describes predicts, to a small degree, which regions it reports within a self-form. The context of an individual at a given moment was not measured.

##### Abstracts with anatomical and result statements filtered

An abstract can state the findings that are being predicted. We therefore compared the original text with abstracts after filtering anatomical and result statements, termed filtered abstracts, on the same eligible publications. Only 77 abstracts had sections with headings for aims, objectives or methods, and these abstracts were unevenly distributed across self-forms. A fixed filter of sentences was therefore applied to abstracts without such headings. Every title was omitted, and statements of findings, names of brain regions and expressions of statistical results were removed. The rules, and the corrections made to the text alone, were fixed before the filtered abstracts were embedded and before any coordinates were loaded.

The final rule kept 867 of 910 publications, or 95.3%. The numbers in the ten self-forms, in the usual order, were 138, 133, 131, 116, 83, 74, 53, 50, 46 and 43, which is 91.4–100% of each. Seventy-four publications contributed headed sections and 793 contributed filtered sentences, and the median length of the text kept was 80 words. The models that added original text and filtered abstracts were both fitted to these publications within the original folds, and their shared comparison model kept its fit to all training publications. Filtering the text is thereby the only difference between the two. For both, adding text was chosen over the comparison model alone in all ten outer folds.

On these publications, adding the original text improved Brier loss by 0.0000880 (95% interval 0.0000322–0.0001466, Holm-adjusted $p = 0.0082$), and adding the filtered abstracts improved it by 0.0000573 (0.0000367–0.0000780, $p = 0.0003$). The difference between the two in Brier loss was uncertain ($p = 0.2069$). The filtered abstracts had no clear advantage in log loss over the comparison model ($p = 0.5611$), although its log loss was better than that of the original text (Supplementary Table 12).

The gain that remains with filtered abstracts makes it less likely that statements of outcome alone explain the original result. Two cautions apply. The filter can remove information about tasks along with findings, and the categories were still formed from the complete text. The source-data index lists the rules and thresholds of the filter, the normalisation of characters, the decisions for each section and sentence, the text kept, hashes, checks of the comparison model, records of validation and the folds.

#### Specification of the fitting within a self-form

The mean embedding of each category was removed within the relevant training set before the principal axes were found and the values standardised. The values of each record on these axes were then fitted to the regional residuals of a comparison model containing the category and the reporting characteristics, with or without the indicators. The residuals are the differences between the observed reports and the predictions of the comparison model. The penalty of the comparison model was chosen first, and its predictions were identical with and without the added text. The added regression had no intercept. The number of axes, 3, 10 or 30, and the ridge penalty, 1, 10, 100, 1,000 or 10,000, were chosen by validation within the training set, and the comparison model alone was also a candidate. The result was evaluated in two repeats of the fivefold scheme. The four Brier comparisons, from two comparison models in two coordinate analyses, were FDR-corrected together, and the tests of log loss formed a separate family.

#### Specification of the filtered-abstract comparison

Under the fixed rule, we first looked for headed sections of aims, objectives, methods, design, setting or participants, and otherwise applied the recorded filter of sentences. Both routes omitted titles, sections of results and conclusions, explicit statements of outcome, names of brain regions and expressions of statistical results. A sentence was kept only if it had five alphabetic words, and a record only if twenty words remained. The exact expressions and the corrections made to the text are indexed with the source data.

Both versions of the text were embedded with BAAI/bge-large-en-v1.5 and analysed in the original two repeats of five folds, with two folds inside each training set. Only the models that added text were restricted to the eligible records, and the shared comparison model kept its original training publications and preparation. The settings of each added model were chosen independently from 3, 10 or 30 axes, the five penalties from 1 to 10,000, and the comparison model alone. Losses were calculated for each repeat, averaged within publications and then averaged with equal weight for each self-form. The three paired comparisons were evaluated with 10,000 bootstrap samples within self-forms and two-sided tests by reversal of sign, with Holm correction separately for Brier loss and for log loss.

#### Supplementary Note 5 | The aspect of consciousness

*Related main text. Results 2.3.2 and Methods 4.4.2.*

We asked whether each self-form relates to consciousness in one uniform way, or in a way that depends on the aspect of consciousness concerned. The comparison is based on publications that belong to one corpus only, and on the seven final groupings of the companion study [9], each of which concerns an aspect of consciousness. The assignments and names of the groupings were imported before any comparison, and publications in both corpora were removed before any map was calculated. Each map gives the proportion of publications that reported each region (main Fig. 6).

##### The whole table and its individual pairings

Correlating the map of every self-form with the map of every grouping gives a table of ten rows and seven columns, and two statistics summarise it. The first, $H$, is a measure of how much the self-forms differ in their profiles of correspondence across the groupings. The second, $I$, is a measure of the remaining variation between pairings after the mean for each self-form and the mean for each consciousness grouping are subtracted and the overall mean is restored. A self-form whose map resembles every consciousness map more strongly therefore cannot produce a large value of $I$ by that alone. The equations and the shuffling of labels are specified below.

Both statistics exceeded the values obtained from shuffled labels in both coordinate analyses. $H$ was 0.01180 with all coordinates and 0.01114 with known coordinates, and $I$ was 0.00989 and 0.00938, respectively. Each uncorrected p value was 0.0001, and the Holm-adjusted p values were $p = 0.0002$ for $H$ and $p = 0.0002$ for $I$ (main Fig. 6c). By shuffling the labels of self-forms within groups of similar publications, we tested whether the labels are associated with the pattern of the table, given reporting and clinical content.

The correlations ranged from 0.070 to 0.691 with all coordinates and from 0.065 to 0.671 with known coordinates, and the two tables correlated at 0.988. Individual cells were examined with two families of tests, each FDR-corrected over the 70 cells. The first compared each correlation with its values under shuffled labels, and there were 19 departures with all coordinates and 17 with known coordinates. The second compared each remainder, the value left after the means for the self-form and the grouping are subtracted, with its values under shuffled labels. There were 30 and 24 departures, respectively. All 24 found with known coordinates were also found with all coordinates, with the same direction relative to their shuffled means. The two tables of remainders correlated at 0.989.

Supplementary Figure 3 displays both sets of values. Panel a shows each correlation minus its mean under shuffled labels, so a negative value there means that a correlation is lower than its mean under shuffled labels. Panel b shows the remainders themselves, so a negative value there means that the correlation is lower than expected from the means for its self-form and its grouping. Stars mark cells that depart from their own distribution under shuffled labels. A star is a test of neither a correlation against zero nor the difference between two chosen correlations.

##### The illustrative pairings and their resampling

Autobiographical memory and Ownership and agency illustrate the result for the whole table with two well-represented groupings. Autobiographical memory corresponded more closely to Awareness in memory and learning than to Conscious perception and attention, with correlations of 0.636 and 0.363, and Ownership and agency showed the reverse ordering, with correlations of 0.398 and 0.666. Taking each difference in its observed direction gives 0.273 and 0.268, which sum to 0.541. The two self-form maps were based on 139 and 84 publications, and the two consciousness maps on 82 and 113. In the known-coordinate analysis the sum was 0.549, and the corresponding numbers of publications were 113, 79, 75 and 91.

The sum stayed positive in all 2,000 resamples of publications in both coordinate analyses. More directly, both orderings held together in 1,999 resamples of the all-coordinate analysis and in all 2,000 of the known-coordinate analysis. The second count matters because a positive sum can arise without the orderings being opposite. The central 95% ranges of the sum were 0.258–0.642 and 0.258–0.661. The pairings were chosen to illustrate, so these ranges describe stability under resampling and add no further test (main Fig. 6a,b).

Interoception also corresponded to Interoceptive awareness and pain experience, at 0.691 with all coordinates and 0.671 with known coordinates. The seven groupings, in the order C1–C7 of the figure, had 113, 85, 82, 75, 17, 57 and 36 publications with all coordinates and 91, 68, 75, 69, 17, 51 and 30 with known coordinates. The small grouping on bodily self-consciousness is part of the full table and plays no part in the illustration. The summary of the resampling is indexed with the source data.

#### Specification of the two corpora and their maps

Publications in both corpora were identified by their identifiers and by exact title with year, and the 113 found were removed before any map was made. The seven groupings were Conscious perception and attention, Disorders and states of consciousness, Awareness in memory and learning, Affective awareness and fear processing, Bodily self-consciousness and agency, Altered states of consciousness, and Interoceptive awareness and pain experience. Their numbers of publications with all coordinates were 113, 85, 82, 75, 17, 57 and 36. The ten self-forms, in the order of the figure, had 139, 135, 84, 120, 87, 53, 54, 48, 35 and 44.

Each map was the mean of the regional profiles of its publications over all 1,041 regions. Publications with empty profiles were included, two self and three consciousness publications with all coordinates and one and two with known coordinates. Correlations included regions that neither map reported. These maps of reporting frequency are a different measure from ALE convergence.

#### Specification of the statistics of correspondence and the shuffling of labels

Let $R_{ij}$ be the correlation between the map of self-form $i$ and the map of grouping $j$. The first statistic is a measure of how much self-forms differ around the mean of each column,

$$H = \text{mean}_{ij} \left[ (R_{ij} - \bar{R}_{\cdot j})^2 \right] .$$

The second is a measure of the variation that remains after the overall resemblance of each row and each column is removed,

$$E_{ij} = R_{ij} - \bar{R}_{i\cdot} - \bar{R}_{\cdot j} + \bar{R}_{\cdot\cdot}, \qquad I = \text{mean}_{ij}(E_{ij}^2).$$

All means weight the ten self-forms and the seven groupings equally. In each of 9,999 shuffles, the labels of self-forms were shuffled within quarters of the number of reported locations, crossed with a clinical-title indicator that matches the stems `patient`, `disorder`, `depress`, `schizophren`, `autis`, `anorexi` or `clinical`. Each publication kept its whole regional profile, and the number of publications of each self-form in each group was preserved. The consciousness maps stayed fixed, and the self maps, the correlations and both statistics were recalculated. One-sided p values, from the upper tail, were

Holm-corrected across the two statistics within each coordinate analysis (main Fig. 6c). The question is whether the labels of self-forms predict the pattern of pairings within these groups. Subtracting row and column means defines a statistic of departure from additivity. Shuffling self-form labels allows the row and column means to vary.

For individual cells, two-sided p values doubled the smaller tail of the cell's distribution under shuffled labels. The tests of correlations and of remainders were FDR-corrected separately over 70 cells in each coordinate analysis.

##### Specification of the resampling of the illustrative pairings

In each of 2,000 draws, publications were resampled with replacement within each of the ten self-forms and each of the seven groupings, and both sets of maps were rebuilt. For Autobiographical memory we took its correlation with Awareness in memory and learning minus its correlation with Conscious perception and attention, and for Ownership and agency the reverse. We recorded the sum of the two differences and whether both were positive in the same draw, which is the direct check that the orderings stay opposite.

#### Supplementary Note 6 | Systems shared with other functions

*Related main text. Results 2.3.3 and Methods 4.4.3.*

If self-forms draw on systems that serve other functions, the map of each should resemble maps of those functions, and regions reported together in self-related studies should follow the organisation seen across imaging studies in general. This Note takes the two predictions in turn.

##### Correspondence with maps of general functions

We compared the map of each self-form with independent maps of familiar research topics. All 50 official topics of Neurosynth version 7 were kept. A topic is a set of words that recur together, found by latent Dirichlet allocation. Each topic map combined maps made by multilevel kernel density analysis (MKDA), in which each publication marks the voxels within 10 mm of its coordinates. A publication contributed in proportion to how strongly it expressed the topic [10]. Of the 14,371 source publications, 459 matched our corpus by PMID, DOI or normalised title. A further search of titles for self returned 312 publications, 90 of them new. Together these removed 549 publications and left 13,822. The exclusions change which publications contribute to the maps and leave the topics themselves as published. The evidence for each match is indexed with the source data.

The maps of self-forms and of topics were compared on a common grey-matter mask of 142,985 voxels. Some locations are reported often whatever the topic. To reduce resemblance from this source, we adjusted both maps for the mean MKDA map of all the remaining publications by regression before correlating them. Autobiographical memory corresponded to the topic memory retrieval encoding at $r = 0.508$ with all coordinates and 0.495 with known coordinates. Ownership and agency corresponded to action actions observation at 0.541 and 0.540, and to spatial body human at 0.533 and 0.538. Interoception corresponded to pain somatosensory stimulation at 0.392 and 0.381, and to food taste weight at 0.351 and 0.357. The full profiles of adjusted correlations were similar in the two coordinate analyses, correlating at 0.9897–0.9992 across self-forms.

Smooth maps resemble one another to some degree by chance, so each correlation was tested against surrogate maps of matching smoothness. In the all-coordinate analysis, the map of each self-form was phase-randomised 10,000 times and adjusted for intercept and reporting density on the same mask.

FDR correction across all 500 comparisons left 310 significant. Main Figure 2k shows 20 topics, chosen by their official names, and 128 of its 200 cells are significant after correction. Every self-form had between six and 16 significant correspondences among those displayed. The three examples given in the main text each had $p = 0.00010$ and $q = 0.000286$.

Cells that are not significant after correction are left white in the figure, and the source tables hold every correlation. Each self-form's map resembles maps of topics studied without reference to the self. Complete correlations and tests are indexed with the source data.

#### What the gradients are and why ten are shown

A gradient orders regions by how similarly they are reported across publications. The gradients were estimated from the regional profiles alone, without the categories, and they describe the literature as a whole and no individual brain. To describe a gradient, we examined the research of the publications that lie near each of its ends, and the labels we give summarise that research (main Fig. 7a).

Three language models proposed labels from the 40 records at each end of a gradient, their distinctive terms and example titles. The sources of the proposals were hidden when one was chosen, and the names of the categories were withheld. Each final label was checked against all the evidence for its end and kept broad when the research there was mixed. Because the vocabulary for labels had been restricted to the self and experience, the labels describe the gradients in those terms. The evidence and the history of each label are indexed with the source data.

The embedding produced 25 gradients, and the first ten were displayed and tested, the later ones being generally less reproducible. The choice of ten is one of convenience. The number ten also appears as the number of categories and as the number of components in the best graded model, and the three numbers have nothing to do with one another.

#### Gradients shared with imaging research in general

We asked whether the gradients of the self literature also organise imaging research in general. For each self gradient, we took its strongest absolute correlation with any of 25 general-imaging gradients. Searching among many candidates produces large correlations by itself, and smooth maps resemble one another for reasons of space alone. Each surrogate therefore went through the whole search for its best match, so that the test allows for both (main Fig. 7a and Supplementary Fig. 4).

Across the ten gradients, the strongest correlations ranged from 0.342 to 0.818 with all coordinates and from 0.373 to 0.853 with known coordinates. G1–G7 passed both corrected spatial tests with all coordinates, and G1–G4 and G6 passed both in both coordinate analyses. Supplementary Table 13 gives the full estimates.

The first kind of surrogate, from BrainSMASH, was generated from a model of how the difference between two regions grows with the distance between their centres. That relation is called a variogram, and the variograms of the surrogates reproduced the observed ones only in part. We therefore repeated the test with a second kind of surrogate, which randomises the signs of a gradient's coefficients on a set of spatial patterns derived from the distances between regions, known as a Moran eigenbasis. This preserves the variance of the gradient and its spatial autocorrelation as measured by Moran's $I$. The two tests are based on different models of spatial structure on the same data.

Reliability is a separate reason for caution about the later gradients. In the all-coordinate analysis, the mean correlation between matched gradients from the two halves of the corpus fell from 0.878 for G1 to 0.306 for G10 (main Fig. 7c). A gradient can therefore match a general-imaging gradient significantly and still be poorly reproducible, and each should be judged on both counts.

The general-imaging sample was drawn from publications that the self query did not retrieve and was matched to the corpus on year and number of reported locations. It was matched on nothing else, so it may differ in tasks and may contain some self-related research, and excluding unknown spaces changes its composition. Within those limits, the result supports a correspondence between the organisation of the self literature and that of imaging research in general.

#### Positions of self-forms on the gradients

Gradients describe how regions are organised, and they need not separate the self-forms. To test whether they do, each publication was given the mean gradient value of its reported regions as its position. Differences in position between self-forms show whether publications grouped by their text occupy different parts of gradients estimated from regional profiles (main Fig. 7b).

Self-forms differed in position on seven of the ten gradients after FDR correction in each coordinate analysis. These were G1–G3, G5–G6 and G8–G9 with all coordinates and G1–G3, G5–G7 and G9 with known coordinates, six being the same. The seventh and eighth gradients of the known-coordinate analysis were exchanged to align them with the all-coordinate display. Supplementary Table 13 reports every association and its corrected p value, alongside the matches with general-imaging gradients.

Membership of a self-form explained 1.4–21.3% of the variance in position across gradients with all coordinates, which leaves 78.7–98.6% within self-forms. The corresponding ranges with known coordinates were 1.0–22.0% and 78.0–99.0%. The positions of single publications on the first two gradients show how different means coexist with overlapping distributions (Supplementary Fig. 5a–j). The variation within self-forms includes differences of sampling and reporting between publications, and it describes publications and no change within a person.

On G1, Ownership and agency had a mean position of −0.316 and Autobiographical memory a mean of 0.472, in units of the standard deviation across regions. On G2, Body image/identity had a mean of −0.314 and Interoception a mean of 0.267. All four of these means had $q = 0.00222$ with all coordinates and also survived correction with known coordinates. Of the 100 combinations of self-form and gradient, 18 survived with all coordinates and 17 with known coordinates, 13 being the same. Each test compares the mean of a self-form with the means obtained when labels are shuffled among publications of similar reporting and clinical content.

#### Specification of topic maps and spatial testing

The map of a topic was the average of the binary 10 mm MKDA maps of the remaining publications, weighted by each publication's weight on the topic. Publications were excluded when they matched a publication of our corpus, or one of its other identifiers, by PMID, DOI or normalised title, and separately when their title contained self. The topics stayed as published. Correlations were based on the unthresholded ALE $z$ maps of the self-forms on the shared mask of 142,985 voxels. An intercept and the mean binary map of the remaining publications were regressed out of both maps before they were correlated.

For each of the 10,000 surrogates of a self-form's map with all coordinates, the adjusted map was placed in its full three-dimensional grid with zeros outside the mask. Its Fourier phases were replaced by those of real Gaussian noise, which preserves the amplitude spectrum of the grid and with it the smoothness of the map. The surrogate was then sampled on the shared mask and adjusted again for intercept and density, so the spectrum is preserved exactly only before that last step. Two-sided tests compared absolute correlations, with FDR correction over all 500 comparisons of a self-form with a topic. The topics displayed were chosen from their official names before any correlation was inspected.

#### Specification of gradients and positions

The gradients were estimated from 912 self publications and 914 general-imaging publications with all coordinates, and from 785 and 698 with known coordinates. These counts include publications with empty regional profiles, which contribute to the estimate and cannot themselves be placed on a gradient.

The link between two regions was the Pearson correlation, across publications, between their entries in the regional profiles. The diagonal of the matrix of links was set to zero, the smallest value in the matrix was subtracted from every entry, and the diagonal was set to zero again. A correlation that could not be computed would have been replaced by zero, but every region varied across publications. With BrainSpace 0.2.1, we retained the largest 10% of links for each region and measured the similarity between the resulting patterns of links with its normalised-angle kernel. Diffusion mapping of these similarities placed regions along 25 gradients, with $\alpha = 0.5$, a diffusion time of zero and seed 20260906.

The values of each gradient were standardised over all 1,041 regions. The position of a publication was the average value over its reported regions, which gave positions for 910 and 784 publications in the two coordinate analyses. The association between self-form and position was $\eta^2$, the proportion of the variance in position that membership explains. The gradients stayed fixed while labels were shuffled 9,999 times within quarters of the number of reported locations, crossed with the clinical-title indicator of seven stems. The tests of the ten gradients were FDR-corrected within each coordinate analysis. Separate two-sided tests of the mean position of each self-form on each gradient were FDR-corrected over their 100 cells. Labels for the ends of gradients were based on the 40 most extreme records, TF–IDF terms and titles, with the names of categories withheld.

#### Specification of reliability and spatial testing

The corpus was split into two complementary halves 25 times, and the embedding was repeated in each half. The gradients of each half were matched one to one with those of the full solution, by the assignment that maximises absolute correlation, and their signs were aligned. Matching can change the order of gradients and never mixes them. Reliability was the correlation between the matched gradients of the two halves. In the known-coordinate analysis, the seventh and eighth gradients were exchanged for display after matching. Each general-imaging publication was chosen, without replacement, as the nearest remaining candidate to a self record in year and $\log(1 + \text{reported locations})$, each scaled by its standard deviation among the candidates, after the order of the self records had been shuffled with seed zero. The restriction to known coordinates was applied after matching. In the all-coordinate analysis the general-imaging sample resolved to 914 publications. Each self gradient was matched separately, so the same general-imaging gradient could be chosen more than once.

For each of the first ten self gradients, 999 BrainSMASH surrogates were generated from the Euclidean distances between the centres of regions, and the whole search over 25 general-imaging gradients was repeated for each. The second kind of surrogate randomised the signs of coefficients on the eigenvectors of a centred matrix of inverse distances, which preserves variance and Moran's $I$ for those weights, and there were also 999 surrogates with the whole search repeated. Each test was FDR-corrected over the ten matches within each coordinate analysis. Displays have a common range of $\pm 2.5$ standard deviations without thresholding. The cortical surfaces show 1,000 parcels, and the correlations include all 1,041 regions.

### *How do systems combine?*

#### Supplementary Note 7 | The prediction task

*Related main text. Results 2.4 and Methods 4.5.1.*

The last question is how contributions combine to produce the regions that a publication reports. For each unseen publication, a model was given the regions reported in one hemisphere, the observed hemisphere, and predicted those reported in the other, the concealed hemisphere (main Fig. 8a,b).

The principal analysis predicts 987 fragments of cortical parcels, 489 on the left and 498 on the right. A fragment is the part of a parcel that lies at $x \leq -8$ or $x \geq 8$ mm. The two sides are thereby at least 16 mm apart, more than twice the 6 mm margin within which a reported location marks a region, and no parcel contributes to both sides. Trimming in this way keeps 15,922 eligible voxels from 259 parcels that cross into the excluded band. Thirteen parcels that lie entirely within the band are excluded.

All fragments were used, including 28 with fewer than ten voxels, 59 with fewer than twenty and the smallest with one. Fragments have equal weight whatever their volume, which should be kept in mind when reading the estimate for any single fragment. The analysis includes 910 publications with all coordinates and 784 with known coordinates. Four in each report no fragment and stay in their original folds. The publications were fixed by the earlier criterion for the full atlas, before any of these predictions were made.

##### The whole-parcel version

The stricter version is restricted to the parcels that lie wholly outside the band around the midline. It has 728 whole parcels and leaves out 24.4% of the cortical volume of the atlas, including eligible cortex that the fragments keep. It is the more conservative choice and avoids very small fragments.

We treat the fragments as the principal version because they keep all eligible cortex while meeting the same requirement of separation between the hemispheres. Both versions are reported, and the choice changed nothing in the models, the folds, the candidate settings or the rules of inference. The original specification, the sealed outputs and the record of this choice are kept with the source data.

In both versions, cortex within 8 mm of the midline takes no part in the prediction between hemispheres. Trimming parcels into fragments keeps their eligible parts outside that band, and the band itself, which prevents one reported location from marking both hemispheres, is the same in both. Either version can be run in either coordinate analysis, and no other analysis is affected by the trimming. Supplementary Note 10 reports the results for whole parcels.

##### Specification of fragments and whole parcels

Fragments were made by removing the cortical voxels with $-8 < x < 8$ mm and keeping every nonempty remainder of a parcel on either side. Regional profiles, and the count of fragments reported in the observed hemisphere, were rebuilt on the fragments, and the regression on reporting characteristics was refitted within every inner and outer training set. The folds and the rules for choosing settings were unchanged. The whole-parcel version kept 354 parcels on the left and 374 on the right. It excluded 272 cortical parcels holding 32,274 of the 132,032 cortical voxels of the atlas, 15,922 of which lie outside the band. Subcortical and cerebellar regions belong to neither version.

#### Supplementary Note 8 | The three models

*Related main text. Results 2.4 and Methods 4.5.2.*

The inventory model describes a publication by one of a set of complete patterns, and when it is uncertain which pattern applies it averages their predictions. The on/off model builds a pattern from components that are each present or absent, and the graded model lets the strength of each component vary. The on/off model already allows components to be combined, so the decisive comparison is whether graded strengths predict better than combinations alone (main Fig. 8a,b). A number of patterns and a number of components are different measures of size, since three components can produce many combinations and three patterns only three.

The Methods give the equation of each model. In those equations and the ones below, $X_{ij}$ indicates whether publication $i$ reported unit $j$, and $o_{ij}$ is the log odds of reporting given the reporting characteristics alone. A unit is a fragment in the principal analysis and a whole parcel in the stricter version, and units differ in volume. In all three models, the reports for a publication's units are independent once its pattern or strengths and its reporting characteristics are given. Beyond what the reporting characteristics explain, any tendency for units to be reported together therefore arises from the patterns or components.

##### The regression on reporting characteristics

Let $N$ be the number of training publications and $\ell$ the Bernoulli negative log likelihood, which penalises a model for assigning low probability to the observed report or absence of a report. The regression on reporting characteristics minimised

$$N^{-1}\sum_{ij}\ell(X_{ij}, o_{ij}) + \frac{10}{2N}\|B_{\text{slope}}\|_2^2.$$

The first term fits the reports, and the second is a ridge penalty of ten that restrains the coefficients of the characteristics. The characteristics were publication year, log(1 + reported locations), log(1 + analyses), the fraction of coordinates in unknown spaces and log(1 + units reported in the observed hemisphere). The first four are fixed properties of a publication, and the last was calculated from the observed hemisphere for each direction of prediction. Missing values were replaced by training medians, and all five characteristics were standardised within the training set. Intercepts carried no penalty and started at the log odds of the rate of reporting with add-one smoothing, and all other coefficients started at zero. The regression was refitted within every inner and outer training set.

##### The inventory model

For the inventory model, the likelihood was calculated by summing exactly over patterns before taking the negative logarithm. The objective was the mean negative log likelihood plus $\lambda\|A\|_2^2/(2N) - 0.1\sum_k \log\pi_k/N$. The ridge penalty $\lambda$ applied only to the adjustments $A$ of the patterns, and the last term corresponds to a symmetric Dirichlet(1.1) prior on the prevalences. The model had no intercept for each unit beyond $o_{ij}$. Predictions averaged over the patterns by their probabilities.

##### The graded model

In the graded model, $\mathbf{z}_i$ holds the strengths of publication $i$, $W$ the weights of the components on the units and $\mathbf{b}$ the intercepts of the units. The uncertainty about a publication's strengths was approximated by a Gaussian distribution $q_i$ with independent dimensions, known as a diagonal Gaussian variational posterior. The objective was

$$N^{-1} \sum_i \left[ \sum_j \mathrm{E}_{q_i}\, \ell(X_{ij}, o_{ij} + b_j + \mathbf{w}_j^\top \mathbf{z}_i) + \mathrm{KL}\{q_i \,\|\, \mathcal{N}(0, I)\} \right] + \frac{\lambda}{2N} \left( \|W\|_2^2 + \|\mathbf{b}\|_2^2 \right).$$

The expectation averages the fit over the uncertainty in the strengths. The divergence term keeps their distribution close to the standard Gaussian prior and is counted once for each publication. The ridge penalty applied to the weights and the intercepts, and the means and log variances of $q_i$ carried no further penalty. Log variances were bounded to $[-9, 3]$ when the objective was evaluated.

The strengths of an unseen publication were estimated from the observed hemisphere alone. The expected log likelihood was evaluated by Gauss–Hermite quadrature with seven nodes, a numerical method that approximates an average over a Gaussian distribution from a few chosen values, and predicted probabilities were averaged with twenty nodes. Predictions therefore average over the uncertainty in the strengths.

##### The on/off model

For the on/off model, the likelihood was calculated by summing exactly over all $2^R$ combinations of present and absent components before taking the negative logarithm. The objective was the mean negative log likelihood plus $\lambda(\|W\|_2^2 + \|\mathbf{b}\|_2^2)/(2N) - 0.1 \sum_r [\log p_r + \log(1 - p_r)]/N$. The ridge penalty applied to the weights and the intercepts, and the last term corresponds to a Beta(1.1,1.1) prior on each prevalence, which carried no other penalty. In numerical evaluation the prevalences were held within $[10^{-4}, 1 - 10^{-4}]$, and the prior was left untruncated.

#### Supplementary Note 9 | Fitting and choice of settings

*Related main text. Methods 4.5.3.*

Within each version, fragments or whole parcels, the three models had the same reporting characteristics, the same folds and the same targets of prediction. Five outer folds supplied the unseen publications, and the size and penalty of each model were chosen in two inner folds within each training set by Brier loss on the concealed hemisphere. All folds kept similar proportions of the ten self-forms. Size and penalty were chosen separately for each model and for each outer fold, each direction of prediction and each coordinate analysis, from the losses of publications pooled over the two inner folds. The penalties were 1, 10, 100 and 1,000. The inventory model could have 1, 2, 3, 5, 10 or 20 patterns, and the graded and on/off models 1, 2, 3, 5 or 10 components.

The chosen graded model always had ten components and a penalty of 100. The inventory model had three, five, ten or twenty patterns with a penalty of ten, and the on/off model two, three or five components with a penalty of ten or 100. In the analysis on fragments with all coordinates, the on/off model had three components with a penalty of ten in four of its ten fits, three components with a penalty of 100 in four and five components with a penalty of 100 in two. With known coordinates, it had two components in two fits, one with each penalty, three components with a penalty of ten in three fits and with a penalty of 100 in four, and five components with a penalty of 100 in one. Ten was the largest number of components tested, so the number that would be chosen for the graded model without that limit is unknown. Records of the choice in each fold and direction are kept with the source data.

##### Starting values

Starting patterns of the inventory model came from k-means clustering, with one start, of a singular-value decomposition of the training residuals. Each starting pattern was stabilised with five pseudo-observations at the level given by the reporting characteristics, and starting prevalences were set with add-one smoothing.

Starting weights of the graded model were the singular vectors of the residuals from the regression on reporting characteristics, after standardisation, with Gaussian noise of standard deviation 0.025 added. Starting means of the strengths were the corresponding left singular vectors with noise of standard deviation 0.02. Intercepts started at zero and the log variances of training publications at $-0.3$. For unseen publications, means and log variances started at zero.

The first starting point of the on/off model had the singular vectors of the residuals as weights, with noise of standard deviation 0.025, and prevalences of one half. The second had independent random directions scaled to the lengths of the singular vectors, and log odds of prevalence drawn from $\mathcal{N}(0, 0.25^2)$. Intercepts started at zero.

##### Optimisation and convergence

The inventory and graded models were fitted from two starting points with the Adam optimiser [11], using all training publications at each step, in PyTorch 2.14.0 [5]. Fitting was done with 32-bit floating-point numbers and a learning rate of 0.035, and the estimation of strengths for unseen publications with a rate of 0.07. Fitting stopped when three successive changes in the objective over ten steps were each smaller than $10^{-5}\max(|\text{objective}|, 1)$, after at least 90 steps. The base seed was 20260907. Fits within the inner folds had that seed plus 100 times the number of the outer fold plus the number of the inner fold, and fits on the outer training sets had the base seed plus 100 times the number of the fold plus twenty.

In the principal analysis on fragments, fits within the inner folds were allowed 400 steps, and the estimation of strengths within those folds 120 steps. Forty-nine fits of the graded model within inner folds reached that limit before meeting the stopping rule, one of them in a combination of settings that was later chosen from the scores averaged over folds. These are results of choosing settings under a fixed limit, and the fits used for testing are reported next.

All 91 chosen fits of the inventory model and all twenty chosen fits of the graded model on the outer training sets met the stopping rule within 400 steps. The final estimates of strengths for unseen publications converged in 121–161 steps, under a limit of 1,000, and the regressions on reporting characteristics converged in 111–121 steps. No chosen fit on fragments needed more steps. In the whole-parcel version, one chosen fit of the known-coordinate analysis needed an allowance of 800 steps, with the choice of model, the log odds from reporting characteristics, the publications and the seeds unchanged.

The on/off model was fitted with a learning rate that started at 0.035, and the norm of the gradient was limited to 100. The rate was divided by five whenever ten successive checks, ten steps apart, showed no relative improvement of $10^{-5}$ in the best objective, down to a minimum of $3.5 \times 10^{-6}$. This procedure was fixed for the analysis on fragments before any evaluation. The limit of 1,600 steps could be raised to 3,200 and then 6,400 only after a numerical failure. All 1,680 starts of the 840 fits on fragments converged within 1,600 steps, the longest taking 1,441.

#### Checks of the implementation and the saved outputs

For the on/off model, we used independent enumeration of combinations and calculations at higher precision to check the likelihoods, the probabilities of combinations given the observed hemisphere, the gradients, the derivatives with respect to the prevalences, the moments of the prior and the symmetries among components. The sums over combinations for unseen publications were also checked by an independent calculation at higher precision. The on/off model with one component was checked against the equivalent inventory model with two patterns. Changing the reports of the concealed hemisphere left the predictions unchanged, as it should. Quadrature at higher precision changed the likelihood and the predictions of the graded model negligibly.

#### Recovery simulations

A result in favour of the graded model is more convincing if the same procedure can favour a discrete model when a discrete model generated the data. We therefore simulated data from known models and asked which model predicted new observations better. The simulations are a test of the procedure for the generators specified here.

The first simulation set the inventory model against the graded model. The generators were a fitted inventory model with three patterns and a fitted graded model with three components, and each produced twenty datasets with the observed number of publications and known values of the log odds from reporting characteristics. The models fitted to these data had three patterns or three components and a penalty of one. The simulation was run on the 728 whole parcels, with the 354 on the left observed and the 374 on the right predicted, and each dataset had 682 training and 228 test profiles. Fitting allowed 400 steps, with 120 steps for estimating the strengths of unseen profiles and two starting points for each model. Brier loss and log loss both favoured the model that had generated the data in all twenty datasets from each generator, for which the exact 95% interval is 0.832–1.000. The mean advantage of the model that had generated the data was 0.000560 in Brier loss and 0.002247 in log loss for data from the inventory model, and 0.002547 and 0.009317 for data from the graded model. The simulation did not repeat the nested choice of settings or vary the number of patterns.

The second simulation set the on/off model against the graded model. Each generator produced ten datasets with three components, 48 regions, 400 training observations and 200 independent test observations. The two generators shared weights and log odds from reporting characteristics, and their strengths had matched means and variances. In each hemisphere, eight regions were mainly associated with each component, with equal numbers of main weights at $+1.1$ and $-1.1$ and Gaussian weights of standard deviation 0.15 on the other components. The log odds from reporting characteristics were generated separately and treated as known. The prevalences of the on/off components were 0.25, 0.50 and 0.75. Each model was fitted to 300 training observations, given a penalty of one or ten chosen on the other 100, and was refitted to all 400 before predicting in both directions. Both scores favoured the model that had generated the data in all ten datasets from each generator.

All 240 fits from their starting points and all 120 estimates of strengths for unseen profiles converged, in at most 321 and 121 steps.

#### Supplementary Note 10 | Scoring

*Related main text. Results 2.4 and Methods 4.5.4.*

The three models were scored on the regions reported in the concealed hemisphere, with the weights given in Supplementary Method 4. Predicted probabilities were bounded to $[10^{-6}, 1 - 10^{-6}]$ before scoring, which is a separate bound from that of the text models.

##### The graded model compared with the inventory and on/off models

The graded model predicted better than the inventory model with its chosen number of patterns in both coordinate analyses. The gains in Brier loss were 0.001998 with all coordinates ($q = 0.0001$) and 0.001916 with known coordinates ($q = 0.0001$), and both comparisons held for log loss. The p values for Brier loss were FDR-corrected over five comparisons with the inventory model within each coordinate analysis. Those for log loss are uncorrected secondary results and took no part in that decision. Supplementary Tables 14 and 15 report the scores and intervals.

The graded model also predicted better than the on/off model with its chosen number of components, in both coordinate analyses and on both scores, with Holm-adjusted $p = 0.0004$ across those four comparisons (main Fig. 8b). In a further comparison, the on/off model was fixed at ten components, which allows all 1,024 combinations, and only its penalty was chosen. Both scores still favoured the graded model, with positive descriptive intervals (Supplementary Table 16). The advantage therefore comes from graded strength, over and above the freedom to combine components.

##### Separating prediction of the pattern from prediction of the amount reported

A model can score better by predicting how much a publication reports or by predicting which regions it reports. To tell these apart, the Brier loss was separated into two parts for each publication and each direction of prediction. One is the squared difference between the mean predicted probability and the mean of the observed reports. The other is the mean squared difference between the predicted and observed patterns after each has had its own mean subtracted. The two parts sum exactly to the loss. The same paired resamples of publications give descriptive intervals for each part, and no further tests were made (main Fig. 8d and Supplementary Table 17).

On fragments, the gain in the pattern was close to the whole gain over both the inventory model and the on/off model, and every interval for the gain in the amount reported included zero. The two parts are signed and add up to the whole, so the share due to the pattern can slightly exceed 100% when a small disadvantage in the amount reported offsets a larger advantage in the pattern. Main Figure 8c shows the gain of the graded model over the on/off model for each fragment, without tests at individual fragments.

##### What the observed hemisphere contributes

In a second analysis, we asked how much the observed hemisphere contributes beyond the reporting characteristics. For the inventory and graded models with their chosen settings, we compared predictions based on the observed hemisphere with predictions based only on the reporting characteristics. The latter averaged over the patterns or strengths expected before observing the hemisphere's regional pattern. Both came from the same fitted model and the same reporting characteristics. The graded model gained from the observed hemisphere, by 0.003430 in Brier loss with all coordinates and 0.003452 with known coordinates. The gains lay mostly in the pattern, and they stayed positive among publications with a single analysis and among publications reporting in both hemispheres (Supplementary Table 19).

#### Scoring in restricted sets of publications and by self-form

These checks reuse the predictions of the principal analysis on fragments and score them in restricted sets of publications, to see whether particular publications drive the result. Scoring was based on the existing fitted models, whose training sets could include publications left out of scoring.

Supplementary Table 18 gives the scores for publications with a single extracted analysis, publications reporting in both hemispheres, and publications with both properties. The first restriction avoids the pooling of several analyses within a publication, and the second excludes publications whose concealed hemisphere is empty. A single extracted analysis is a property of the extraction and does not show that the publication studied a single psychological operation.

The word self, standing alone, appears in the title or abstract of 720 publications and in the title of 417, and all ten self-forms remain represented among them. The advantage of the graded model over both other models, on both scores and in both sets, survived Holm correction over the eight tests, with $p = 0.0008$ in each (Supplementary Table 20), and the mean for each fold and direction also favoured the graded model. Requiring the word changes the mix of publications, most of all for Interoception, of which 22 of 79 publications remain in the title-or-abstract set, compared with 86 of 87 for Trait evaluation. It leaves eligibility and categories as they were.

The word also occurs in methodological compounds such as self-paced and self-report. Discounting those uses, under the rules given below, removed six publications of Interoception from the title-or-abstract set and one of Negative self-reference from the title set, leaving 714 and 416. In the 714 publications, the gains in Brier loss were 0.001742 over the inventory model, with a descriptive interval of 0.001404–0.002093, and 0.001879 over the on/off model, with an interval of 0.001569–0.002203. In the title set the gains were 0.001373 and 0.001549, also with positive intervals, and the intervals for log loss were also positive.

Finally, the estimate for every one of the ten self-forms favoured the graded model over both other models on both scores, although some intervals for Face/voice recognition included zero. Weighting the self-forms equally gave gains in Brier loss of 0.001922 over the inventory model (interval 0.001598–0.002270) and 0.002050 over the on/off model (0.001748–0.002359), close to the gains of 0.001998 and 0.002105 obtained when publications are weighted equally (Supplementary Table 21). The larger self-forms therefore do not drive the result. The estimates and intervals for single self-forms are descriptive. Identifiers of publications, scores and hashes are kept with the principal outputs, and the lists of publications that mention self are in the source-data index.

#### The whole-parcel version

On whole parcels, the graded model still predicted better than both other models. Over the inventory model, the gains in Brier loss were 0.001882 with all coordinates (interval 0.001523–0.002245, $q = 0.0001$) and 0.002090 with known coordinates (0.001681–0.002517, $q = 0.0001$). Over the on/off model, they were 0.002491 (0.002118–0.002876, Holm-adjusted $p = 0.0004$) and 0.002589 (0.002185–0.003006, $p = 0.0004$), and both comparisons held for log loss. The advantage of the graded model therefore does not depend on the use of fragments (Supplementary Tables 14 and 16). The two versions predict different sets of targets, so their losses cannot be compared with each other directly.

#### Specification of restricted scoring

The sets with a single analysis, with reports in both hemispheres and with both were scored from the existing predictions without refitting. Supplementary Note 1 defines when a publication mentions self. The title-or-abstract set and the title set held 720 and 417 publications of the all-coordinate analysis. Their two scores and two comparisons were based on 10,000 paired bootstrap samples of publications and reversals of sign, with Holm correction over the eight comparisons.

The rule for methodological uses discounted self-paced and self-pacing, the variants of self-report, and self-rated and self-rating. Self-administered was discounted only directly before questionnaire, survey, scale or inventory. Any other use kept the publication in the set. The resulting 714 and 416 publications were scored from the same predictions with 10,000 bootstrap samples and without new tests. The gains for single self-forms, and the gain with self-forms weighted equally, were based on 10,000 bootstrap samples within self-forms, and in the latter each self-form carried one tenth of the weight.

### Supplementary Note 11 | Illustrating the fitted configuration space

*Related main text. Results 2.4 and Methods 4.5.5.*

The comparison of models scores predictions of unseen reports. The displays of the configuration space serve a different purpose, which is to illustrate how the regional patterns predicted by the fitted graded model change as the strengths of its components vary. The positions of publications and the conversion from strengths to patterns come from the data. The paths, their directions and the spacing of points along them were chosen by us for illustration (main Fig. 8e–h).

#### Paths through the graded model's configuration space

All displays are based on one graded model, with ten components and a penalty of 100, from the first fold of left-to-right prediction in the all-coordinate analysis on fragments. A principal component analysis of the estimated strengths of its 728 training publications gives the three axes of the display, which hold 40.8% of the variation in strengths. The model predicts 987 fragments and keeps all ten dimensions, of which the display draws three combinations.

Paths are cubic Bézier curves, smooth curves steered by four control points, drawn through all ten dimensions with the mean strengths of self-forms as control points. The weights of the control points are nonnegative and sum to one, so every point on a curve is a weighted average of those control points. The main curve starts at Interoception and ends at Autobiographical memory, and is steered between them by Ownership and agency and Self-appraisal. Two alternatives are steered by Body image/identity and Face/voice recognition, or by Monitoring and insight and Negative self-reference (main Fig. 8g).

Each curve is converted into regional patterns at five equally spaced points. At each point, the model is evaluated at that set of strengths with the log odds from reporting characteristics held at their mean in the training set. The values are probabilities that each fragment is reported, evaluated at an exact position, without the averaging over uncertainty that is used for unseen publications. Curves with the same end points therefore have the same patterns at their ends and can differ between them.

The maps have a scale of probability from 0 to 0.30, with larger values given the top colour and labelled 0.30+, six views of the cortex and the sampling described in Supplementary Method 2. The points sampled, their regional probabilities and the identity of the fitted model are indexed with the source data.

##### Converting strengths into patterns

The five points of a path were converted into patterns with the fitted weights and one common value of the log odds from reporting characteristics,

$$P_j(t) = \text{logit}^{-1}\left(\bar{o}_j + b_j + \mathbf{w}_j^{\mathsf{T}}\mathbf{z}(t)\right).$$

Here $\bar{o}_j$ is the mean of the fitted log odds for fragment $j$ in the training set, held fixed along the path, and $\mathbf{z}(t)$ is the set of strengths at point $t$. The fragments are the same at every point.

### *All analyses*

#### Supplementary Note 12 | The known-coordinate analysis

*Related main text. Methods 4.2.2.*

In the primary, all-coordinate analysis, we transformed coordinates stated to be in Talairach space and treated unknown or other spaces as MNI. The known-coordinate analysis excludes the unknown or other spaces. It keeps 785 of the 911 ALE maps, 784 of the 910 regional profiles, and 686 self-only and 401 consciousness-only publications in place of 799 and 465. The grouping of text still includes 915 records, and the analysis of filtered abstracts has its own matched sample of 867 publications. This Note summarises what changed, under the three questions of the main text, and the Notes above give the numbers.

##### Is there a core?

In comparisons of whole regional profiles, 25 pairs of self-forms differed after correction with all coordinates and 21 with known coordinates, with 21 found in both (Supplementary Table 4). The overall gain from knowing the self-form over a pooled pattern stayed positive. Individual self-forms were less uniform. Trait evaluation was predicted slightly worse by its own pattern than by the pooled pattern, a difference that was uncertain with all coordinates and survived correction with known coordinates. A continuous description of text did not predict better on Brier loss than the reporting characteristics with the task and clinical indicators, in either coordinate analysis. No voxel was significant in nine or ten of the original maps in either coordinate analysis, and the advantage of the whole brain over the best division in classification stayed positive (Supplementary Notes 1–3).

##### Where does the variation come from?

Differences in text within a self-form improved Brier loss in all four corrected comparisons, whereas log loss did not improve uniformly. Outcome-reduced text was analysed only with all coordinates (Supplementary Note 4).

Both tests of the whole table of correspondence with consciousness held in both coordinate analyses.

The two coordinate analyses gave similar profiles of correspondence with maps of general functions. Tests against surrogate maps were run only with all coordinates. Two questions were asked of the gradients, whether they match gradients of imaging research in general and whether self-forms differ in position on them. G1–G7 matched general-imaging gradients under both spatial tests with all coordinates, and G1–G4 and G6 did so in both coordinate analyses. Self-forms differed on seven gradients in each coordinate analysis, six of them the same (Supplementary Note 6).

#### How do systems combine?

The principal comparison on fragments favoured the graded model over both the inventory model and the on/off model in both coordinate analyses, and both comparisons held for log loss. P values for the Brier comparisons with the inventory model were corrected within their family, and those for log loss are uncorrected secondary results. P values for the comparisons with the on/off model were corrected across both coordinate analyses and both scores. The whole-parcel version kept both advantages (Supplementary Note 10).

#### Supplementary Tables

**Supplementary Table 1**: **The corpus by self-form.** Records are the entries of title and abstract used for the grouping of text. ALE publications are the publications with a nonempty modelled-activation map. Locations are counted after the analyses of each publication are combined and duplicates removed, so their total is smaller than the total before combination in main Fig. 1. Extent is the share of the brain mask covered by the significant voxels of each map. Supplementary Method 2 explains these counts. Self-forms are in the order of the main figures.

| Letter | Self-form | Full name of category | Records | ALE publications | Locations | Extent (%) |
|---|---|---|---|---|---|---|
| a | Autobiographical memory | Autobiographical memory retrieval | 146 | 146 | 7,141 | 7.6 |
| b | Negative self-reference | Negative self-referential processing in depression | 141 | 139 | 3,043 | 3.0 |
| c | Ownership and agency | Body ownership and agency | 136 | 135 | 4,245 | 5.2 |
| d | Self-appraisal | Self-appraisal and decision making | 125 | 125 | 4,289 | 5.3 |
| e | Trait evaluation | Trait self-evaluation and social feedback | 87 | 87 | 2,605 | 3.1 |
| f | Interoception | Interoceptive awareness | 80 | 79 | 2,516 | 2.0 |
| g | Monitoring and insight | Self-monitoring and insight in schizophrenia | 58 | 58 | 1,655 | 1.6 |
| h | Self–other distinction | Self-other distinction in empathy | 52 | 52 | 1,625 | 2.2 |
| i | Face/voice recognition | Face and voice self-recognition | 46 | 46 | 1,105 | 1.2 |
| j | Body image/identity | Body image and self-identity | 44 | 44 | 1,248 | 1.2 |
| **Total** | | | **915** | **911** | **29,472** | — |

**Supplementary Table 2**: **ALE maps under other assumptions about coordinates and spread.** Rows for spread vary the uncertainty around each reported location by assigning an assumed sample size of 10, 20, 40 or 80 to every analysis. Other rows restrict the coordinate spaces or the types of analysis. The Dice coefficient of two maps is twice the volume they share divided by the sum of their volumes, and the table gives its mean over pairs of maps. Union is the share of the mask covered by at least one map. All rows analyse the same corpus. Supplementary Method 2 describes these analyses.

| Analysis | Maps | Locations | Mean Dice | Union (%) | Most maps sharing a voxel | Voxels in all ten |
|---|---|---|---|---|---|---|
| Assumed sample of 10 | 911 | 29,472 | 0.175 | 22.4 | 9 | 0 |
| Assumed sample of 20 | 911 | 29,472 | 0.151 | 19.0 | 8 | 0 |
| Assumed sample of 40 | 911 | 29,472 | 0.139 | 17.3 | 8 | 0 |
| Assumed sample of 80 | 911 | 29,472 | 0.132 | 16.3 | 8 | 0 |
| Known coordinates, assumed sample of 20 | 785 | 25,770 | 0.136 | 16.8 | 8 | 0 |

| Analysis | Maps | Locations | Mean Dice | Union (%) | Most maps sharing a voxel | Voxels in all ten |
|---|---|---|---|---|---|---|
| Other types of analysis removed, assumed sample of 20 | 885 | 28,601 | 0.145 | 18.6 | 8 | 0 |

**Supplementary Table 3**: **Maps of self-forms under correction of the family-wise error rate.** Tests of cluster size and cluster mass form clusters at a p value of 0.001. The voxelwise test evaluates individual voxels directly. The value of 0.005 for each map adjusts across the ten maps. Extents and overlaps are calculated after each correction and can differ from those of the primary maps, which were FDR-corrected together. The table shows how far the findings depend on the method of correction.

| Correction | $\alpha$ | Mean Dice | Most maps sharing a voxel | Self-forms detected | Voxels in all ten |
|---|---|---|---|---|---|
| Cluster size | 0.050 | 0.126 | 7 | 10 | 0 |
| Cluster size | 0.005 | 0.117 | 6 | 10 | 0 |
| Cluster mass | 0.050 | 0.121 | 7 | 10 | 0 |
| Cluster mass | 0.005 | 0.114 | 7 | 10 | 0 |
| Voxelwise | 0.050 | 0.061 | 5 | 10 | 0 |
| Voxelwise | 0.005 | 0.047 | 5 | 9 | 0 |

**Supplementary Table 4**: **Comparisons of regional profiles for every pair of self-forms.** Each test compares the whole regional profiles of two self-forms within groups of publications of similar reporting and clinical content. P values are Holm-corrected over the 45 pairs within each coordinate analysis. The effect is the root mean square of the adjusted differences in the probability of reporting each region, in percentage points, and it summarises the whole profile. Supplementary Figure 1 shows the effects and corrected tests with all coordinates.

| Pair of self-forms | Effect (all coordinates) | $p_{\mathrm{Holm}}$ (all) | Effect (known coordinates) | $p_{\mathrm{Holm}}$ (known) |
|---|---|---|---|---|
| a vs b | 5.50 | 0.1188 | 5.89 | 0.8844 |
| a vs c | 9.91 | 0.0045 | 10.07 | 0.0045 |
| a vs d | 4.86 | 0.0560 | 5.22 | 0.0946 |
| a vs e | 4.90 | 0.2064 | 5.58 | 0.0816 |
| a vs f | 9.00 | 0.0045 | 9.21 | 0.0045 |
| a vs g | 8.79 | 1.0000 | 10.34 | 1.0000 |
| a vs h | 10.38 | 0.0045 | 10.69 | 0.0045 |
| a vs i | 7.41 | 0.0045 | 7.82 | 0.0045 |
| a vs j | 6.87 | 0.3874 | 7.17 | 1.0000 |
| b vs c | 7.89 | 0.0045 | 8.47 | 0.0045 |
| b vs d | 5.12 | 0.5304 | 5.69 | 0.6860 |
| b vs e | 5.07 | 1.0000 | 5.82 | 1.0000 |
| b vs f | 5.74 | 0.0045 | 6.26 | 0.0045 |
| b vs g | 4.70 | 1.0000 | 5.21 | 0.6860 |
| b vs h | 5.94 | 0.0184 | 5.93 | 0.2635 |

| Pair of self-forms | Effect (all coordinates) | $p_{\text{Holm}}$ (all) | Effect (known coordinates) | $p_{\text{Holm}}$ (known) |
|---|---|---|---|---|
| b vs i | 6.69 | 0.0308 | 7.24 | 0.2261 |
| b vs j | 5.79 | 0.1309 | 6.52 | 0.1176 |
| c vs d | 8.26 | 0.0045 | 8.37 | 0.0045 |
| c vs e | 8.26 | 0.0045 | 8.01 | 0.0045 |
| c vs f | 6.93 | 0.0045 | 7.03 | 0.0045 |
| c vs g | 9.90 | 0.0045 | 10.41 | 0.0084 |
| c vs h | 7.57 | 0.0045 | 7.37 | 0.0045 |
| c vs i | 6.89 | 0.0045 | 6.83 | 0.0045 |
| c vs j | 7.85 | 0.0045 | 8.10 | 0.0045 |
| d vs e | 3.98 | 1.0000 | 4.40 | 1.0000 |
| d vs f | 7.21 | 0.0045 | 7.18 | 0.0045 |
| d vs g | 7.80 | 1.0000 | 8.46 | 1.0000 |
| d vs h | 8.19 | 0.0045 | 8.26 | 0.0045 |
| d vs i | 6.74 | 0.0045 | 6.99 | 0.0208 |
| d vs j | 6.11 | 1.0000 | 6.50 | 1.0000 |
| e vs f | 7.77 | 0.0045 | 7.67 | 0.0045 |
| e vs g | 8.58 | 1.0000 | 9.52 | 1.0000 |
| e vs h | 8.84 | 0.0045 | 8.53 | 0.0045 |
| e vs i | 6.97 | 0.0072 | 7.29 | 0.0208 |
| e vs j | 6.15 | 1.0000 | 6.62 | 1.0000 |
| f vs g | 6.40 | 0.5304 | 6.75 | 0.5040 |
| f vs h | 5.96 | 0.2460 | 5.88 | 1.0000 |
| f vs i | 7.00 | 0.0560 | 7.15 | 0.2261 |
| f vs j | 7.52 | 0.0045 | 7.96 | 0.0108 |
| g vs h | 6.54 | 1.0000 | 6.92 | 1.0000 |
| g vs i | 9.45 | 1.0000 | 10.36 | 1.0000 |
| g vs j | 7.90 | 0.0052 | 8.85 | 0.0045 |
| h vs i | 8.15 | 0.0052 | 8.07 | 0.0816 |
| h vs j | 7.47 | 0.0462 | 7.70 | 0.1860 |
| i vs j | 7.55 | 0.2842 | 8.12 | 0.4800 |

**Supplementary Table 5**: **Prediction from the pattern of each self-form compared with a pooled pattern.** Positive differences in Brier loss mean that the pattern of the self-form predicts the reported regions better than the pooled pattern. The ten tests are FDR-corrected within each coordinate analysis, and intervals hold the fitted models fixed. Supplementary Note 1 reports the further comparison models that address the different sizes of the self-forms. The predicted probabilities are a separate measure from thresholded ALE maps. Gains and intervals are in units of $10^{-3}$.

| Self-form | Coordinates | $n$ | $\Delta$ Brier | 95% interval | $q$ |
|---|---|---|---|---|---|
| Autobiographical memory | All | 146 | 3.113 | [1.285, 4.976] | 0.006 |
| Negative self-reference | All | 139 | 0.531 | [-0.097, 1.130] | 0.133 |
| Ownership and agency | All | 135 | 2.204 | [1.474, 2.946] | 0.001 |
| Self-appraisal | All | 124 | -0.441 | [-0.839, -0.041] | 0.077 |
| Trait evaluation | All | 87 | -0.405 | [-0.738, -0.067] | 0.072 |
| Interoception | All | 79 | 0.642 | [0.041, 1.231] | 0.077 |
| Monitoring and insight | All | 58 | 0.004 | [-0.693, 0.642] | 0.990 |

| Self-form | Coordinates | $n$ | Δ Brier | 95% interval | $q$ |
|---|---|---|---|---|---|
| Self–other distinction | All | 52 | 0.079 | [-0.736, 0.862] | 0.990 |
| Face/voice recognition | All | 46 | -0.008 | [-0.635, 0.591] | 0.990 |
| Body image/identity | All | 44 | -0.641 | [-1.350, -0.006] | 0.117 |
| Autobiographical memory | Known | 120 | 3.756 | [1.457, 6.148] | 0.006 |
| Negative self-reference | Known | 106 | 0.437 | [-0.266, 1.082] | 0.367 |
| Ownership and agency | Known | 126 | 2.119 | [1.374, 2.858] | 0.001 |
| Self-appraisal | Known | 108 | -0.201 | [-0.652, 0.270] | 0.498 |
| Trait evaluation | Known | 73 | -0.695 | [-1.045, -0.344] | 0.002 |
| Interoception | Known | 70 | 0.837 | [0.044, 1.584] | 0.101 |
| Monitoring and insight | Known | 52 | 0.081 | [-0.624, 0.732] | 0.821 |
| Self–other distinction | Known | 48 | -0.332 | [-1.092, 0.394] | 0.498 |
| Face/voice recognition | Known | 41 | 0.135 | [-0.559, 0.806] | 0.782 |
| Body image/identity | Known | 40 | -0.595 | [-1.334, 0.094] | 0.221 |

**Supplementary Table 6**: **Prediction of reported regions from reporting characteristics and descriptions of studies.** Lower Brier loss or log loss means better prediction. Losses here are averaged over publications. The paired comparisons in Supplementary Table 7 average gains within each self-form and then equally across self-forms, so the difference between two losses in this table need not equal a gain in that one. Learned groups and continuous descriptions of text were fitted within the training set. These models are separate from the inventory, on/off and graded models that predict one hemisphere from the other.

| Model | Brier all | Brier known | Log loss all | Log loss known |
|---|---|---|---|---|
| Single shared pattern | 0.095411 | 0.096648 | 0.3344 | 0.3376 |
| Reporting characteristics | 0.089280 | 0.090584 | 0.3159 | 0.3196 |
| Characteristics and indicators | 0.089520 | 0.090968 | 0.3223 | 0.3278 |
| Characteristics and ten categories | 0.088754 | 0.090090 | 0.3109 | 0.3149 |
| Characteristics, indicators and ten categories | 0.089374 | 0.090827 | 0.3222 | 0.3277 |
| Characteristics, indicators and three learned groups | 0.089470 | 0.090953 | 0.3224 | 0.3283 |
| Characteristics, indicators and ten learned groups | 0.089357 | 0.090838 | 0.3222 | 0.3277 |
| Characteristics, indicators and continuous description of text | 0.090164 | 0.091726 | 0.3139 | 0.3193 |
| Reduced-rank description, rank three | 0.089300 | 0.090788 | 0.3245 | 0.3299 |
| Reduced-rank description, rank ten | 0.089760 | 0.091142 | 0.3269 | 0.3372 |

**Supplementary Table 7**: **What the categories and finer descriptions of studies add to prediction.** Positive differences in Brier loss favour the model named first. Gains are averaged within each self-form and then equally across the ten. Intervals resample publications and hold the fitted models fixed, and FDR correction covers the seven comparisons within each coordinate analysis. The comparison of ten groups with three uses groups of text learned within each training set, before the regions of unseen publications are predicted. Gains and intervals are in units of $10^{-3}$.

| Coordinates | Comparison | Gain | 95% interval | $q$ |
|---|---|---|---|---|
| All | Ten categories added to characteristics and indicators | 0.151 | [0.087, 0.213] | 0.0002 |

| Coordinates | Comparison | Gain | 95% interval | $q$ |
|---|---|---|---|---|
| Known | Ten categories added to characteristics and indicators | 0.138 | [0.074, 0.202] | 0.0002 |
| All | Ten categories and indicators added to characteristics | -0.185 | [-0.417, 0.058] | 0.1364 |
| Known | Ten categories and indicators added to characteristics | -0.391 | [-0.663, -0.114] | 0.0072 |
| All | Ten categories added to characteristics | 0.447 | [0.330, 0.567] | 0.0002 |
| Known | Ten categories added to characteristics | 0.394 | [0.262, 0.528] | 0.0002 |
| All | Ten learned groups against three | 0.121 | [0.065, 0.177] | 0.0002 |
| Known | Ten learned groups against three | 0.115 | [0.061, 0.170] | 0.0002 |
| All | Continuous description against characteristics and indicators | -0.428 | [-0.735, -0.131] | 0.0090 |
| Known | Continuous description against characteristics and indicators | -0.466 | [-0.831, -0.105] | 0.0145 |
| All | Continuous description against ten categories | -0.578 | [-0.879, -0.287] | 0.0003 |
| Known | Continuous description against ten categories | -0.605 | [-0.964, -0.251] | 0.0013 |
| All | Reduced rank ten against rank three | -0.373 | [-0.512, -0.247] | 0.0002 |
| Known | Reduced rank ten against rank three | -0.333 | [-0.429, -0.237] | 0.0002 |

**Supplementary Table 8**: **Detection of a feature shared by every simulated self-form.** The probability of reporting the feature is raised in every self-form by construction, from the stated background probability by the stated common increase. The table reports how often the tests of all ten self-forms detect the increase, at the sizes of the real self-forms. A small increase can be missed in one or more self-forms although all share it. The simulation concerns one reported feature, as Supplementary Note 2 describes.

| Background probability | Common increase | Detected in all ten | Mean self-forms detected |
|---|---|---|---|
| 0.05 | 0.02 | 0.0000 | 1.54 |
| 0.05 | 0.05 | 0.0010 | 4.75 |
| 0.05 | 0.10 | 0.2190 | 8.55 |
| 0.05 | 0.20 | 0.9454 | 9.94 |
| 0.05 | 0.40 | 1.0000 | 10.00 |
| 0.10 | 0.02 | 0.0000 | 1.15 |
| 0.10 | 0.05 | 0.0004 | 3.45 |
| 0.10 | 0.10 | 0.0649 | 7.42 |
| 0.10 | 0.20 | 0.8394 | 9.82 |
| 0.10 | 0.40 | 0.9999 | 10.00 |
| 0.20 | 0.02 | 0.0000 | 0.92 |
| 0.20 | 0.05 | 0.0000 | 2.47 |
| 0.20 | 0.10 | 0.0130 | 6.09 |
| 0.20 | 0.20 | 0.6237 | 9.53 |
| 0.20 | 0.40 | 0.9999 | 10.00 |

**Supplementary Table 9**: **Classification of self-forms from whole and restricted regional profiles.** Balanced accuracy gives the self-forms equal weight, and chance is 0.10. The restricted regions and the best division were chosen within the training set. A division is a cortical network, the subcortex or the cerebellum. The sampled set has as many regions as the chosen division, drawn uniformly from the whole atlas. Classifiers after removal were refitted without the stated regions. All scores are for unseen publications. Supplementary Table 10 gives the paired differences.

| Classifier | All coordinates | Known coordinates |
|---|---|---|
| Whole brain | 0.226 | 0.236 |
| Best single region | 0.126 | 0.131 |
| Best division | 0.162 | 0.139 |
| 50 nearby regions | 0.152 | 0.156 |
| Sampled set as large as the best division | 0.190 | 0.195 |
| Characteristics and indicators | 0.417 | 0.420 |
| Whole brain, characteristics and indicators | 0.358 | 0.395 |
| 10 most predictive regions removed | 0.218 | 0.239 |
| 50 most predictive regions removed | 0.218 | 0.239 |
| 50 random regions removed | 0.226 | 0.235 |
| 100 most predictive regions removed | 0.223 | 0.233 |
| 100 random regions removed | 0.224 | 0.236 |
| 200 most predictive regions removed | 0.220 | 0.231 |
| 200 random regions removed | 0.222 | 0.240 |

**Supplementary Table 10**: **Paired comparisons of classifiers given the whole brain or part of it.** Positive differences in accuracy favour the classifier named first. Each comparison uses the same unseen publications and gives the self-forms equal weight. Intervals resample publications and hold the fitted models fixed. P values are FDR-corrected over the eight saved comparisons within each coordinate analysis. The choice of restricted regions and the removal of regions used the training set only. An interval that includes zero leaves the difference uncertain and does not show that two classifiers are equivalent.

| Comparison | Coordinates | Δ accuracy | 95% interval | $q$ |
|---|---|---|---|---|
| Whole brain minus best single region | All | 0.100 | [0.074, 0.126] | 0.0002 |
| Whole brain minus best division | All | 0.064 | [0.038, 0.090] | 0.0002 |
| Whole brain minus 50 nearby regions | All | 0.074 | [0.045, 0.104] | 0.0002 |
| Whole brain minus characteristics and indicators | All | -0.191 | [-0.230, -0.151] | 0.0002 |
| Most predictive minus random removal of 50 | All | -0.007 | [-0.019, 0.005] | 0.3263 |
| Most predictive minus random removal of 100 | All | -0.001 | [-0.015, 0.012] | 0.8648 |
| Most predictive minus random removal of 200 | All | -0.002 | [-0.017, 0.013] | 0.8648 |
| Whole brain added to characteristics and indicators | All | -0.059 | [-0.092, -0.027] | 0.0013 |
| Whole brain minus best single region | Known | 0.105 | [0.079, 0.133] | 0.0002 |
| Whole brain minus best division | Known | 0.098 | [0.070, 0.125] | 0.0002 |
| Whole brain minus 50 nearby regions | Known | 0.080 | [0.048, 0.111] | 0.0002 |
| Whole brain minus characteristics and indicators | Known | -0.183 | [-0.224, -0.141] | 0.0002 |
| Most predictive minus random removal of 50 | Known | 0.004 | [-0.007, 0.016] | 0.5273 |

| Comparison | Coordinates | Δ accuracy | 95% interval | $q$ |
|---|---|---|---|---|
| Most predictive minus random removal of 100 | Known | -0.002 | [-0.017, 0.012] | 0.7384 |
| Most predictive minus random removal of 200 | Known | -0.010 | [-0.026, 0.007] | 0.3399 |
| Whole brain added to characteristics and indicators | Known | -0.025 | [-0.057, 0.007] | 0.2291 |

**Supplementary Table 11**: **What differences in text within a self-form add to prediction.** Each comparison adds the differences in text among publications of the same self-form to a comparison model that already contains the category and the reporting characteristics, with or without the task and clinical indicators. The subtraction of category means and all fitting used the training set. Positive differences favour adding text. The four Brier comparisons, from two comparison models in two coordinate analyses, were corrected as one family, and the comparisons of log loss are a separate family that need not agree. Intervals hold the fitted predictions fixed. Gains and intervals are in units of $10^{-3}$.

| Coordinates | Score | Comparison model | Gain | 95% interval | $q$ |
|---|---|---|---|---|---|
| All | Brier | Characteristics and ten categories | 0.114 | [0.049, 0.181] | 0.0008 |
| All | Brier | Characteristics, indicators and ten categories | 0.056 | [0.042, 0.070] | 0.0002 |
| Known | Brier | Characteristics and ten categories | 0.120 | [0.062, 0.181] | 0.0004 |
| Known | Brier | Characteristics, indicators and ten categories | 0.046 | [0.030, 0.064] | 0.0002 |
| All | Log loss | Characteristics and ten categories | -0.718 | [-1.269, -0.187] | 0.0420 |
| All | Log loss | Characteristics, indicators and ten categories | 0.093 | [-0.089, 0.268] | 0.5076 |
| Known | Log loss | Characteristics and ten categories | -0.201 | [-0.711, 0.302] | 0.5076 |
| Known | Log loss | Characteristics, indicators and ten categories | 0.064 | [-0.122, 0.256] | 0.5076 |

**Supplementary Table 12**: **Prediction within self-forms after filtering anatomical and result statements.** The original text and the filtered abstracts are compared on the same 867 eligible publications, with the same comparison model fitted to all training publications. Positive gains favour the prediction named first. Scores and differences weight the self-forms equally. Intervals hold the fitted models fixed. Tests were Holm-corrected over the three contrasts for each score. The filter reduces statements of findings and names of brain regions, and some may remain. Losses, gains and intervals are in units of $10^{-3}$.

| Contrast | Score | Loss of the model named second | Loss of the model named first | Gain [95% interval] | $p_{\text{Holm}}$ |
|---|---|---|---|---|---|
| Original text versus comparison model | Brier | 86.936 | 86.848 | 0.088 [0.032, 0.147] | 0.0082 |
| Cleaned text versus comparison model | Brier | 86.936 | 86.879 | 0.057 [0.037, 0.078] | 0.0003 |
| Original text versus cleaned text | Brier | 86.879 | 86.848 | 0.031 [-0.014, 0.077] | 0.2069 |
| Original text versus comparison model | Log loss | 306.665 | 307.217 | -0.553 [-1.032, -0.081] | 0.0520 |

| Contrast | Score | Loss of the model named second | Loss of the model named first | Gain [95% interval] | $p_{\text{Holm}}$ |
|---|---|---|---|---|---|
| Cleaned text versus comparison model | Log loss | 306.665 | 306.596 | 0.068 [-0.168, 0.295] | 0.5611 |
| Original text versus cleaned text | Log loss | 306.596 | 307.217 | -0.621 [-1.017, -0.232] | 0.0072 |

**Supplementary Table 13**: **Positions of self-forms on the gradients, and matches with general-imaging gradients.** The first test gives the proportion of the variance in the positions of publications that membership of a self-form explains, with FDR correction across the ten gradients within each coordinate analysis. The second searches 25 general-imaging gradients for the strongest absolute correlation with each self gradient and tests that search against the two kinds of spatial surrogate. The two tests are separate and need not single out the same gradients. Ten gradients were tested for convenience. With known coordinates, the seventh and eighth gradients were exchanged for display after matching. Matches with general-imaging gradients use the original numbering.

| Self gradient | $\eta^2$ | $q_{\text{self-form}}$ | General gradient | $r$ | $q_{\text{BrainSMASH}}$ | $q_{\text{Moran}}$ |
|---|---|---|---|---|---|---|
| **All coordinates** | | | | | | |
| G1 | 0.213 | 0.0002 | G1 | 0.796 | 0.0025 | 0.0020 |
| G2 | 0.078 | 0.0002 | G3 | 0.818 | 0.0025 | 0.0020 |
| G3 | 0.057 | 0.0002 | G2 | 0.802 | 0.0025 | 0.0020 |
| G4 | 0.019 | 0.0616 | G4 | 0.677 | 0.0025 | 0.0020 |
| G5 | 0.038 | 0.0010 | G5 | 0.567 | 0.0080 | 0.0171 |
| G6 | 0.044 | 0.0002 | G6 | 0.562 | 0.0117 | 0.0020 |
| G7 | 0.019 | 0.0540 | G6 | 0.511 | 0.0314 | 0.0117 |
| G8 | 0.025 | 0.0241 | G11 | 0.426 | 0.1575 | 0.0187 |
| G9 | 0.047 | 0.0002 | G14 | 0.375 | 0.3267 | 0.0900 |
| G10 | 0.014 | 0.1634 | G14 | 0.342 | 0.3680 | 0.0644 |
| **Known coordinates** | | | | | | |
| G1 | 0.220 | 0.0003 | G1 | 0.853 | 0.0033 | 0.0025 |
| G2 | 0.099 | 0.0003 | G2 | 0.762 | 0.0033 | 0.0025 |
| G3 | 0.069 | 0.0003 | G3 | 0.759 | 0.0033 | 0.0025 |
| G4 | 0.015 | 0.2936 | G4 | 0.608 | 0.0150 | 0.0271 |
| G5 | 0.053 | 0.0003 | G5 | 0.534 | 0.0383 | 0.0511 |
| G6 | 0.034 | 0.0020 | G6 | 0.538 | 0.0383 | 0.0025 |
| G7 | 0.023 | 0.0439 | G6 | 0.481 | 0.0843 | 0.0271 |
| G8 | 0.027 | 0.0884 | G5 | 0.430 | 0.1125 | 0.0060 |
| G9 | 0.047 | 0.0006 | G13 | 0.373 | 0.2670 | 0.1420 |
| G10 | 0.010 | 0.5484 | G13 | 0.376 | 0.2670 | 0.0375 |

**Supplementary Table 14**: **The graded model compared with the inventory model.** Positive differences mean lower loss for the graded model. The number of patterns in the inventory model is fixed or chosen within the training set, as indicated. Results are given separately for fragments and for whole parcels, and for each coordinate analysis. P values for Brier loss are FDR-corrected over five comparisons within each version and coordinate analysis, and those for log loss are uncorrected secondary results. Intervals resample publications and hold the fitted models fixed. Fragments and whole parcels are different sets of targets, so losses cannot be compared between them. Gains and intervals are in units of $10^{-3}$.

| Version | Coordinates | Inventory model | $\Delta B$ | 95% interval | $q_B$ | $\Delta L$ | 95% interval | $p_L$ |
|---|---|---|---|---|---|---|---|---|
| Fragments | All | One pattern | 3.426 | [2.942, 3.911] | 0.0001 | 13.044 | [11.517, 14.570] | 0.0001 |
| Fragments | All | Two patterns | 2.679 | [2.271, 3.082] | 0.0001 | 9.903 | [8.661, 11.139] | 0.0001 |
| Fragments | All | Three patterns | 2.316 | [1.973, 2.663] | 0.0001 | 8.607 | [7.543, 9.694] | 0.0001 |
| Fragments | All | Ten patterns | 1.874 | [1.552, 2.193] | 0.0001 | 7.057 | [6.052, 8.058] | 0.0001 |
| Fragments | All | Chosen number (1–20) | 1.998 | [1.666, 2.334] | 0.0001 | 7.473 | [6.436, 8.514] | 0.0001 |
| Fragments | Known | One pattern | 3.440 | [2.964, 3.951] | 0.0001 | 12.891 | [11.388, 14.467] | 0.0001 |
| Fragments | Known | Two patterns | 2.729 | [2.343, 3.140] | 0.0001 | 9.959 | [8.765, 11.193] | 0.0001 |
| Fragments | Known | Three patterns | 2.265 | [1.934, 2.619] | 0.0001 | 8.270 | [7.225, 9.380] | 0.0001 |
| Fragments | Known | Ten patterns | 1.893 | [1.573, 2.218] | 0.0001 | 7.163 | [6.153, 8.206] | 0.0001 |
| Fragments | Known | Chosen number (1–20) | 1.916 | [1.593, 2.246] | 0.0001 | 7.261 | [6.255, 8.313] | 0.0001 |
| Whole parcels | All | One pattern | 3.768 | [3.209, 4.323] | 0.0001 | 13.746 | [12.043, 15.423] | 0.0001 |
| Whole parcels | All | Two patterns | 2.903 | [2.422, 3.384] | 0.0001 | 10.356 | [8.913, 11.789] | 0.0001 |
| Whole parcels | All | Three patterns | 2.444 | [2.030, 2.874] | 0.0001 | 8.635 | [7.418, 9.878] | 0.0001 |
| Whole parcels | All | Ten patterns | 1.955 | [1.599, 2.318] | 0.0001 | 7.054 | [5.982, 8.143] | 0.0001 |
| Whole parcels | All | Chosen number (1–20) | 1.882 | [1.523, 2.245] | 0.0001 | 6.972 | [5.876, 8.077] | 0.0001 |
| Whole parcels | Known | One pattern | 3.939 | [3.356, 4.555] | 0.0001 | 14.136 | [12.368, 15.993] | 0.0001 |
| Whole parcels | Known | Two patterns | 3.133 | [2.639, 3.651] | 0.0001 | 10.928 | [9.477, 12.442] | 0.0001 |
| Whole parcels | Known | Three patterns | 2.477 | [2.054, 2.918] | 0.0001 | 8.613 | [7.367, 9.937] | 0.0001 |
| Whole parcels | Known | Ten patterns | 2.062 | [1.663, 2.478] | 0.0001 | 7.383 | [6.166, 8.633] | 0.0001 |
| Whole parcels | Known | Chosen number (1–20) | 2.090 | [1.681, 2.517] | 0.0001 | 7.506 | [6.280, 8.780] | 0.0001 |

**Supplementary Table 15**: **Scores of the reporting characteristics alone, the inventory model and the graded model.** Lower loss means better prediction of the regions reported in the concealed hemisphere. Within each version, the models share folds, targets and reporting characteristics. The scores of the two directions are weighted by their numbers of targets and then averaged over publications. Fragments and whole parcels are different sets of targets, so a difference in loss between them says nothing about which is better. Supplementary Table 14 gives the paired comparisons.

| Version | Coordinates | Model | Brier loss | Log loss |
|---|---|---|---|---|
| Fragments | All | Reporting characteristics alone | 0.081620 | 0.285893 |
| Fragments | All | One pattern | 0.081620 | 0.285893 |
| Fragments | All | Two patterns | 0.080874 | 0.282752 |
| Fragments | All | Three patterns | 0.080511 | 0.281456 |
| Fragments | All | Ten patterns | 0.080068 | 0.279906 |

| Version | Coordinates | Model | Brier loss | Log loss |
|---|---|---|---|---|
| Fragments | All | Chosen number (1–20) | 0.080192 | 0.280322 |
| Fragments | All | Graded model | 0.078194 | 0.272849 |
| Whole parcels | All | Reporting characteristics alone | 0.086784 | 0.300195 |
| Whole parcels | All | Inventory, chosen number | 0.084898 | 0.293421 |
| Whole parcels | All | Graded model | 0.083016 | 0.286450 |
| Fragments | Known | Reporting characteristics alone | 0.083054 | 0.293639 |
| Fragments | Known | One pattern | 0.083054 | 0.293639 |
| Fragments | Known | Two patterns | 0.082342 | 0.290707 |
| Fragments | Known | Three patterns | 0.081879 | 0.289017 |
| Fragments | Known | Ten patterns | 0.081507 | 0.287911 |
| Fragments | Known | Chosen number (1–20) | 0.081529 | 0.288009 |
| Fragments | Known | Graded model | 0.079613 | 0.280748 |
| Whole parcels | Known | Reporting characteristics alone | 0.088501 | 0.309343 |
| Whole parcels | Known | Inventory, chosen number | 0.086652 | 0.302713 |
| Whole parcels | Known | Graded model | 0.084562 | 0.295207 |

**Supplementary Table 16**: **The graded model compared with the on/off model.** Positive differences favour the graded model. The on/off model chooses its number of components and its penalty within the training set. A further on/off model is fixed at ten components, which allows all 1,024 combinations, and chooses only its penalty. Within each version, fragments or whole parcels, Holm correction covers the comparison with the chosen on/off model in both coordinate analyses and on both scores. The comparison with ten components is descriptive and adds no test. Intervals resample publications and hold the fitted models fixed. Losses, gains and intervals are in units of $10^{-3}$.

| Coordinates | Score | Graded | On/off | Difference | 95% interval | $p_{\text{Holm}}$ |
|---|---|---|---|---|---|---|
| On/off model with chosen components on fragments | | | | | | |
| All | Brier | 78.194 | 80.300 | 2.105 | [1.806, 2.410] | 0.0004 |
| All | Log loss | 272.849 | 280.551 | 7.702 | [6.736, 8.698] | 0.0004 |
| Known | Brier | 79.613 | 81.797 | 2.183 | [1.869, 2.510] | 0.0004 |
| Known | Log loss | 280.748 | 288.723 | 7.975 | [6.960, 9.014] | 0.0004 |
| On/off model fixed at ten components on fragments | | | | | | |
| All | Brier | 78.194 | 80.598 | 2.404 | [2.099, 2.706] | – |
| All | Log loss | 272.849 | 281.547 | 8.698 | [7.642, 9.753] | – |
| Known | Brier | 79.613 | 82.186 | 2.573 | [2.245, 2.915] | – |
| Known | Log loss | 280.748 | 289.802 | 9.054 | [7.888, 10.251] | – |
| On/off model with chosen components on whole parcels | | | | | | |
| All | Brier | 83.016 | 85.507 | 2.491 | [2.118, 2.876] | 0.0004 |
| All | Log loss | 286.450 | 295.131 | 8.681 | [7.527, 9.858] | 0.0004 |
| Known | Brier | 84.562 | 87.151 | 2.589 | [2.185, 3.006] | 0.0004 |
| Known | Log loss | 295.207 | 303.771 | 8.563 | [7.319, 9.827] | 0.0004 |

**Supplementary Table 17**: **Where the advantage of the graded model over the inventory model lies.** The difference in Brier loss is separated into gains in predicting the mean probability of reporting and gains in predicting the pattern after each mean is subtracted. The two signed parts sum exactly to the whole before rounding, so one part can exceed the whole when the other is negative. Positive values favour the graded model. Intervals resample publications and hold the fitted models fixed, and no test was made of either part. Main Figure 8d shows the same separation for the comparison with the on/off model. Gains and intervals are in units of $10^{-3}$.

| Coordinates | Set | Total | Pattern | 95% interval | Amount reported | 95% interval |
|---|---|---|---|---|---|---|
| All | All publications | 1.998 | 2.008 | [1.680, 2.350] | -0.010 | [-0.083, 0.065] |
| All | Single analysis and both hemispheres | 1.513 | 1.503 | [1.120, 1.916] | 0.010 | [-0.053, 0.080] |
| Known | All publications | 1.916 | 1.909 | [1.593, 2.233] | 0.007 | [-0.064, 0.085] |
| Known | Single analysis and both hemispheres | 1.461 | 1.446 | [1.052, 1.889] | 0.015 | [-0.045, 0.079] |

**Supplementary Table 18**: **The graded model compared with the inventory model in restricted sets of publications.** The existing predictions are scored for publications with a single extracted analysis, for publications reporting in both hemispheres, and for publications with both properties. Differences are the Brier loss of the inventory model minus that of the graded model on the 987 fragments, so positive values favour the graded model. No model was refitted. The sets address the pooling of several analyses within a publication and publications whose concealed hemisphere is empty. A single extracted analysis is a property of the extraction and not of the psychology studied. Intervals describe the saved predictions in each set. Gains and intervals are in units of $10^{-3}$.

| Coordinates | Set | $n$ | $\Delta B$ | 95% interval |
|---|---|---|---|---|
| All | Single extracted analysis | 374 | 1.278 | [0.924, 1.666] |
| All | Reports in both hemispheres | 855 | 2.155 | [1.804, 2.510] |
| All | Single analysis and both hemispheres | 328 | 1.513 | [1.115, 1.945] |
| Known | Single extracted analysis | 309 | 1.219 | [0.850, 1.619] |
| Known | Reports in both hemispheres | 737 | 2.066 | [1.728, 2.418] |
| Known | Single analysis and both hemispheres | 270 | 1.461 | [1.048, 1.921] |

**Supplementary Table 19**: **What the observed hemisphere contributes to prediction.** For each of the inventory and graded models, predictions using the observed hemisphere are compared with predictions from the same fitted model using only the reporting characteristics. These latter predictions average over the patterns or strengths expected before observing the hemisphere's regional pattern. Both use the same reporting characteristics. Positive differences favour using the observed hemisphere. The comparison is within each model, and its separation into pattern and amount reported need not match that of the comparisons between models. Scores for restricted sets reuse the same fits, and intervals hold the fitted predictions fixed. Gains and intervals are in units of $10^{-3}$.

| Coordinates | Set | $n$ | Model | $\Delta B$ | 95% interval | Pattern | 95% interval | Amount reported | 95% interval |
|---|---|---|---|---|---|---|---|---|---|
| All | All publications | 910 | Graded | 3.430 | [2.947, 3.930] | 3.411 | [2.927, 3.903] | 0.019 | [-0.057, 0.097] |
| All | All publications | 910 | Inventory | 1.432 | [1.197, 1.676] | 1.412 | [1.184, 1.645] | 0.020 | [-0.020, 0.061] |
| All | Single extracted analysis | 374 | Graded | 2.177 | [1.650, 2.759] | 2.158 | [1.641, 2.729] | 0.019 | [-0.041, 0.081] |
| All | Single extracted analysis | 374 | Inventory | 0.897 | [0.652, 1.158] | 0.870 | [0.629, 1.125] | 0.027 | [-0.003, 0.057] |
| All | Reports in both hemispheres | 855 | Graded | 3.684 | [3.177, 4.183] | 3.658 | [3.158, 4.157] | 0.025 | [-0.056, 0.107] |

| Coordinates | Set | $n$ | Model | $\Delta B$ | 95% interval | Pattern | 95% interval | Amount reported | 95% interval |
|---|---|---|---|---|---|---|---|---|---|
| All | Reports in both hemispheres | 855 | Inventory | 1.530 | [1.281, 1.780] | 1.511 | [1.269, 1.751] | 0.019 | [-0.025, 0.063] |
| Known | All publications | 784 | Graded | 3.452 | [2.959, 3.949] | 3.361 | [2.876, 3.855] | 0.090 | [0.011, 0.177] |
| Known | All publications | 784 | Inventory | 1.528 | [1.272, 1.787] | 1.451 | [1.205, 1.702] | 0.077 | [0.035, 0.119] |
| Known | Single extracted analysis | 309 | Graded | 2.129 | [1.566, 2.731] | 2.106 | [1.556, 2.698] | 0.023 | [-0.040, 0.090] |
| Known | Single extracted analysis | 309 | Inventory | 0.910 | [0.650, 1.184] | 0.878 | [0.624, 1.144] | 0.032 | [0.002, 0.063] |
| Known | Reports in both hemispheres | 737 | Graded | 3.704 | [3.195, 4.232] | 3.603 | [3.109, 4.120] | 0.101 | [0.016, 0.193] |
| Known | Reports in both hemispheres | 737 | Inventory | 1.630 | [1.358, 1.908] | 1.552 | [1.289, 1.819] | 0.078 | [0.035, 0.121] |

**Supplementary Table 20**: **The three models among publications that explicitly mention self.** The sets are the publications with the word self in the title or abstract and those with it in the title. Categories and fitted models are unchanged. Positive gains favour the graded model. Holm correction covers the two sets, the two comparisons and the two scores. The sets restrict scoring by a word in the text, within the same corpus. Losses, gains and intervals are in units of $10^{-3}$.

| Compared with | Score | Graded | Other model | Gain [95% interval] | $p_{\text{Holm}}$ |
|---|---|---|---|---|---|
| Title or abstract (720 publications) | | | | | |
| Inventory | Brier | 75.819 | 77.563 | 1.744 [1.396, 2.089] | 0.0008 |
| On/off | Brier | 75.819 | 77.706 | 1.888 [1.575, 2.204] | 0.0008 |
| Inventory | Log loss | 266.496 | 273.314 | 6.818 [5.707, 7.946] | 0.0008 |
| On/off | Log loss | 266.496 | 273.668 | 7.172 [6.118, 8.242] | 0.0008 |
| Title (417 publications) | | | | | |
| Inventory | Brier | 74.280 | 75.649 | 1.369 [0.961, 1.785] | 0.0008 |
| On/off | Brier | 74.280 | 75.825 | 1.546 [1.165, 1.942] | 0.0008 |
| Inventory | Log loss | 262.778 | 268.418 | 5.640 [4.333, 7.016] | 0.0008 |
| On/off | Log loss | 262.778 | 269.546 | 6.768 [5.423, 8.151] | 0.0008 |

**Supplementary Table 21**: **The advantage of the graded model in each self-form.** Positive gains favour the graded model over the inventory model or the on/off model, as indicated. Estimates reuse the predictions of the principal analysis. One summary gives each self-form one tenth of the weight, and the principal summary weights publications equally. Intervals come from bootstrap samples within self-forms, hold the fitted models fixed and are descriptive. Some intervals include zero although every estimate is positive. Gains and intervals are in units of $10^{-3}$.

| Self-form | $N$ | Inventory minus graded | On/off minus graded |
|---|---|---|---|
| **Brier loss** | | | |
| Autobiographical memory | 146 | 3.013 [2.005, 4.118] | 2.745 [1.908, 3.652] |
| Negative self-reference | 139 | 1.160 [0.572, 1.820] | 1.560 [0.963, 2.202] |
| Ownership and agency | 135 | 2.663 [1.691, 3.681] | 2.321 [1.441, 3.238] |
| Self-appraisal | 124 | 1.610 [0.650, 2.829] | 2.121 [1.222, 3.156] |
| Trait evaluation | 87 | 1.620 [0.852, 2.423] | 1.691 [1.018, 2.370] |

| Self-form | $N$ | Inventory minus graded | On/off minus graded |
|---|---|---|---|
| Interoception | 79 | 1.583 [0.718, 2.505] | 2.129 [1.173, 3.152] |
| Monitoring and insight | 58 | 2.692 [1.665, 3.868] | 2.623 [1.554, 3.873] |
| Self–other distinction | 52 | 2.480 [1.208, 3.947] | 2.359 [1.377, 3.437] |
| Face/voice recognition | 46 | 0.493 [-0.616, 1.683] | 1.058 [-0.069, 2.260] |
| Body image/identity | 44 | 1.907 [0.717, 3.276] | 1.888 [0.809, 3.087] |
| All publications | 910 | 1.998 [1.668, 2.333] | 2.105 [1.813, 2.414] |
| Self-forms weighted equally | 910 | 1.922 [1.598, 2.270] | 2.050 [1.748, 2.359] |
| **Marginal log loss** | | | |
| Autobiographical memory | 146 | 9.597 [6.690, 12.725] | 8.409 [5.913, 11.123] |
| Negative self-reference | 139 | 4.801 [2.774, 7.040] | 6.095 [4.085, 8.157] |
| Ownership and agency | 135 | 10.793 [7.633, 14.050] | 9.075 [6.262, 11.968] |
| Self-appraisal | 124 | 6.323 [3.567, 9.542] | 7.852 [4.979, 10.976] |
| Trait evaluation | 87 | 6.214 [3.514, 8.968] | 7.362 [4.478, 10.408] |
| Interoception | 79 | 5.894 [3.060, 8.757] | 7.885 [4.682, 11.325] |
| Monitoring and insight | 58 | 9.774 [6.291, 13.770] | 9.220 [5.602, 13.215] |
| Self–other distinction | 52 | 9.455 [5.372, 14.024] | 8.235 [4.744, 11.848] |
| Face/voice recognition | 46 | 2.147 [-1.996, 6.511] | 4.756 [0.820, 8.776] |
| Body image/identity | 44 | 7.434 [3.363, 11.723] | 6.594 [2.621, 10.912] |
| All publications | 910 | 7.473 [6.430, 8.500] | 7.702 [6.742, 8.691] |
| Self-forms weighted equally | 910 | 7.243 [6.192, 8.367] | 7.548 [6.526, 8.586] |

**Supplementary Table 22**: **Filtered-abstract categories and prediction.** All fits use the same 867 eligible publications and the original outer and inner fold memberships, restricted before training. Groups are fixed corpus annotations. Learned ten- and three-group comparisons use principal axes and k-means fitted to training publications only. New-group weighting averages over the 10 filtered-abstract categories, and original-group weighting averages over the original ten. Gains are reductions in Brier loss, and positive gains favour the first named model. Gain and interval units are $\times 10^{-3}$. Relative gains divide the gain by the equally weighted reference loss. Benjamini–Hochberg correction covers seven model comparisons or three pooling comparisons separately within weighting. Intervals condition on fitted models.

| Weighting | Comparison | Gain | 95% interval | Gain (%) | $q$ |
|---|---|---|---|---|---|
| New groups | Reporting + filtered groups vs Reporting characteristics | 0.451 | [0.329, 0.576] | 0.502 | 0.00014 |
| New groups | Indicators + filtered groups vs Reporting and task/clinical indicators | 0.166 | [0.107, 0.227] | 0.185 | 0.00014 |
| New groups | Reporting + original groups vs Reporting characteristics | 0.503 | [0.368, 0.642] | 0.559 | 0.00014 |
| New groups | Indicators + original groups vs Reporting and task/clinical indicators | 0.149 | [0.086, 0.213] | 0.165 | 0.00014 |
| New groups | Ten learned filtered-text groups vs Three learned filtered-text groups | 0.058 | [0.008, 0.108] | 0.065 | 0.0273 |
| New groups | Ten learned full-text groups vs Three learned full-text groups | 0.133 | [0.079, 0.186] | 0.147 | 0.0002333 |
| New groups | Reporting characteristics vs Intercept only | 6.253 | [5.333, 7.241] | 6.504 | 0.00014 |
| New groups | Filtered categories vs All-publication pool | 0.776 | [0.431, 1.120] | 0.807 | 0.0001 |
| New groups | Filtered categories vs Size-matched pool | 1.570 | [1.224, 1.916] | 1.620 | 0.0001 |
| New groups | Filtered categories vs Other-category pool | 1.177 | [0.787, 1.568] | 1.219 | 0.0001 |
| Original groups | Reporting + filtered groups vs Reporting characteristics | 0.405 | [0.289, 0.521] | 0.463 | 0.00014 |
| Original groups | Indicators + filtered groups vs Reporting and task/clinical indicators | 0.145 | [0.089, 0.202] | 0.166 | 0.00014 |
| Original groups | Reporting + original groups vs Reporting characteristics | 0.447 | [0.323, 0.575] | 0.511 | 0.00014 |
| Original groups | Indicators + original groups vs Reporting and task/clinical indicators | 0.141 | [0.079, 0.205] | 0.160 | 0.00014 |
| Original groups | Ten learned filtered-text groups vs Three learned filtered-text groups | 0.045 | [-0.001, 0.094] | 0.052 | 0.0726 |
| Original groups | Ten learned full-text groups vs Three learned full-text groups | 0.108 | [0.054, 0.161] | 0.123 | 0.0004667 |
| Original groups | Reporting characteristics vs Intercept only | 5.491 | [4.709, 6.302] | 5.910 | 0.00014 |
| Original groups | Filtered categories vs All-publication pool | 0.533 | [0.269, 0.795] | 0.574 | 0.0001 |
| Original groups | Filtered categories vs Size-matched pool | 1.297 | [1.031, 1.561] | 1.385 | 0.0001 |

| Weighting | Comparison | Gain | 95% interval | Gain (%) | $q$ |
|---|---|---|---|---|---|
| Original groups | Filtered categories vs Other-category pool | 0.879 | [0.576, 1.182] | 0.943 | 0.0001 |

**Supplementary Table 23**: **Paired log-loss comparisons and a Brier reference scale.** These use the saved original predictions for 910 publications with all coordinates and 784 with known spaces. Losses are averaged across repeats within publication before inference with the original categories weighted equally. Positive gains favour the first model. Benjamini–Hochberg correction covers the same seven model comparisons as the original Brier family, separately per policy. Asterisks identify unadjusted paired probabilities for the separate reporting-versus-intercept Brier reference. Gain and interval units are $\times 10^{-3}$. Intervals condition on fitted models.

| Policy | Score | Comparison | Gain | 95% interval | $q$ or $p^*$ |
|---|---|---|---|---|---|
| All | Log loss | Original categories and indicators vs Reporting and task/clinical indicators | -0.018 | [-0.642, 0.589] | 0.9547 |
| All | Log loss | Original categories and indicators vs Reporting characteristics | -7.608 | [-9.891, -5.340] | 0.000175 |
| All | Log loss | Reporting and original categories vs Reporting characteristics | 4.725 | [3.781, 5.747] | 0.000175 |
| All | Log loss | Ten learned groups vs Three learned groups | 0.475 | [-0.124, 1.054] | 0.1205 |
| All | Log loss | Continuous text vs Reporting and task/clinical indicators | 9.933 | [8.030, 11.903] | 0.000175 |
| All | Log loss | Continuous text vs Original categories and indicators | 9.951 | [7.977, 11.962] | 0.000175 |
| All | Log loss | Rank-ten continuous text vs Rank-three continuous text | -1.690 | [-3.167, -0.193] | 0.0371 |
| All | Brier | Reporting characteristics vs intercept only | 5.449 | [4.697, 6.240] | 0.0001* |
| Known | Log loss | Original categories and indicators vs Reporting and task/clinical indicators | 0.039 | [-0.554, 0.622] | 0.8998 |
| Known | Log loss | Original categories and indicators vs Reporting characteristics | -9.786 | [-12.380, -7.240] | 0.00014 |
| Known | Log loss | Reporting and original categories vs Reporting characteristics | 4.347 | [3.430, 5.324] | 0.00014 |
| Known | Log loss | Ten learned groups vs Three learned groups | 0.880 | [0.264, 1.503] | 0.0049 |
| Known | Log loss | Continuous text vs Reporting and task/clinical indicators | 10.560 | [8.254, 12.945] | 0.00014 |
| Known | Log loss | Continuous text vs Original categories and indicators | 10.521 | [8.181, 12.886] | 0.00014 |
| Known | Log loss | Rank-ten continuous text vs Rank-three continuous text | -7.069 | [-8.315, -5.850] | 0.00014 |
| Known | Brier | Reporting characteristics vs intercept only | 5.410 | [4.561, 6.287] | 0.0001* |

**Supplementary Table 24**: **Boundary records in the original semantic partition.** Counts refer to the 915 original text records. Negative silhouettes compare mean cosine distances to all own-category versus nearest-other-category records. Nonpositive centre margins compare distances to category centres and define a separate boundary set. Stability is the fraction of 27 grid solutions in which a record stays with the majority of its fixed category peers. The last two columns separate negative-silhouette records from the remaining records. These are descriptive fixed-partition diagnostics.

| Category | $n$ | Negative | Negative (%) | Margin $\leq 0$ | Stability negative | Stability other |
|---|---|---|---|---|---|---|
| Autobiographical memory | 146 | 31 | 21.2 | 26 | 0.474 | 0.796 |
| Negative self-reference | 141 | 26 | 18.4 | 15 | 0.664 | 0.853 |
| Ownership and agency | 136 | 28 | 20.6 | 15 | 0.753 | 0.837 |
| Self-appraisal | 125 | 50 | 40.0 | 13 | 0.654 | 0.672 |
| Trait evaluation | 87 | 14 | 16.1 | 11 | 0.780 | 0.827 |
| Interoception | 80 | 10 | 12.5 | 5 | 0.596 | 0.920 |
| Monitoring and insight | 58 | 3 | 5.2 | 4 | 0.938 | 0.998 |
| Self–other distinction | 52 | 2 | 3.8 | 4 | 1.000 | 0.869 |
| Face and voice recognition | 46 | 1 | 2.2 | 1 | 1.000 | 0.963 |
| Body image and identity | 44 | 3 | 6.8 | 6 | 0.963 | 0.992 |

**Supplementary Table 25**: **Original categories crossed with filtered-abstract categories.** Both partitions are compared on the same 867 eligible publications. Columns are anonymous categories regrouped from filtered abstracts, ordered by decreasing size, with letters used only as identifiers. The group count follows the amended tie rule described in Supplementary Note 1.

| Original category | A | B | C | D | E | F | G | H | I | J |
|---|---|---|---|---|---|---|---|---|---|---|
| Autobiographical memory | 6 | 11 | 1 | 31 | 9 | 77 | 1 | 1 | 1 | 0 |
| Negative self-reference | 110 | 7 | 0 | 7 | 1 | 0 | 0 | 2 | 1 | 5 |
| Ownership and agency | 0 | 0 | 113 | 2 | 5 | 1 | 3 | 1 | 3 | 3 |
| Self-appraisal | 1 | 22 | 1 | 30 | 21 | 2 | 1 | 0 | 0 | 38 |
| Trait evaluation | 3 | 71 | 0 | 3 | 3 | 1 | 1 | 0 | 0 | 1 |
| Interoception | 17 | 0 | 1 | 3 | 0 | 0 | 0 | 52 | 1 | 0 |
| Monitoring and insight | 0 | 1 | 1 | 4 | 0 | 0 | 0 | 0 | 47 | 0 |
| Self–other distinction | 2 | 2 | 0 | 1 | 44 | 0 | 0 | 0 | 0 | 1 |
| Face and voice recognition | 1 | 0 | 1 | 14 | 3 | 0 | 26 | 0 | 1 | 0 |
| Body image and identity | 0 | 8 | 1 | 0 | 0 | 0 | 34 | 0 | 0 | 0 |

**Supplementary Table 26**: **Anatomical overlap after regrouping filtered abstracts.** Original ten-group maps are shown for the full corpus and matched eligible cohort. The selected filtered-abstract grouping uses the middle tied counts and text stability. The initial smallest-on-ties grouping is retained for transparency. Known excludes coordinates in unknown or other spaces before fitting, leaving 749 matched publications. Counts exclude publication maps with no modelled-activation support. All fits use the broad coordinate policy, fixed assumed sample size 20, the same 2mm mask, and Benjamini–Hochberg correction at 0.05 across group-by-voxel tests within each partition and coordinate analysis. The all-group count is the intersection of significant maps, and maximal sharing is the largest number of maps containing any one voxel. Dice averages over all pairs. No cluster-size cutoff is imposed on the intersections, and these counts do not establish a biologically meaningful or self-specific core.

| Grouping | Policy | $n$ | Groups | All-group voxels | Maximal sharing | Mean Dice |
|---|---|---|---|---|---|---|
| Original full corpus | All | 911 | 10 | 0 | 8 | 0.151 |
| Original full corpus | Known | 785 | 10 | 0 | 8 | 0.136 |
| Original groups, matched cohort | All | 867 | 10 | 0 | 8 | 0.137 |
| Original groups, matched cohort | Known | 749 | 10 | 0 | 8 | 0.126 |
| Filtered, central rule | All | 867 | 10 | 3 | 10 | 0.172 |
| Filtered, central rule | Known | 749 | 10 | 0 | 9 | 0.159 |
| Filtered, initial rule | All | 867 | 8 | 6 | 8 | 0.183 |
| Filtered, initial rule | Known | 749 | 8 | 1 | 8 | 0.166 |

**Supplementary Table 27**: **Anatomical overlap with family-wise error correction.** The original full-corpus categories and the categories regrouped from filtered abstracts were evaluated with the same NiMARE correction, fixed assumed sample size 20 and the broad coordinate policy. Each category used 1,000 random-coordinate datasets. Clusters were formed at uncorrected $p < 0.001$ using six face neighbours. Cluster size, cluster mass and voxelwise correction are all reported. The per-map level 0.05 controls FWE within each map, and 0.005 additionally corrects across ten maps by Bonferroni. All is the all-coordinate analysis, which treats coordinates in unknown or other spaces as MNI, and Known excludes them. The filtered-abstract fits retain 867 and 749 publications, respectively. The original reference uses its full corpus, so it is not a matched-cohort FWE comparison. All-ten voxels counts the intersection of ten corrected maps. Maximal sharing counts the largest number of corrected maps containing any voxel. No minimum extent is applied to their intersection. The additional FWE check of the filtered-abstract categories was specified after their three-voxel FDR intersection was known.

| Categories | Coordinates | Correction | Per-map $\alpha$ | All-ten voxels | Maximal sharing |
|---|---|---|---|---|---|
| Original, full corpus | All | Cluster size | 0.05 | 0 | 7 |
| Original, full corpus | All | Cluster size | 0.005 | 0 | 6 |
| Original, full corpus | All | Cluster mass | 0.05 | 0 | 7 |
| Original, full corpus | All | Cluster mass | 0.005 | 0 | 7 |
| Original, full corpus | All | Voxel | 0.05 | 0 | 5 |
| Original, full corpus | All | Voxel | 0.005 | 0 | 5 |
| Filtered abstracts | All | Cluster size | 0.05 | 0 | 8 |
| Filtered abstracts | All | Cluster size | 0.005 | 0 | 7 |
| Filtered abstracts | All | Cluster mass | 0.05 | 0 | 7 |
| Filtered abstracts | All | Cluster mass | 0.005 | 0 | 7 |
| Filtered abstracts | All | Voxel | 0.05 | 0 | 5 |
| Filtered abstracts | All | Voxel | 0.005 | 0 | 5 |
| Filtered abstracts | Known | Cluster size | 0.05 | 0 | 7 |
| Filtered abstracts | Known | Cluster size | 0.005 | 0 | 7 |
| Filtered abstracts | Known | Cluster mass | 0.05 | 0 | 7 |
| Filtered abstracts | Known | Cluster mass | 0.005 | 0 | 7 |
| Filtered abstracts | Known | Voxel | 0.05 | 0 | 5 |
| Filtered abstracts | Known | Voxel | 0.005 | 0 | 4 |

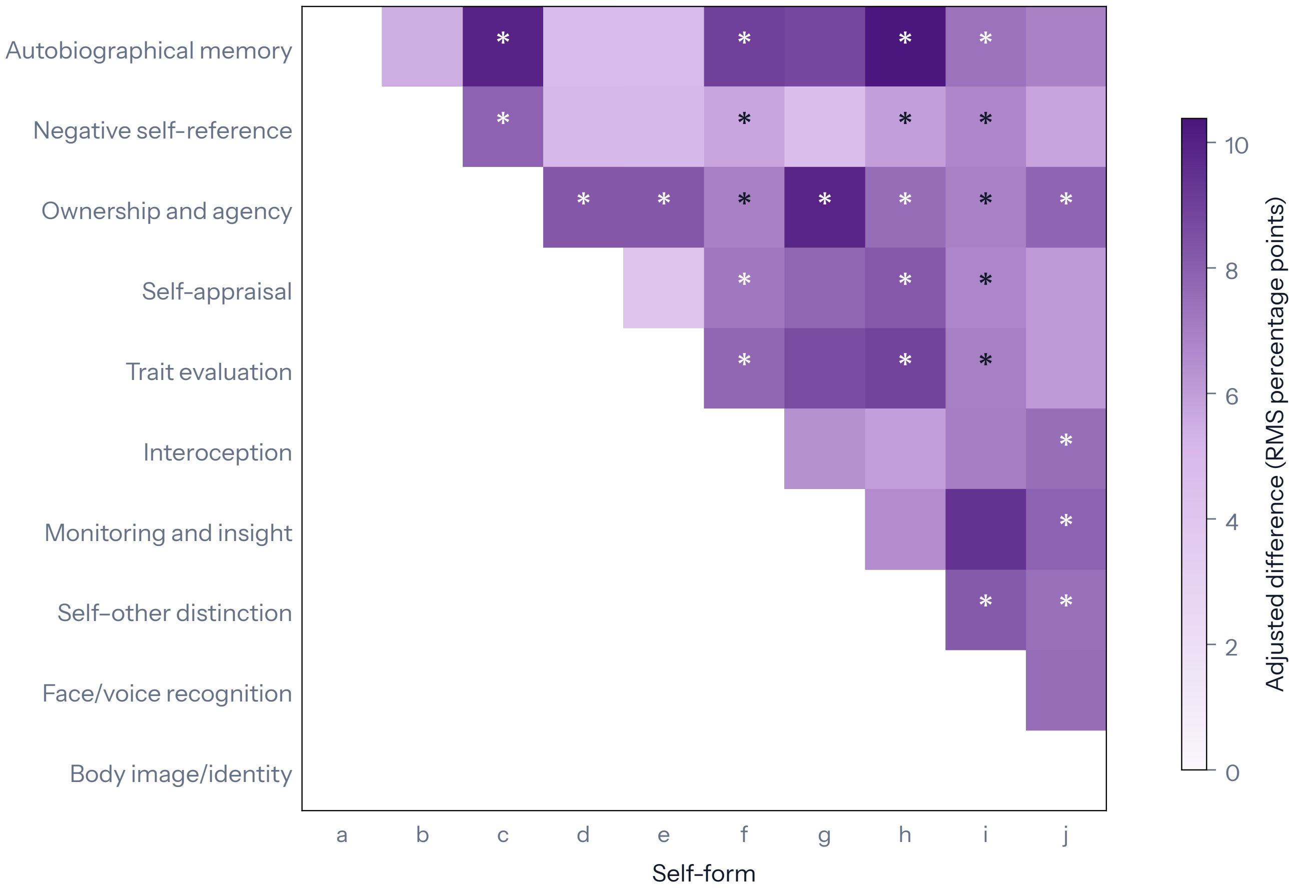

**Supplementary Fig. 1 Differences in regional profiles between pairs of self-forms.** Colour shows the root mean square of the differences in the probability of reporting each region, in percentage points, after adjustment for groups of similar publications. Larger values mean that two whole profiles differ more. Stars mark tests of whole profiles with Holm-adjusted $p < 0.05$ across the 45 pairs, from 9,999 shuffles of labels within groups of similar reporting and clinical content. Results are shown with all coordinates. Letters for self-forms follow main Fig. 2, and Supplementary Table 4 reports both coordinate analyses.

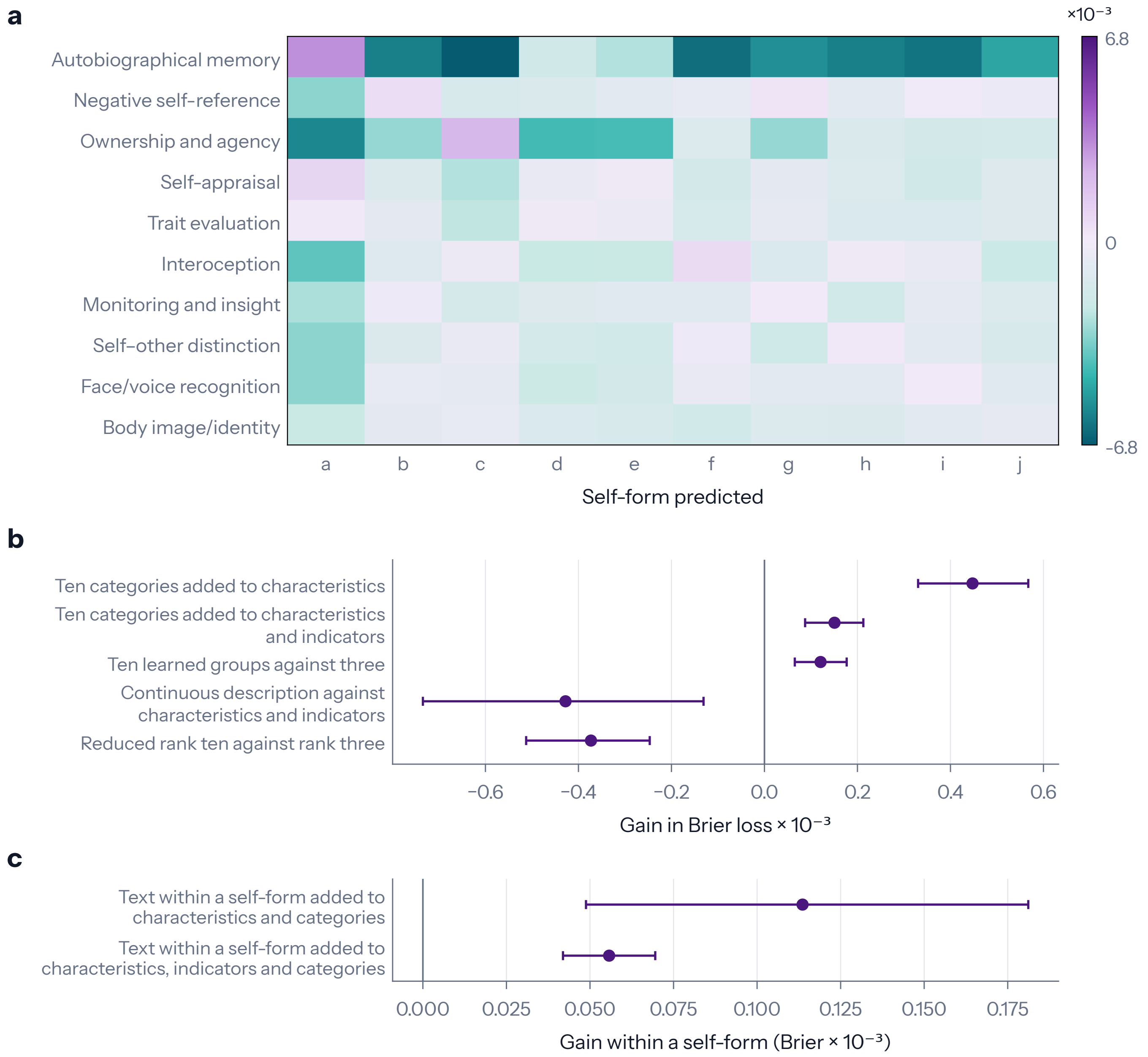

**Supplementary Fig. 2 What a study describes predicts the regions it reports, between and within self-forms.** **a**, Gain in Brier loss when the pattern of the self-form in each row, estimated from training publications, predicts unseen publications of the self-form in each column, relative to the pooled pattern for that column. **b**, Gains from the original categories over reporting characteristics, with or without task and clinical indicators. Further comparisons show gains from ten learned groups over three, from a continuous description of text over the indicators, and from rank ten over rank three in the reduced-rank description. Rank gives the number of patterns retained when predicting regional differences from text. Positive values favour the model named first. **c**, Gains from differences in text within a self-form, over comparison models that already contain the category. Gains in b and c are in units of $10^{-3}$, with 95% bootstrap intervals over publications that hold the fitted models fixed. Supplementary Tables 5, 6, 7 and 11 give the numbers.

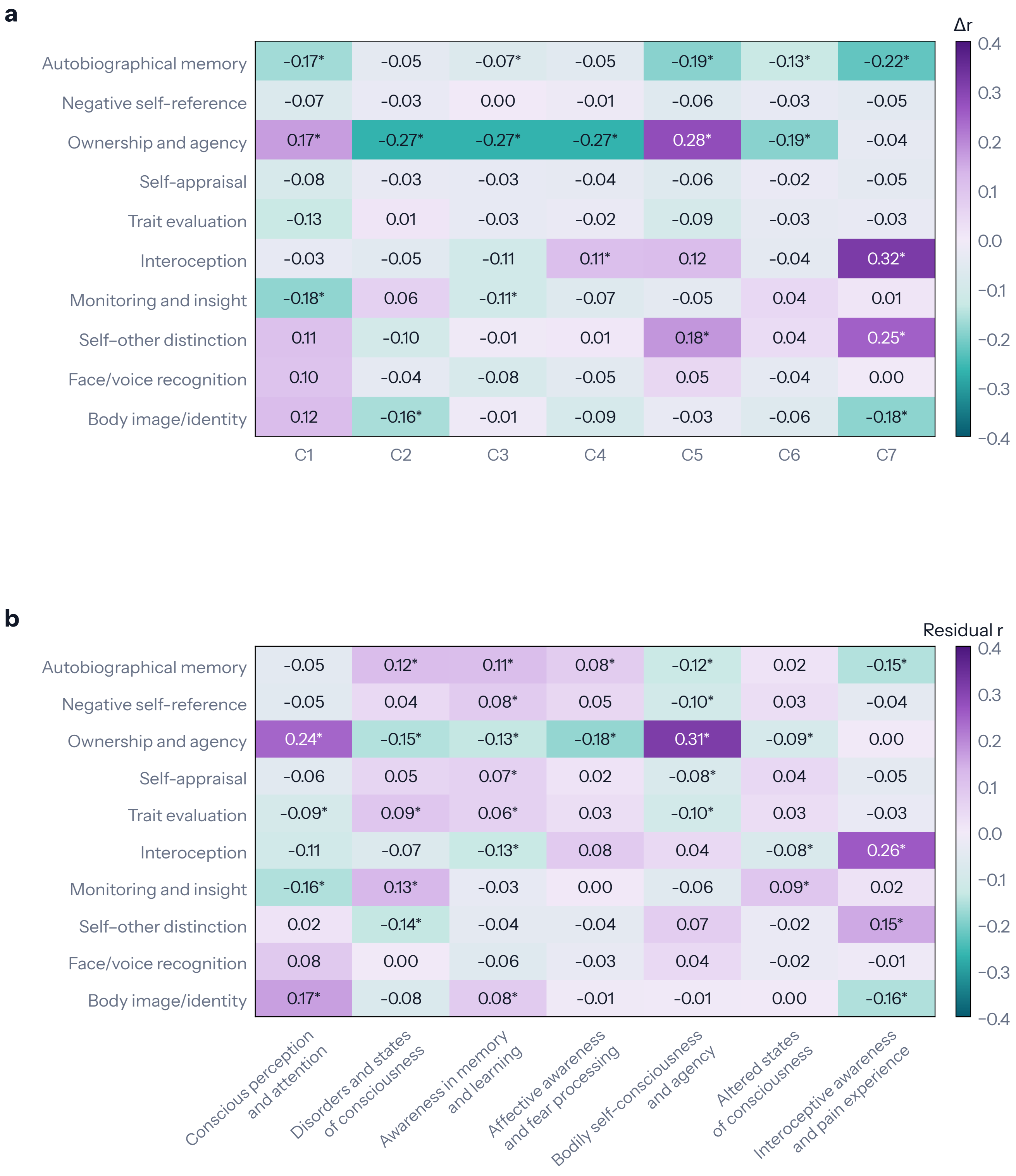

**Supplementary Fig. 3 Particular pairings of self-forms with groupings of consciousness research stand out. a**, Correlations between maps of how often regions were reported for a self-form and a consciousness grouping, after subtracting the mean under shuffled labels. **b**, The same correlations after subtracting the mean for each self-form and each consciousness grouping and restoring the overall mean. In panel a, negative values mark correlations below the shuffled mean. In panel b, they mark correlations below the value predicted by the means for the self-form and consciousness grouping. Stars mark tests of each displayed value against its own distribution under 9,999 shuffles of labels within groups of similar publications, with FDR correction over the 70 cells of each panel. Values in panel b are shown as they are, without subtraction of their shuffled means. C1–C7 follow main Fig. 6. Results are shown with all coordinates.

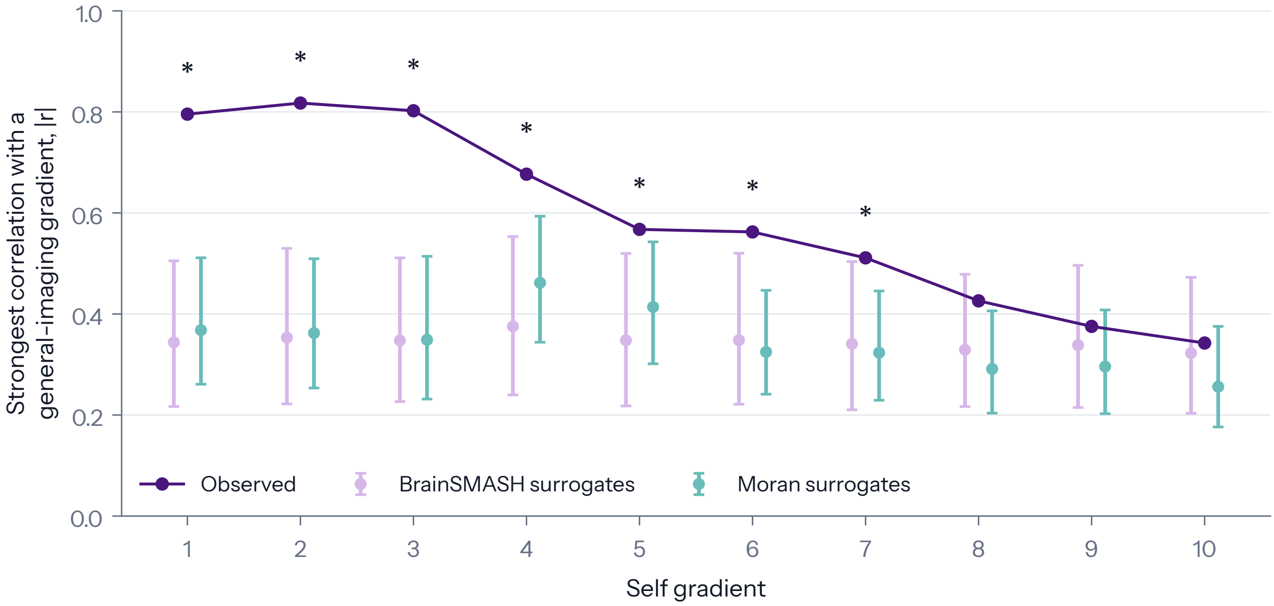

**Supplementary Fig. 4 Gradients of the self literature match gradients of imaging research in general.** Purple points show the strongest absolute correlation of each self gradient with any of 25 general-imaging gradients. Pale points (BrainSMASH) and teal points (Moran spectral randomisation) show the mean of that statistic for the two kinds of spatial surrogate, artificial gradients generated under different models of spatial structure. Whiskers span the central 95% of 999 surrogates, each of which went through the whole search for its best match. Stars mark FDR-adjusted $q < 0.05$ under both tests. Results are shown with all coordinates. Main Figure 7c reports reliability across halves of the corpus, and Supplementary Table 13 compares the two coordinate analyses.

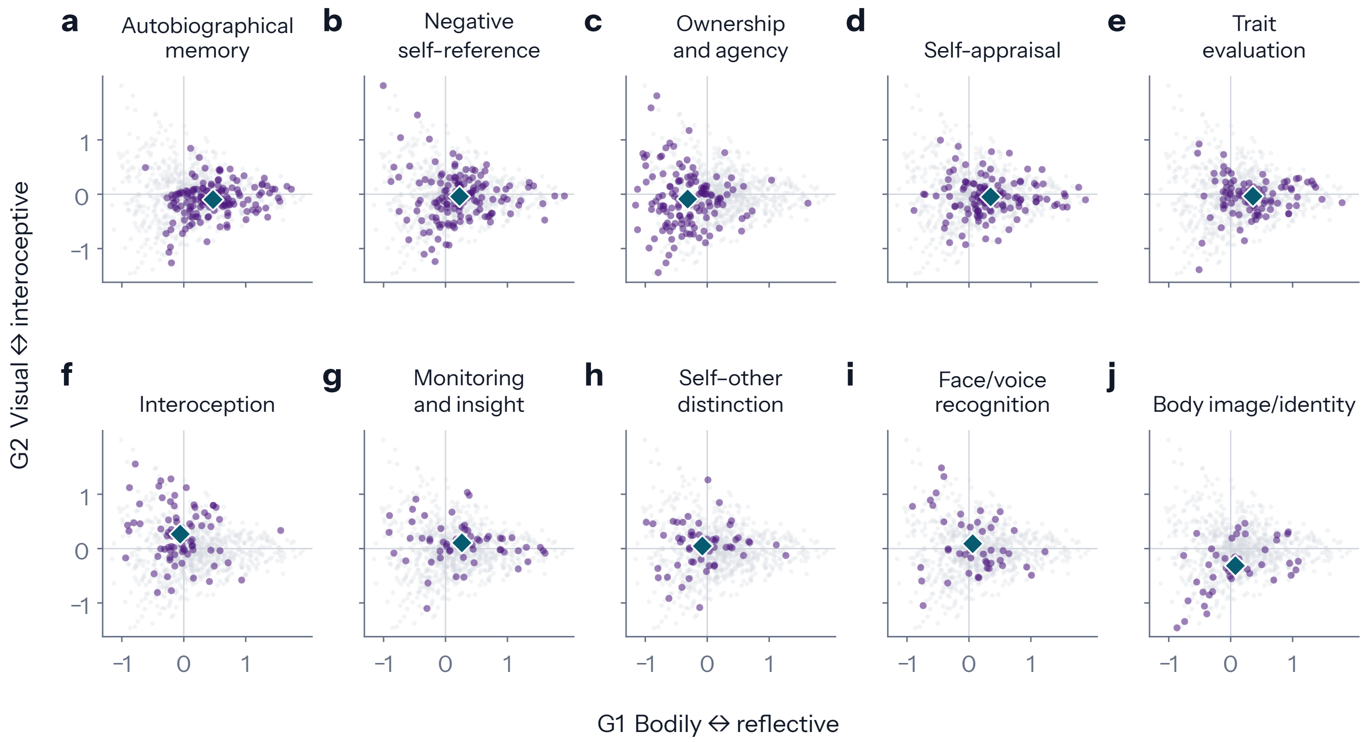

**Supplementary Fig. 5 Self-forms differ in mean position while their publications overlap.** Each point places a publication on the first two gradients at the average gradient value of its reported regions. Panels **a**–**j** show each self-form in purple against all other publications in grey, and teal diamonds mark the mean of the self-form. Panels follow the order of self-forms in main Fig. 2, with abbreviated names and identical axes. Main Figure 7b and Supplementary Table 13 report the tests of position.
